# Definition of alleles and altered regulatory motifs across Cas9-edited cell populations

**DOI:** 10.1101/775361

**Authors:** Kirk T. Ehmsen, Matthew T. Knuesel, Delsy Martinez, Masako Asahina, Haruna Aridomi, Keith R. Yamamoto

## Abstract

**Background:** Genetic alteration of candidate response elements at their native chromosomal loci is the only valid determinant of their potential transcriptional regulatory activities. Targeted DNA cleavage by Cas9 coupled with cellular repair processes can produce arrays of alleles that can be defined by massively parallel sequencing by synthesis (SBS), presenting an opportunity to generate and survey edited cell populations that include informative alterations. Such editing efforts commonly rely on subclonal enrichment to isolate cells with preferred genotypic properties at target loci; short nucleotide adducts (indices/barcodes) allow PCR-amplified molecules from diverse sample sources to be pooled, sequenced, and demultiplexed to resolve source-specific content. Not widely available, however, are capabilities for barcoding thousands of clones, or for automated analysis of individual candidate regulatory loci PCR-amplified and sequenced from a genetically heterogeneous population—specifically, imputation of discrete genotype(s) by allele definition and abundance, and identification of altered regulatory factor binding motifs.

**Results:** We describe a panel of 192 8-nucleotide barcode primers compatible with Illumina® sequencing platforms, and the application of these barcodes to genotypic analysis of Cas9-edited clones. Permutations of the ninety-six i7 (read 1) and ninety-six i5 (read 2) barcodes allow unique labeling of up to 9,216 distinct samples. We created three independent Python scripts: *SampleSheet.py* automates construction of Illumina® Sample Sheets encoding up to 9,216 barcode:sample relationships; *ImputedGenotypes.py* defines alleles and imputes genotypes from demultiplexed fastq files; *CollatedMotifs.py* flags transcription factor recognition motif matches altered in alleles relative to a reference sequence.

**Conclusions:** Code-enabled definition of alleles and regulatory motifs in sequenced, demultiplexed amplicons facilitates evaluation of genetic diversity in up to 9,216 distinct samples. Here, we demonstrate the utility of three scripts in analysis of cell populations targeted by Cas9 for disruption of glucocorticoid receptor (GR) binding sites near *FKBP5*, a GR-regulated gene in the human adenocarcinoma cell line A549. SampleSheet.py, ImputedGenotypes.py, and CollatedMotifs.py operate independently and are broadly applicable beyond the case described here.

## Background

The glucocorticoid receptor (GR) is a transcriptional regulatory factor (TF) that binds to specific sequence motifs at genomic glucocorticoid response elements (GREs) and nucleates combinatorial assembly of multicomponent transcriptional regulatory complexes, which modulate expression of cognate target genes. Three features greatly complicate determination of transcriptional regulatory activity by a genomic GR-occupied region (GOR). First, in any given context, most GORs appear to lack function^1^, outnumbering glucocorticoid-responsive genes by an order of magnitude or more^2–4^. Second, a GRE may comprise multiple GORs scattered over tens or hundreds of kilobases, each contributing distinct regulatory outcomes. Third, GRE activities are highly context-dependent, and must be assessed in their normal chromosomal environments. As a result, very few GREs or other response elements have been functionally validated^5^.

In principle, functional validation could be addressed by Cas9-driven targeted genomic editing of candidate response elements, coupled with regulatory analysis of target gene(s). Edited subclones could be identified from Cas9-treated bulk cell populations, using sequencing by synthesis (SBS)^6, 7^ to assess target regions. However, this would require multiplexing hundreds to thousands of samples and a computational workflow for deconvolution and analysis. Specifically, three unmet needs were apparent:

### 1. An index (barcode) set and Sample Sheet sufficient for SBS discrimination of thousands of subclonal genotypes

In many Illumina® sequencing applications, DNA sequences from distinct samples are labeled with barcodes, pooled (multiplexed) as a library applied to a single flow cell, and demultiplexed based on sample:index relationships specified in a Sample Sheet, which relates user-specified workflow parameters to Illumina® sequencer control software. Illumina® systems formally limit barcode assignments to <96 (single-index) or <384 (dual-indexed/paired-end) distinct samples. SBS analysis of independent samples at larger scale requires an augmented barcode set, and automated sample:barcode assignment in a Sample Sheet that specifies thousands of data relationships.

### 2. Rapid clonal genotype imputation to locate potentially scarce mutant clones with desired characteristics

Excellent web-based and command line interface (CLI) tools, *e.g.*, CRISPR-GA^8^, AGEseq^9^, CRISPResso^10^, CRISPR-DAV^11^, Cas-analyzer^12^, BATCH-GE^13^, BE-Analyzer^14^, and CRIS.py^15^, perform aggregate mutation counts and efficiencies from next-generation sequencing (NGS) data, reporting population-distributed allele type resolution relative to bundled mutation frequencies returned by aggregate analyses (*e.g.*, TIDE ^16^, TIDER^17^). We sought to develop a tool that specifically imputes genotypes for clonal isolates based on computationally defined alleles, and visually maps Cas9 guide RNA sequence(s) on alignments for evaluation of Cas9-associated mutations.

### 3. Automated TF binding site (TFBS) collation, to infer potential consequences of Cas9 edits to TF function in mutagenized response elements

Cas9-edited cell populations typically display a broad spectrum of alleles at targeted loci, generated as insertion/deletion (indel) outcomes of non-homologous DNA double-strand break repair processes^18^. For indels in putative response elements, identifying mutation-associated loss or gain of putative TFBSs would inform interpretations and/or predictions of functional consequences.

To address these needs, we set out to automate sequence processing from input to output of a Cas9-editing effort focused on candidate response elements, seeking to expedite clone selection for retrieval and archiving, to prioritize clones for analysis based on imputed genotypic definitions, and to anticipate potential functional consequences based on altered TF motif matches (TFBS).

## Implementation

We developed three scripts (SampleSheet.py, ImputedGenotypes.py and CollatedMotifs.py; https://github.com/YamamotoLabUCSF, supporting resources at DOI 10.5281/zenodo.3406862) that automate principal steps in *(i)* sample preparation for massively parallel amplicon sequencing, including Sample Sheet creation for Illumina® sequencing by synthesis (SBS) platforms; *(ii)* genotype imputation (allele definition) for up to 9,216 pooled, independently barcoded amplicon samples; and *(iii)* TFBS motif comparison between imputed alleles and a reference (*e.g.*, wild-type) allele. All scripts require Python 3.7 or greater for operation on Mac OSX or Windows systems; scripts are available as annotated Jupyter notebooks^19^ for interactive use in a web browser, as program files (.py) that can be run at a command-line interface (CLI), and pre-compiled in Open Virtualization Format for virtualization (*e.g.*, in Oracle VM VirtualBox, https://www.virtualbox.org/). Text editor and PDF reader software are required to access output files.

### External dependencies

When operated from a CLI, SampleSheet.py suggests but does not require download and installation of a Python package *prettytable*. Whether operated from a CLI or Jupyter notebook, ImputedGenotypes.py and CollatedMotifs.py require the Python packages NumPy, SciPy and psutil, plus download and availability of BLASTN (BLAST+ suite, NCBI^20^); ImputedGenotypes.py additionally requires fdpf and PyPDF2 libraries as well as a BLASTN reference genome database for sequencing alignments; CollatedMotifs.py further requires MAKEBLASTDB (BLAST+ suite), Meme suite^21^ installation (for FIMO and FASTA-GET-MARKOV)^22^, and a FIMO-compatible TFBS motif reference (position frequency matrix) file for transcription factor binding site queries^20^. In Windows OS, Meme suite programs (FIMO and FASTA-GET-MARKOV) require virtualization and CollatedMotifs.py must be run from within a hypervisor (*e.g.*, Oracle VM VirtualBox).

## Results & Discussion

A Cas9 editing effort is typically applied to cell populations, but genotypic characterization of subclones requires clonal isolation, locus-specific PCR amplification, and sequencing. We developed a custom library of 192 Illumina® platform-compatible, uniquely barcoded oligonucleotide primers—96 ‘forward’ and 96 ‘reverse’ as defined by sense/antisense to read1 orientation in the Illumina® MiSeq workflow—as indices assignable to unique clones (**Supp. Fig. 1**). Barcodes at this 96×96 scale exceed commercial barcode availability, increasing the number of independent amplicon sources that can be pooled for paired-end sequencing and demultiplexing of reads to 9,216.

We generated hundreds of independent mutant clones in the human lung adenocarcinoma cell line A549, targeting genomic regions occupied by the glucocorticoid receptor (GR, product of the *NR3C1* gene)^23^, a DNA-binding transcriptional regulatory factor. We developed code to support and expedite a workflow that reports alleles and genotypes for up to 9,216 clones from a single Illumina® SBS run (**Fig. 1**). SampleSheet.py automates preparation of an Illumina® Sample Sheet, the text document that defines well:barcode assignments for demultiplexing on Illumina® sequencing platforms (**Fig. 2**). ImputedGenotypes.py facilitates rapid convergence to genotype from demultiplexed fastq files^24^, simplifying identification of cultured clones of interest to archive and examine (**Fig. 3**). Finally, CollatedMotifs.py summarizes alterations to transcription factor binding site (TFBS) motif matches for each clonal isolate relative to a reference sequence, capitalizing on public repositories of sequence-selective position frequency matrices for characterized DNA-binding regulatory factors (**Fig. 4**). DNA sequence alterations associated with Cas9 editing in putative regulatory elements may cause losses or gains of binding sites for transcription regulatory factors, making their annotation in clones useful for interpretation of potential functional consequences (**Fig. 5**).

**Figure 1.**
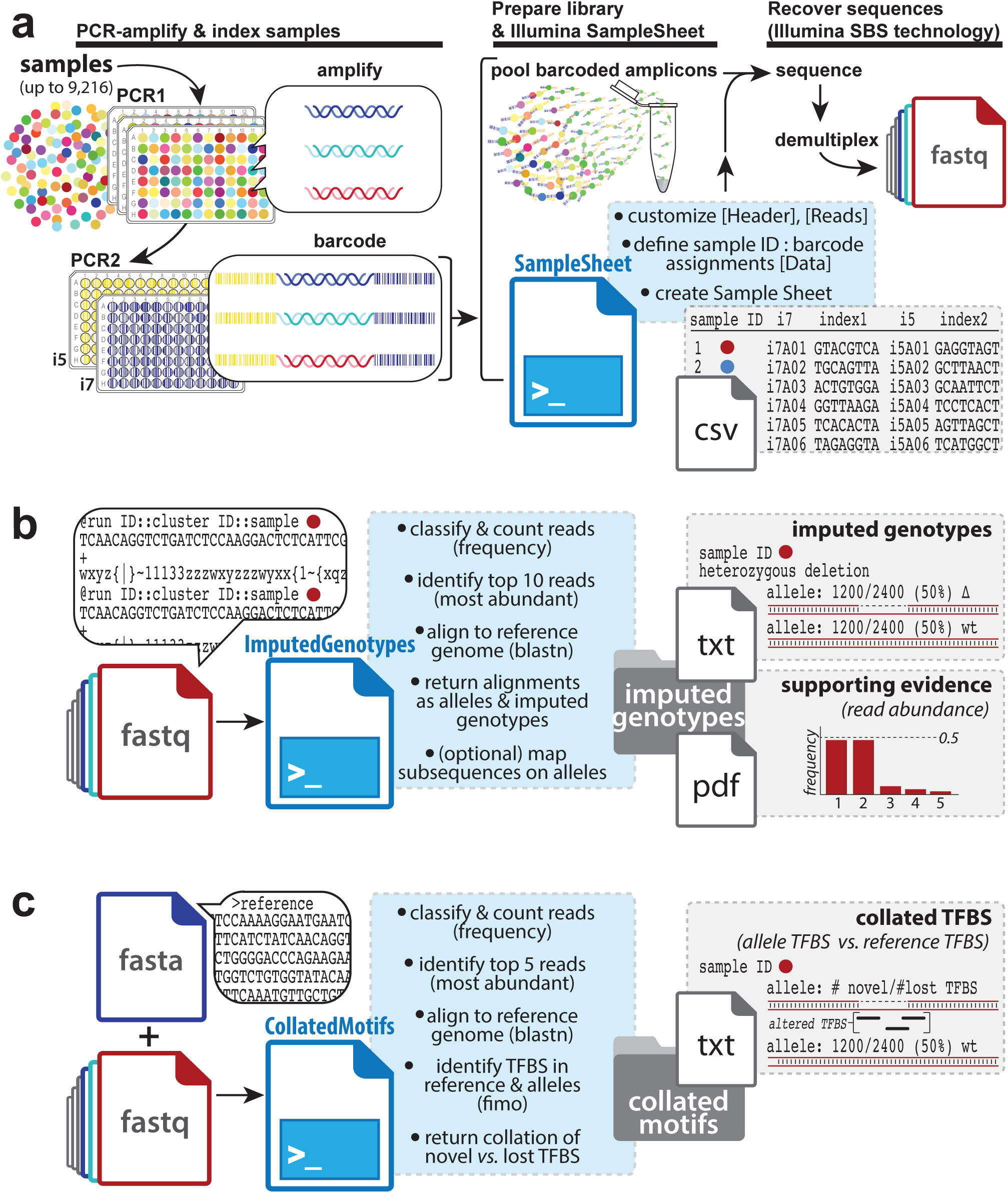
Three Python scripts facilitate analysis of genetic diversity in deeply sequenced amplicons. ***(a)* SampleSheet.py** operates in workflows that require demultiplexing of barcoded sequences, automating construction of an Illumina® Sample Sheet with up to 9,216 sample:barcode relationships defined in its [Data] table. Briefly, target loci from up to 9,216 samples can be amplified and indexed in two consecutive PCRs (PCR1 & PCR2), from essentially any nucleic acid source (*e.g.*, population samples, Cas9-edited clonal isolates [*colored circles*]). After arraying genetic source material from individual samples in 96-well or 384-well (not shown) plates for amplification (PCR1), small amounts of each PCR1 product are used as templates in second reactions (PCR2) primed by pairs of uniquely barcoded forward and reverse primers compatible with Illumina® sequencing platforms (ninety-six i7, ninety-six i5 barcode possibilities). Barcoded amplicons are pooled as a library; user-supplied values at SampleSheet.py prompts are expanded to populate a Sample Sheet with sample:barcode designations, enabling read demultiplexing into up to 9,216 sample-specific fastq files following Illumina® SBS; ***(b)* ImputedGenotypes.py** accepts any number of fastq files as input, applying Python counter functions to classify and count read frequencies. After aligning the most abundant reads to a reference genome (BLASTN), alleles are hypothesized and defined based on relative read frequencies and alignment comparison to the wild-type reference (*e.g.*, SNPs, indels); genotypes are imputed based on allele definitions. Optionally, DNA subsequences (short oligonucleotide sequences) can be mapped onto allele outputs to flag positions and/or presence/absence of specific sequence motifs; ***(c)* CollatedMotifs.py** accepts fastq files as input, along with a single fasta file defining reference sequence(s). Like ImputedGenotypes.py, CollatedMotifs.py identifies candidate alleles by read frequency and alignment to a reference sequence; it then identifies and compares matches to TFBS motifs in reference and allele sequences (Meme FIMO), returning a visualization of novel and lost TFBS in each allele.

**Figure 2.**
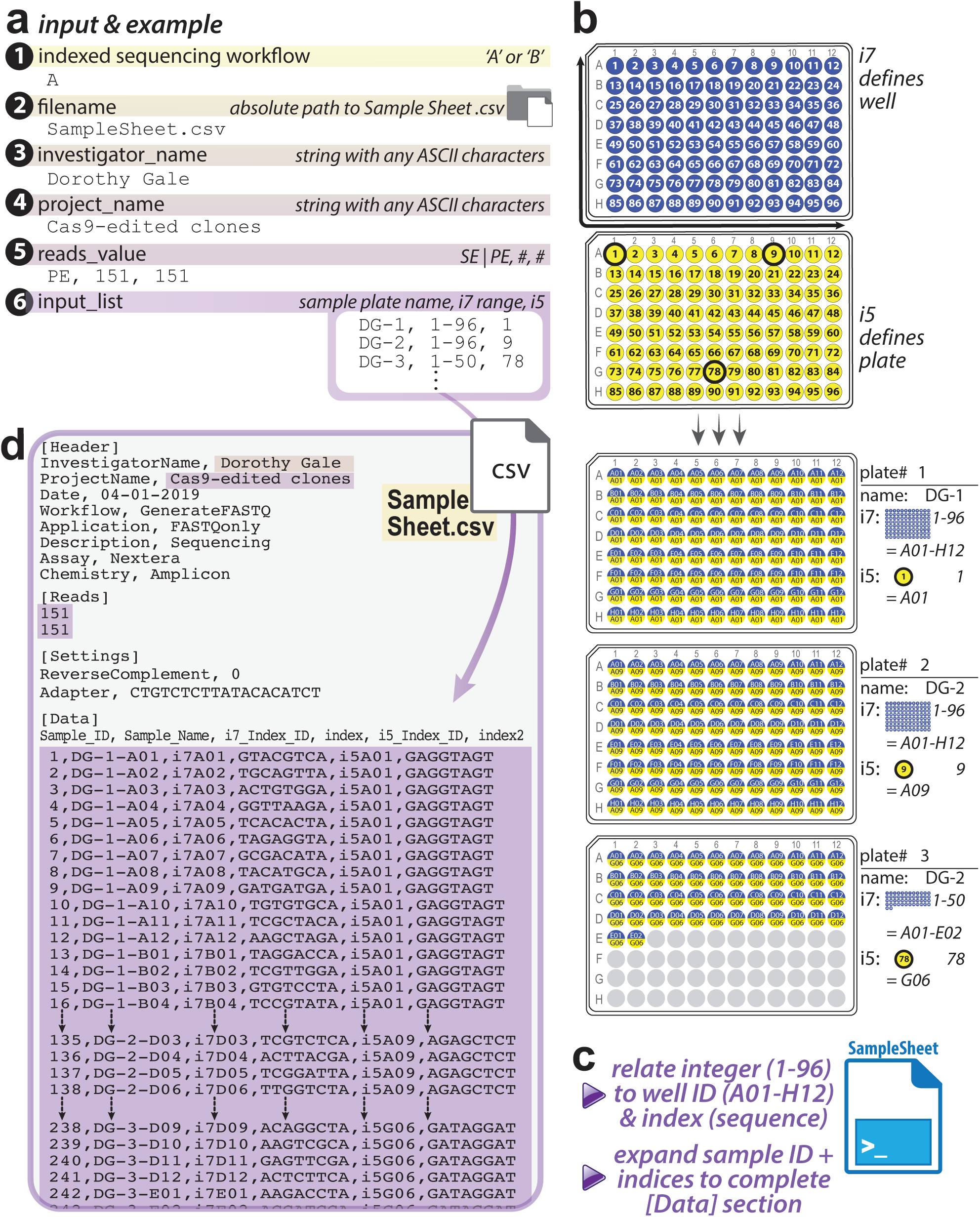
SampleSheet.py: automated Illumina® Sample Sheet construction for sequencing and demultiplexing of up to 9,216 barcoded samples. ***(a)* SampleSheet.py** anticipates user-defined, console-supplied entries for six variables, which define *(1)* the Illumina® Indexed Sequencing Workflow (‘A’ or ‘B’), *(2)* the absolute path at which the Sample Sheet file will be created, plus subsections of an Illumina® Sample Sheet: two **[Header]** values (for the keys *(3)* ‘InvestigatorName’ & *(4)* ‘ProjectName’), the **[Reads]** section (indicating *(5)* the number of sequencing cycles for single-end or paired-end formats), and *(6)* a list of sample plate names and i7+i5 barcode permutations, the principal **[Data]** output; ***(b)*** PCR strategy that links amplicons to barcodes underlies the relationship that SampleSheet.py creates between an individual sample identity (plate and well ID) and its distinctive i7+i5 barcode combination. In this strategy, i7 sequence (blue) defines individual wells of a 96-well plate, and is used across plates; i5 (yellow), in contrast, defines up to all wells of a single plate; ***(c)*** SampleSheet.py delineates relationships between i7 and i5 identities and barcode identities, enabling automated expansion of appropriate sample:index relationships in the Sample Sheet [Data] table, ***(d)*** and their output in the Sample Sheet for up to 9,216 samples. In the example given, samples arrayed in three ninety-six well plates have been uniquely labeled, and an input list with three lines of text is expanded into a list of 242 entries, with index sequences accurately presented in the Sample Sheet.

**Figure 3.**
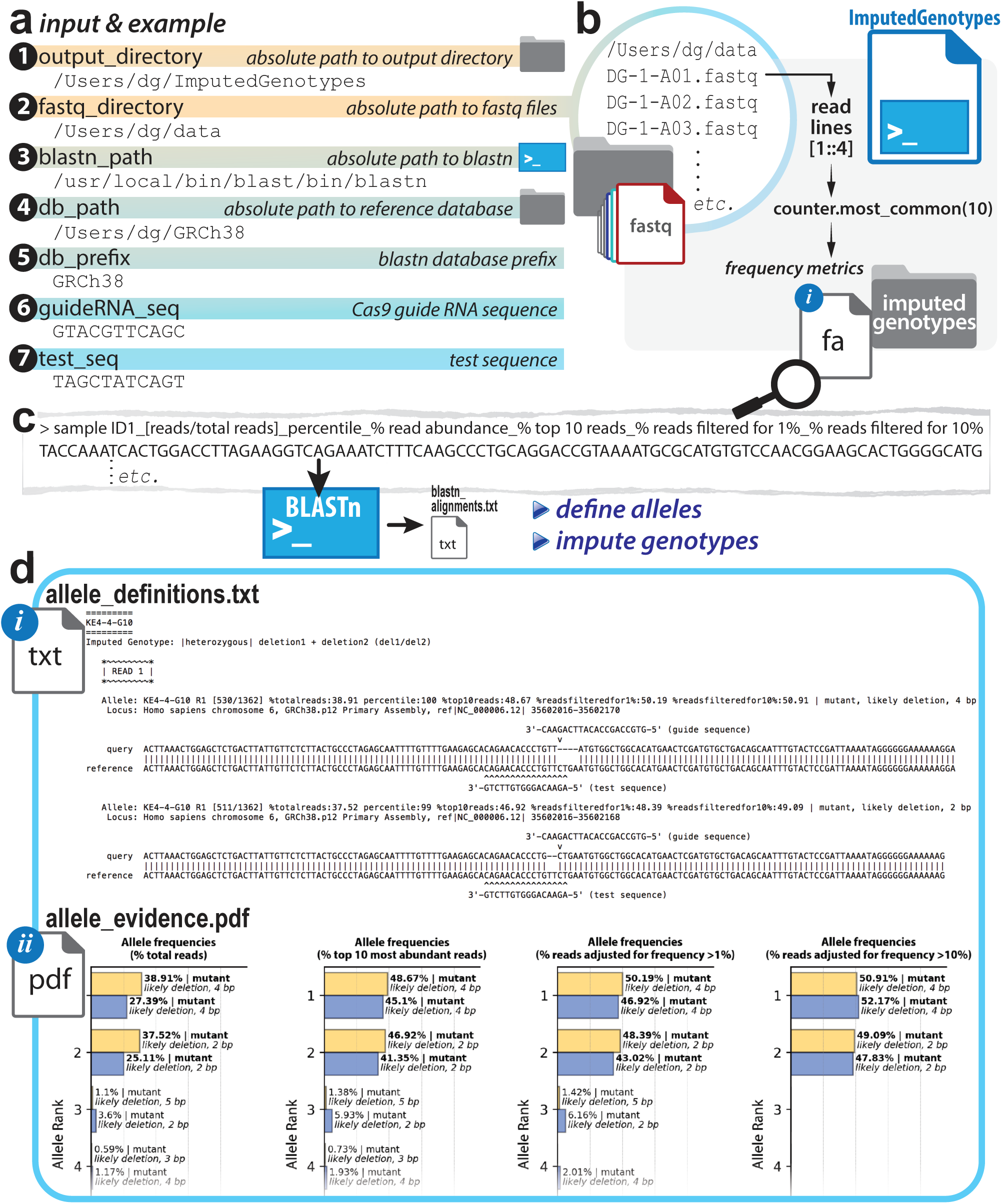
ImputedGenotypes.py: allele definition and genotype imputation based on read abundance in deeply sequences amplicons. ***(a)*** Seven user-defined inputs specify locations of three directories and one executable file (BLASTN), as well as two optional short nucleotide sequences to be mapped onto analyzed sequence outputs—ultimately generating eight output files; ***(b)*** fastq files are read into ImputedGenotypes.py, which classifies reads by relative representation (Python counter function) and defines alleles based on calculated frequency. The top ten most abundant reads are labeled with sample ID and abundance/representation and populated into a fasta file (*fasta.fa*), which is ***(c)*** passed to BLASTN for alignment to a position in the reference genome/sequence provided as a BLASTN database in *(a)* (*input #3*). The output alignment file (*blastn_alignments.txt*) is parsed from html format to populate lists and dictionaries with read-specific metadata (allele identifier, ‘hit’ position in reference sequence, alignment to ‘hit’); these data are the basis of allele type *definition* (deletion, insertion, wild-type, etc.) and subsequent genotype *imputation*; ***(d)*** Eight output files are generated (*allele_definitions.txt, allele_definitions.csv, allele_evidence.pdf* (optional)*, blastn_alignments.txt, fasta.fa, imputed genotypes.txt, population_summary.txt, script_metrics.txt*); portions of the principal output in *allele_definitions.txt* and *allele_evidence.pdf* are shown (see Supplement for further examples).

**Figure 4.**
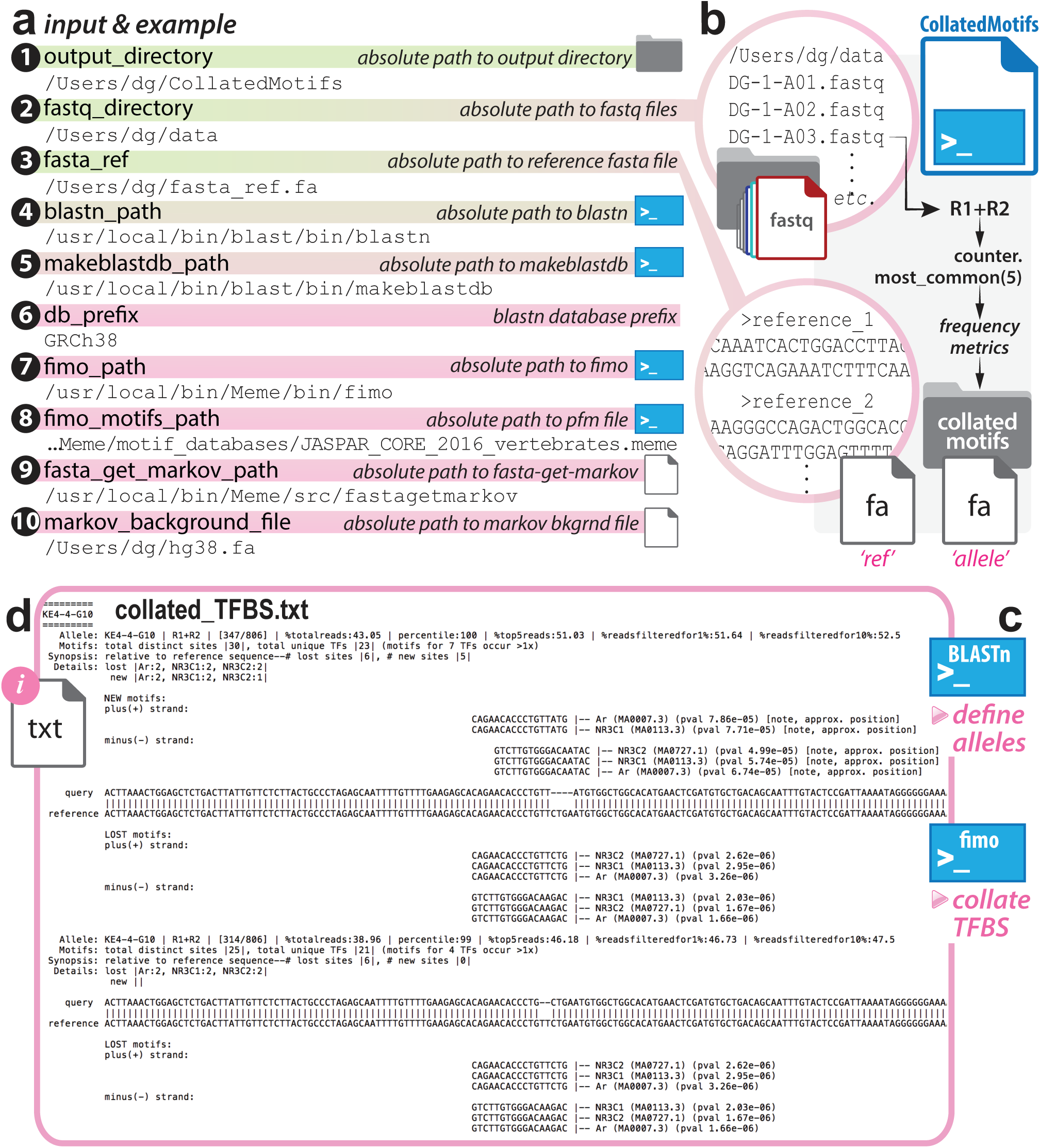
CollatedMotifs.py: identification of altered regulatory motifs in defined alleles, relative to reference sequence. ***(a)*** Ten user-defined inputs specify locations of two directories, four executable files (BLASTN, FIMO, MAKEBLASTDB, FASTA-GET-MARKOV), three files (fasta file with reference sequences, text file with TFBS motifs, text file with sequence(s) from which markov background will be defined), and a single database prefix string; ***(b)*** fastq files are read into CollatedMotfis.py, which merges R1 & R2 sequences (from paired-end sequencing) and identifies the top 5 most abundant read types for each sample; ***(c)*** reference sequences in user-provided fasta file and fasta file containing top 5 reads for each sample are provided to BLASTN for alignments and to FIMO for TFBS determinations, ***(d)*** with alignments and collated TFBS (lost and/or gained) displayed in output file *collated_TFBS.txt*.

**Figure 5.**
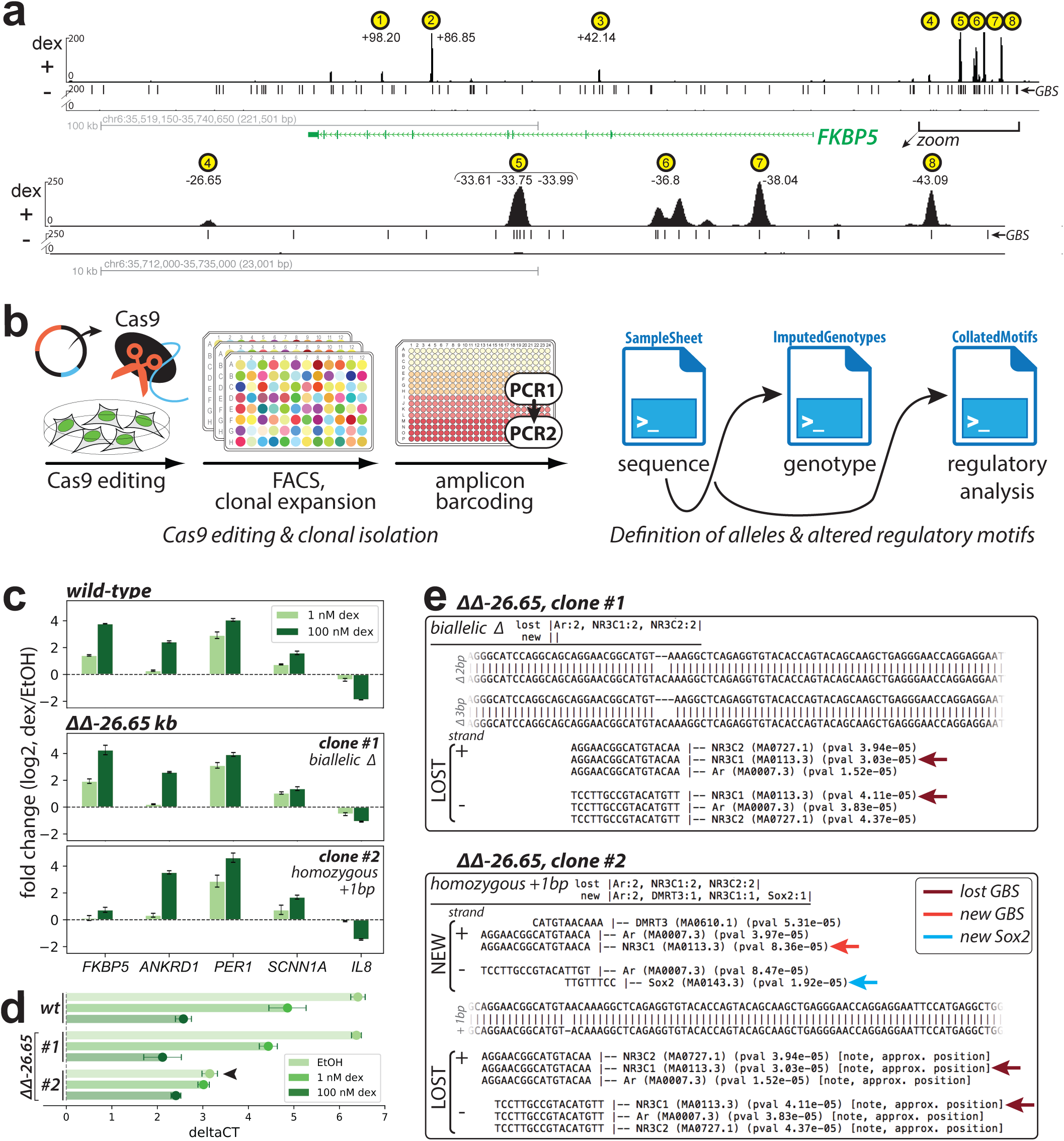
Use case: evaluation of Cas9-altered loci occupied by human glucocorticoid receptor (GR) near a glucocorticoid-regulated gene, FKBP5. ***(a)** top panel*, GR ChIP-seq (A549±100 nM dex, 1.5 h) indicating eight intronic and promoter-proximal dex-dependent GR-occupied regions (GORs (yellow circles)) in vicinity of dex-induced *FKBP5*; GOR coordinates defined as peak summit distance from FANTOM5-defined TSS for *FKBP5* transcript variant 1 (RefSeq NM_004117, coordinates chr6:35,688,937 in GRCh38)^35^; *lower panel*, zoom-in of region comprising GOR4-8; ***(b)*** Cas9-induced mutagenesis, clonal isolation, and amplicon sequencing procedure for genotype imputation and mutant clone identification: Cas9 and sgRNA expressed from transfected episomes; FACS isolation of single cells into wells of 96-well plates; PCR1 amplification of target loci and PCR2 indexing via barcode primers; deep sequencing (MiSeq) supported by SampleSheet.py; genotyping supported by ImputedGenotypes.py; TFBS synopsis *via* CollatedMotifs.py; ***(c)*** regulatory analysis (fold change (log2) of mRNA levels) for five dex-responsive genes (*FKBP5*, *ANKRD1*, *PER1*, *SCNN1A*, *IL8*) sampled from A549 (wild-type and GOR mutants) ±1 nM and 100 nM dex, 4 h (RT-qPCR, ΔΔCT dex relative to ethanol control, n=3, mean±std). Clones #1 and #2 (biallelic Δ and homozygous +1 bp in GBS at GOR4 (-26.65 kb) peak summit) exhibit distinct regulatory consequences for *FKBP5* induction, unique among evaluated genes; ***(d)*** ΔCT analysis (CT *FKBP5* – CT_geometric mean_ for three reference controls (*GAPDH, HBMS, RPL19*)) indicates that loss of *FKBP5* dex induction in ΔΔ-26.65 clone #2 is partly attributable to increased baseline (EtOH) transcript level (arrow) relative to wild-type; ***(e)*** CollatedMotifs.py TFBS annotations in ΔΔ-26.65 clones #1 *vs.* #2 show that both clones lose native GBS (maroon arrow, *lost GBS*), but clone #2 reconstitutes a novel GBS (red arrow, *new GBS*). Furthermore, clone #2 also exhibits TFBS for novel TFs (*e.g.*, blue arrow, *new Sox2*), suggesting an avenue to further examine—and potentially explain—ostensible mutation-specific regulatory functions for response elements under study.

### SampleSheet.py and 96×96 paired-end barcoding of up to 9,216 samples

Illumina® SBS workflows generally entail library construction, cluster generation, sequencing, and data processing. In all cases, a single, plain-text, comma-separated (*.csv) file—the Sample Sheet—mediates communication of user preferences and sequencing workflow specifications to the software in charge of sequencing operations and data acquisition. Fundamental run properties (*e.g.*, sequencing cycle number, chemistry) and key links between library identities and barcodes are documented, allowing reads to be demultiplexed into individual fastq files after sequencing. Sample Sheets can be constructed in Illumina® Experiment Manager (available only for Windows platforms), or in any plain-text editor. Illumina® provides an excellent whitepaper describing Sample Sheet sections and preparation in Pub. No. 970-2017-004-A (2017).

For small numbers of indices (*e.g*., tens), samples and their identifying barcodes are easily compiled manually in a text document. For hundreds to thousands of samples, however, manual compilation becomes time-consuming and prone to error (*e.g.*, typos or mistaken entries that lead to sample:barcode mispairings). SampleSheet.py automates Sample Sheet construction, allowing a Sample Sheet with up to 9,216 data entries to be compiled in seconds from a single list of up to 96 short lines of text (**Fig. 1a, Fig 2**), linking sample IDs in 96-well plates to specified i7 and i5 barcode pairs.

### User inputs and specification of sequencing run properties

SampleSheet.py prompts users for six values, entered as text at individual Jupyter interface or CLI prompts (**Fig. 2a, Table 1, Supp. Fig. 2, Supp. Fig. 3a**). These values include: 1) Illumina® Indexed Sequencing Workflow (A *vs.* B), 2) absolute path to output directory and filename for Sample Sheet, 3) Investigator Name, 4) Project Name, 5) Single-end (SE) or Paired-end (PE) sequencing run with cycle number(s), and 6) list of sample:barcode relationships.

**Table 1.** SampleSheet.py inputs.

| User input | Purpose | Example |
| --- | --- | --- |
| ***Illumina Dual Indexed Sequencing Workflow*** |  |  |
| Workflow A vs. B | Specify whether i5 and i7 indices are reported during PE sequencing as ‘forward’ or ‘reverse complement’ sequences | A |
| ***Absolute path to Sample Sheet file name*** |  |  |
| Output directory & Sample Sheet file name | Define directory in which Sample Sheet file (with designated name) will be created | /Users/name/IlluminaNGS/SampleSheet.csv<br>(MacOS/Linux) or<br>C:\Users\name\IlluminaNGS\SampleSheet.csv (Windows) |
| ***Sample Sheet sections*** |  |  |
| [Header]:<br>Investigator Name, Project Name | Customize investigator name and project name to appear in [Header] section | Dorothy Gale, Cas9-edited clones |
| [Reads]:<br>SE vs. PE, and # of cycles for R1/R2 | Specify cycle number for R1 (SE) or R1 and R2 (PE) | PE, 151, 151 |
| [Data]:<br>Sample:barcode relationships | Specify relationship between sample name common to wells of a 96-well plate, and i7/i5 indices | **CLI:**<br>DG-1, 1-96, 1<br>DG-2, 1-96, 9<br>DG-3, 1-50, 78<br>**Jupyter notebook:**<br>DG-1, 1-96, 1; DG-2, 1-96, 9; DG-3, 1-50, 78 |

#### 1-Barcode sequence orientation in SampleSheet—Workflow ‘A’ vs. ‘B’

Illumina® indexed sequencing uses two different paired-end Indexed Sequencing Workflows, depending on sequencer (**Supp. Fig. 4**). In *Workflow A*, index 2 is sequenced before read 2 resynthesis, meaning that i7 is sequenced as the reverse complement and i5 is sequenced on the forward strand (applicable to NovaSeq 6000, MiSeq, HiSeq 2500, HiSeq 2000); in *Workflow B*, index 2 is sequenced *after* read2 synthesis, creating the reverse complement of both index1 (i7) and index2 (i5) (applicable to iSeq100, miniSeq, NextSeq, HiSeq X, HiSeq 3000, 4000). This distinction requires attention to barcode sequence entry in Sample Sheet fields ‘index1’ and ‘index2’; users of SampleSheet.py must verify the index sequencing workflow on the sequencer on which they will load their libraries, as SampleSheet.py fills barcode sequences based on specified workflow (**Fig. 2a**, **input #1; Supp. Fig. 3a-1**).

#### 2-Absolute path to file output

Users are next prompted for a text string that specifies the absolute path to a location where the Sample Sheet file will be created (**Fig. 2a**, **input #2; Supp. Fig. 3a-2**); this string must be entered as a series of directory name(s) beginning at the file system root (*e.g.*, /Users on Mac, C:\ on Windows operating systems), ending in the file name to be created by the script (*e.g*., SampleSheet.csv). Console prompts specify that regardless of the operating system (OS, *i.e*., Mac, Linux, or Windows), directory names must be separated by forward slashes (/); functions in the Python *operating system* module generate OS-appropriate paths from the user-provided string.

The remaining five user-entered values populate Python variables that are printed to Sample Sheet sections to customize content (detailed below as *workflow specifications* and *sample:barcode assignments*). Sample Sheets require three sections—denoted in the *.csv file by the bracketed strings [Header], [Reads], [Data]—and optionally include additional sections (*e.g.*, [Settings], [Manifests]). SampleSheet.py generates Sample Sheets that use four of these sections: [Header], [Reads], [Settings], [Data] (**Supp. Fig. 2**).

#### 3−6-Workflow specifications

[**Header**]—[Header] and [Settings] demarcate lines of comma-separated key:value pairs that encode metadata for the sequencing experiment; each key denotes a metadatum type and each value encodes a corresponding metadatum. SampleSheet.py prints values for eight metadata keys under [Header]: *IEMFileVersion, Investigator Name, Project Name, Date, Workflow, Application, Description*, and *Chemistry*. Values for two keys (*Investigator Name, Project Name*) are user-supplied at console prompts during script operation (**Fig. 2a**, **inputs #3-4; Supp. Fig. 3a-3**), with Date value auto-generated based on the system’s present calendar time. Five keys default to values appropriate for amplicon sequencing on Illumina® instruments (*“IEMFileVersion,4”, “Workflow,GenerateFASTQ”, “Application,FASTQ Only”, “Assay,Nextera”, “Description, Sequencing”*, and *“Chemistry, Amplicon”*).

[**Reads**]**—**The number of nucleotide-step extension and imaging cycles to be completed by the sequencer is specified in the numeric values on lines of a Sample Sheet’s [Reads] section: a single line (*e.g*., 151) communicates that 150 cycles of base acquisition (following a +1 phasing cycle) will be completed to generate a single read (single-end run); two lines (*e.g*., 151 \n 151, where ‘\n’ represents newline character) communicates that 150 cycles will be completed in two directions to generate forward and reverse-complement reads (paired-end run). Users enter the read cycle number(s) to be printed in the Sample Sheet [Reads] section as a single line of comma-separated text (two or three values) at the console prompt, specifying single-end or paired-end run (SE or PE) and cycle number(s) (single number for SE run, two numbers for PE run) (**Fig. 2a**, **input #5; Supp. Fig. 3a-4**). For example, for a single-end run with 35 read cycles (plus a single phasing cycle), a user would enter *SE, 36.* For a paired-end run with 150 read cycles in each direction, a user would enter *PE, 151, 151*.

[**Settings**]**—**The 96×96 oligonucleotides reported here—and whose 8-nt index sequences are embedded in SampleSheet.py data objects—are designed with Nextera sequences flanking the target read sequence (**Supp. Fig. 1**)^25^, meaning that the adapter sequence *5’-CTGTCTCTTATACACATCT-3’* defines the end of amplicon-specific read sequences. SampleSheet.py populates the optional [Settings] section of the Sample Sheet with the key:value pair that communicates adapter trimming during fastq processing (*“Adapter,CTGTCTCTTATACACATCT”)*. *“ReverseComplement,0”* specifies that read sequences are returned as sequenced, not as reverse complements.

##### Sample:bar code assignments

[**Data**]—SampleSheet.py’s key utility is to automate large numbers of sample:barcode relationships in the [Data] table section of a Sample Sheet. In principle, i7 and i5 primers can be used in any combination with one another, but in the workflow described, i7 and i5 primers serve defined sample barcoding roles that are fundamental to the expansion operations performed by SampleSheet.py. Specifically, we assign i7 barcodes as labels for each well (A01-H12) in a 96-well plate, and i5 primers as labels across all wells of a single plate (**Fig. 1a, Fig. 2c**).

These explicit uses of i7 and i5 barcodes are put into practice when preparing amplicons by PCR (**Fig. 1a, Supp. Fig. 1, Supp. Fig. 5**). Samples to be PCR-amplified can be pictured as being arrayed in 96-well plates, with each sample adopting a defined well ID in a defined plate. Samples may be arrayed in 96-well plates in PCR1, which amplifies the target locus from a genetic sample source (such as isolated DNA, cDNA, genomic prep from crude lysate). i7 and i5 barcodes may then be pictured as being overlain on PCR1 amplicons in PCR2: a unique i7 primer is used for each well of a single plate, and therefore i7 defines the well ID of PCR1 plate position; a unique i5 primer is used across all wells of a single plate, and therefore i5 defines the plate ID of a multi-plate PCR1 effort (**Fig 1a, Fig. 2c, Supp. Fig. 5**), and the script outputs each unique full sample name as “Plate name-Sample well position”. Plate name and i7, i5 oligo IDs are entered by user at console prompt (**Fig. 2a, input #6; Supp. Fig. 3a-5;** see *Operations* below for details). SampleSheet.py understands relationships between i7, i5 well ID and barcode sequence, with expectation of i7 barcodes defining wells and i5 barcode defining plate (**Fig. 2b,c**).

### Operations and Sample Sheet output file

SampleSheet.py establishes sample ID:bar code pairings based on user input, and creates a comma-separated file interpretable by Illumina® software as a Sample Sheet. After a user is prompted to specify an absolute path to the intended Sample Sheet file location, the script presents a console table-view of the i7 and i5 barcode sequences (“plateviews”), displaying relationships between the numbers 1-96, well IDs between A01-H12, and corresponding barcode sequences. In CLI format, console table-views require prior installation of PrettyTable, a Python library that supports visual representation of tabular data (freely downloaded from the Python Package Index (PyPI) at https://pypi.org/project/PrettyTable/ or from GitHub at https://github.com/jazzband/prettytable); in Jupyter notebook format, plateviews are automatically presented as images returned after user-definition of variables. In CLI format, SampleSheet.py checks for PrettyTable installation in the system path and a user can choose to bypass PrettyTable at a console prompt that queries whether the user wishes to progress in the script in its absence if PrettyTable cannot be found. Console plateviews represent the script’s effort to provide a user with resources that help to construct accurate sample ID:barcode assignments at the key [Data] user input step in the program (**Supp. Fig 1**). Screen captures of the console plateviews meant to facilitate barcode ID entry can also be found in **Supp. Fig. 6** (CLI and Jupyter notebook formats).

The final prompt for user input requests a list of sample names (*e.g*., overarching sample names uniquely assigned to each 96-well plate) that are comma-separated from integers that specify i7 and i5 barcode assignments to wells. Each line defines a plate name that will be shared across associated well IDs (*e.g*., A01-H12), range of i7 barcodes, and single i5 barcode that uniquely encompass up to 96 barcoded samples in arrayed format. For example, in the following six lines of text,

DG-1, 1-96, 1

DG-2, 1-96, 9

DG-3, 1-50, 78

DG-4, 1-96, 7

DG-5, 1-96, 22

DG-6, 1-68, 34

‘DG-1’ represents a sample plate name, ‘1-96’ represents the range of i7 barcodes used in PCR2 (*i.e*., all 96 i7 primers/barcodes were used (A01-H12), each uniquely labeling a distinct and corresponding well (A01-H12) within the sample plate), and ‘1’ represents the single i5 primer/barcode (from i5 source plate well A01) used to label all wells within this sample plate. SampleSheet.py anticipates for i7 primers to be repeatedly used across 96-well plates to specify individual wells within each sample plate, and an i5 primer to be uniquely assigned to each sample plate to specify the overarching source of i7-labeled wells within each plate. In CLI format, input can be entered line-by-line by a user until a list of entries is complete (a single newline keystroke advances for entry of next sample and barcode range; two consecutive newline keystrokes complete list entry advance the script), or as a single block copied and pasted from advance preparation in a text editor. In Jupyter notebook format, input must be entered with each line entry separated from others by a semicolon character (‘;’, see Jupyter notebook Markdown (GitHub) for details).

From this minimal syntax for sample plate IDs, i7, and i5 barcodes, the six lines of text in the example above are converted to 502 entries in Sample Sheet format; each sample plate is ‘expanded’ to delineate up to 96 individual wells based on the minimal information provided in the i7/i5 range(s) provided as input (*e.g*., DG-1-A01, DG-1-A02, … DG-1-H12, etc.) (**Fig. 2c-d**). For 9,216 entries in Sample Sheet format, 96 lines of text would be required, significantly reducing the labor and potential for erroneous sample:index assignments associated with manual entry of 9,216 assignments. Upon sample:barcode range entry at the command-line, generation and completion of the corresponding Sample Sheet text file is nearly instantaneous (<1 second).

#### ImputedGenotypes.py: allele definition and genotype imputation at specific loci of individual clones

Short-read SBS can yield high-coverage definition of multiplexed sequences. Reads can be digitally tracked as representations of individual template molecules, meaning that a sample with mixed templates (>1 distinct template sequence) can be deconvoluted to restore the identity and relative abundance of the original templates. Allele definition and genotype imputation at experimentally examined loci are common goals of population genetic analyses, including identification of Cas9-edited clones. We developed ImputedGenotypes.py, a script that converts reads in individual sample-specific fastq files into imputed genotypes for those samples at PCR-amplified (queried) loci (**Fig. 1b**, **Fig. 3**). The script defines alleles based on relative read abundance (frequencies), imputes corresponding sample-specific genotypes, and delivers up to eight output files in a user-defined directory location.

### User inputs and program dependencies

ImputedGenotypes.py prompts users for up to seven values—five required and two optional—entered as text at Jupyter notebook or CLI prompts (**Fig. 3, Table 2**). These include: absolute paths to 1) input and 2) output directories, 3) BLASTN executable, and 4) alignment reference database; 5) reference database file prefix; and (optional) 6-7) DNA sub-sequences to display on alignments. At the outset of ImputedGenotypes.py, a user can choose whether to enter input values at ‘coached’ prompts (‘Prompt’) or in a single entry (‘List’) that is parsed by the script into appropriate variables.

**Table 2.**
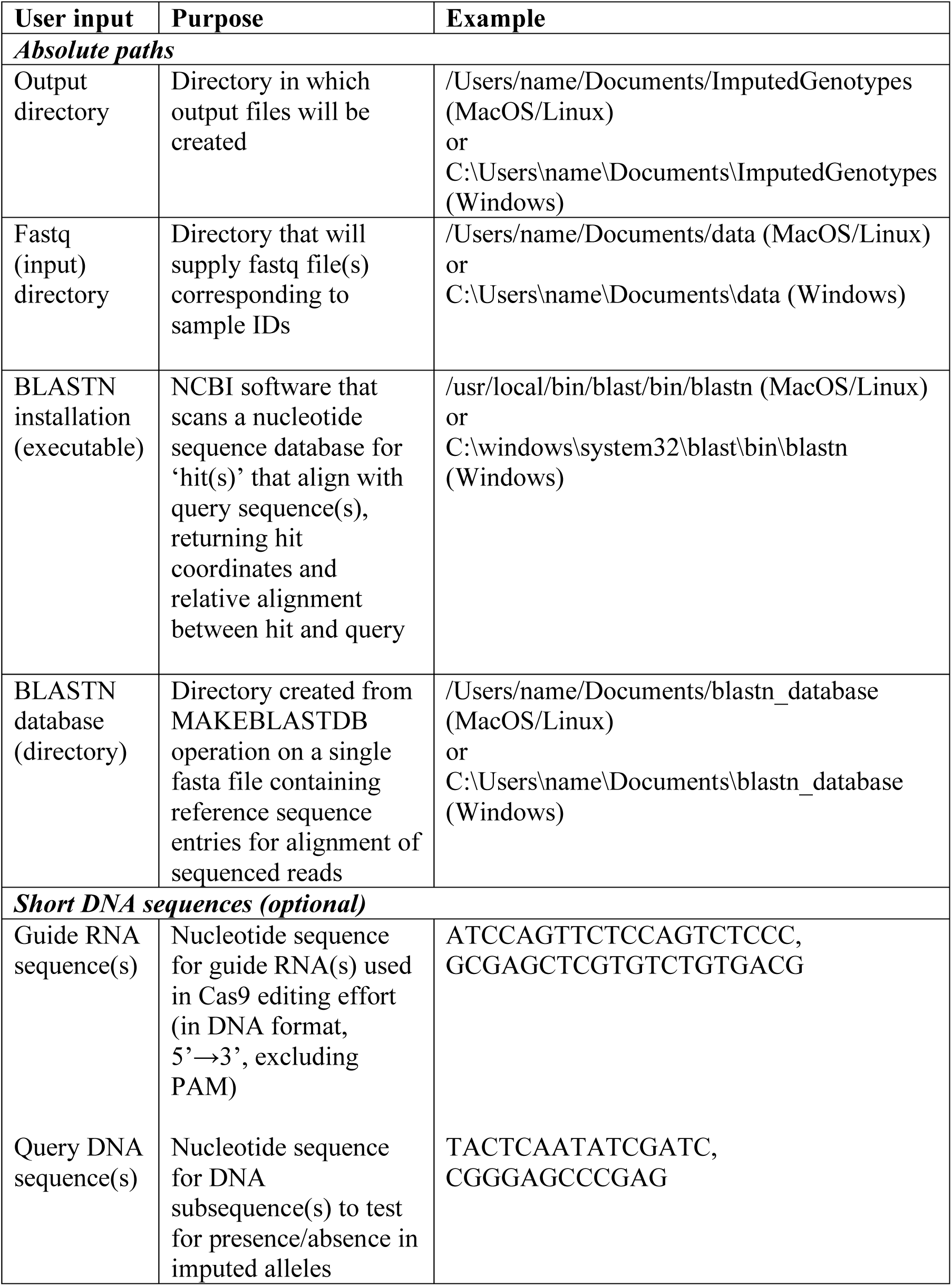
ImputedGenotypes.py inputs.

#### 1− Absolute path to file output

Users are first prompted to enter the location of a directory for output files (absolute path to target destination, empty of files) (**Fig. 3a**, **input #1; Supp. Fig. 3b-1**). The directory can either pre-exist (as long as it is empty), or does not have to pre-exist (the script will create the directory designated by the absolute path if it does not yet exist). Up to eight files (six .txt, one .pdf [optional], one .csv) will ultimately be generated in this directory as script output (**Table 3**).

**Table 3.**
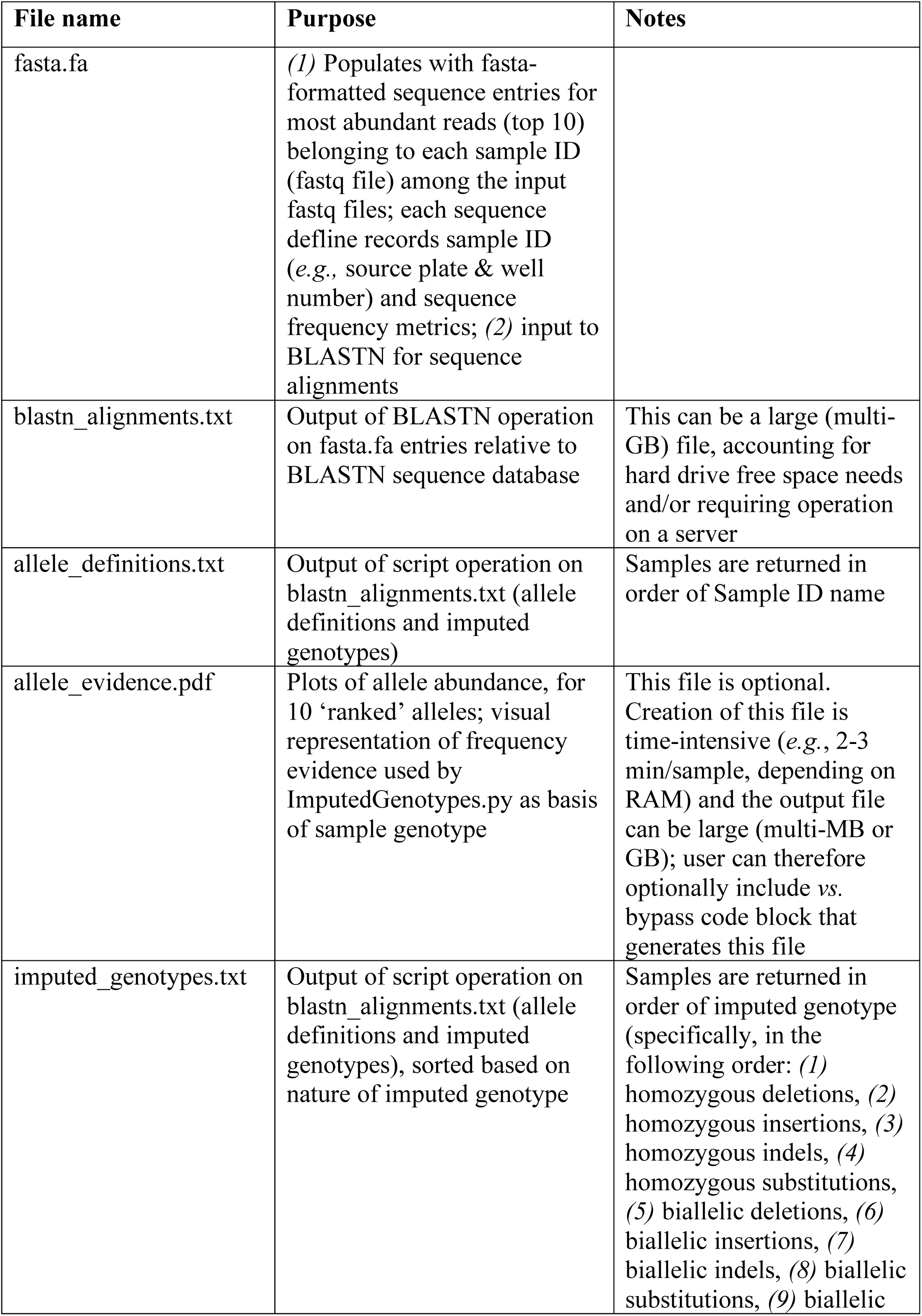
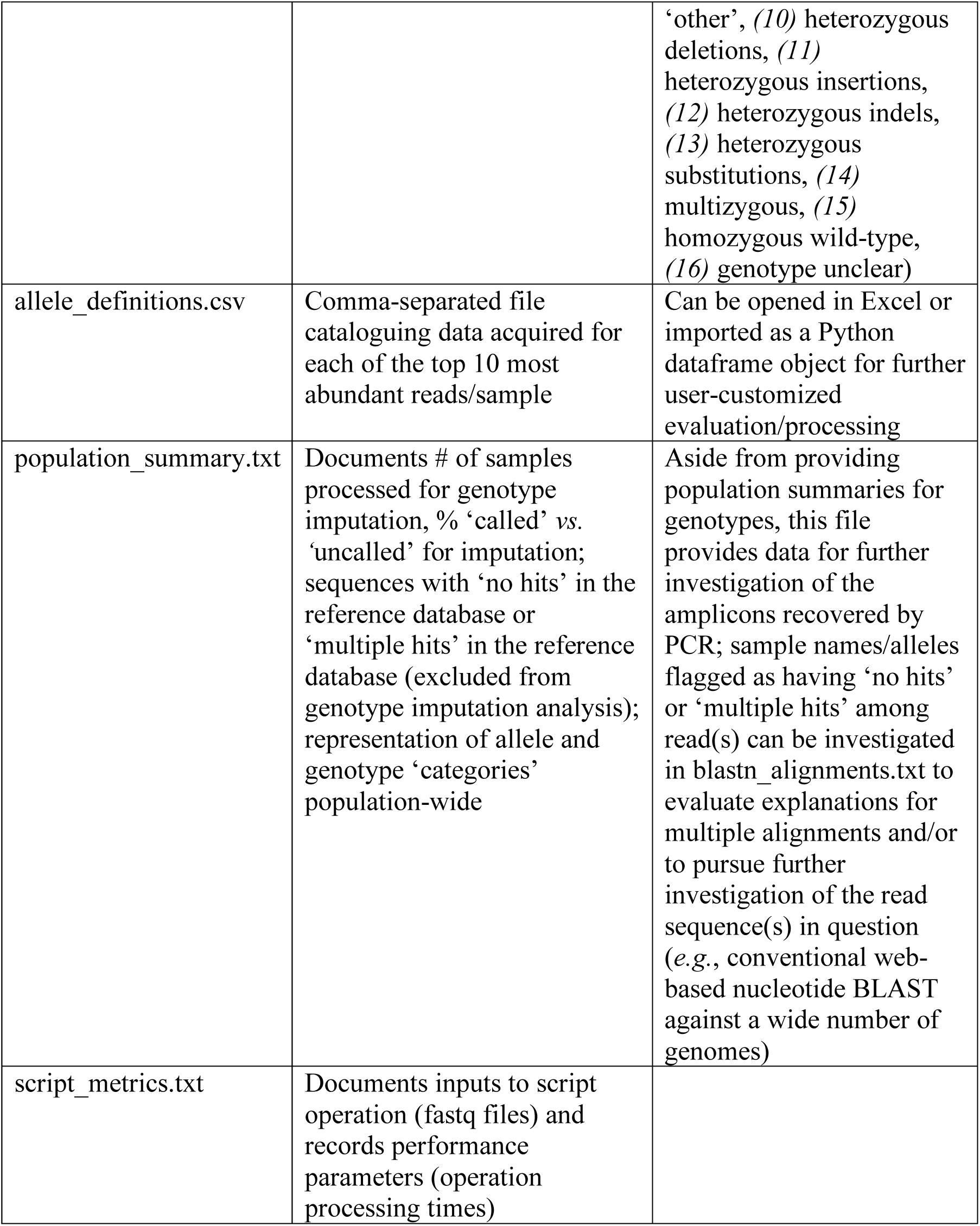
File outputs of ImputedGenotypes.py.

#### 2−Absolute path to file input

The script then requests location of the source file directory—a directory populated with fastq files containing reads derived from amplicon sequencing (**Fig. 3a**, **input #2; Supp. Fig. 3b-2**).

#### 3−Absolute path to BLASTN executable

ImputedGenotypes.py aligns the top 10 reads (abundance defined by frequency) from each fastq file to a reference genome using BLASTN, requiring local pre-installation of BLASTN^20^ (**Fig. 3a**, **input #3; Supp. Fig. 3b-3**; freely available for download with the BLAST+ suite at https://www.ncbi.nlm.nih.gov/guide/howto/run-blast-local/.

#### 4−Absolute path to reference sequence (e.g., genome) database

BLASTN requires a local reference sequence database for alignment operations, typically a set of six files with a common prefix (*e.g*., *‘*GRCh38’) and extensions .nhr, .nin, .nog, .nsd, .nsi, .nsq, generated from a single fasta file containing one or more entries (for example, in the case of a database source file GRCh38.p13_genomic.fna, 457 fasta entries)^26^ (**Fig. 3a**, **input #4; Supp. Fig. 3b-4**). A user-specified genome database (or customized sequence database) is a single directory containing these six files, and can be made by supplying a fasta file containing the target sequence(s) from which BLASTN will seek alignments to MAKEBLASTDB, a CLI program available in the BLAST+ download suite (usage guidelines described in the BLAST Command Line Applications User Manual, https://www.ncbi.nlm.nih.gov/books/NBK279688/).

#### 5−Prefix common to the six files that compose the alignment reference database

The alignment reference database comprises six files with a common prefix; the script requests this prefix from the user (**Fig. 3a**, **input #5; Supp. Fig. 3b-5**).

#### 6& 7-Optional DNA subsequence(s)

Finally, users have the option to supply one or more short nucleotide sequences to be mapped/superimposed above or below allele alignments, if matches are found in the aligned nucleic acid sequences. A Jupyter notebook or CLI prompt first asks whether a user will supply entries for one or both of up to two optional subsequence inputs; these include: *(1)* ‘guide RNA sequence’ (5’→3’, in DNA form, excluding PAM), if user wishes to display position of guide RNA used in Cas9 editing effort (can be useful to gauge the plausibility of an allelic difference relative to reference sequence as being consequence of Cas9-induced break) (**Fig. 3a**, **input #6; Supp. Fig. 3b-6**), and *(2)* ‘test sequence’ (5’→3’), if user wishes to query for presence or absence of a specific subsequence in an allele identified by deep sequencing (relative to a reference, *e.g.*, wild-type, allele) (**Fig. 3a**, **input #7; Supp. Fig. 3b-7**).

##### Other dependencies

Generation of read frequency statistics requires installation of Python NumPy and SciPy libraries (https://www.scipy.org/scipylib/download.html, https://pypi.org/project/numpy/); generation of a PDF file with allele frequency plots (optional) requires Python fpdf and PyPDF2 packages (https://pypi.org/project/fpdf/, https://pypi.org/project/PyPDF2/).

### Operations and output files

Foremost, ImputedGenotypes.py converts raw sequencing read data to proposed allele and genotype representations for demultiplexed samples. Its key operations center on discernment of distinct reads, assessment of relative read frequencies, definition of proposed alleles as wild-type or mutant relative to an alignment reference, and genotype hypotheses based on ranked allele abundances. Four lists and two principal dictionaries organize the data analyses that ultimately appear in sample and population genotype summaries. Its key outputs supply, for user evaluation, visually accessible evidence for allele definitions and hypothesized genotypes (**Table 3, Supp. Fig. 7**).

#### fasta.fa

Core operations begin with fastq file processing, channeling the top ten ranked read types and their quantified frequency metrics to a fasta text file (*fasta.fa*), the input for BLASTN alignments. For each sample, every read sequence is collected in a temporary Python list (*read_lines*), evaluated by the Python Counter function to identify the top ten most represented reads and their frequency metrics expressed in five ways: (1) read count/total sample reads, (2) read percentile rank relative to other reads, (3) % read abundance (raw), (4) % read abundance relative to reads that occur at >1% frequency, (5) % read abundance relative to reads that occur at >10% frequency (**Fig. 3b**). The fasta description line (defline) for each read sequence ingrains both sample ID and frequency metrics (sample ID = Sample_Name defined in Sample Sheet [Data] section and fastq file name), embedding values used in upcoming script operations to assess read contribution to genotype imputation (defline structure: *>samplename-plate-well_R1orR2_*[*read count/total reads for sample*]*_% of all reads_percentile rank relative to all reads_% of all reads adjusted for reads that occur >1%_% of all reads adjusted for reads that occur >10%*) (**Fig. 3c**).

#### blastn_alignments.txt

The script passes *fasta*.*fa* to BLASTN using Python’s System Command function, accessing user-specified paths to the BLASTN executable (*input #3*), the reference sequence database directory (*input #4+input#5*), and *blastn_alignments.txt* with output settings *-gapopen 1 -gapextend 1 -outfmt 5*. Because *blastn_alignments.txt* can be a large file (*e.g.*, several GB depending on the number of fasta entries aligned), users must have sufficient hard drive space available and memory resources to accommodate operations that draw on this file (alternatively, submit ImputedGenotypes.py to a computing cluster with sufficient memory and storage resources).

The alignment content of *blastn_alignments.txt* is populated into a series of Python list objects, in which alignment data are parsed and reformatted (*e.g.*, filtered of queries flagged by ‘No hits found’ and queries that identified multiple hits (‘<Hit num>’ >1) in the reference database). Query sequences that belong to the same sample ID (sequences among the top ten ranked reads for a sample) are grouped and assigned to unique sample ID in a dictionary, *alignmentoutput_dict2*.

#### allele_definitions.txt, imputed_genotypes.txt, and allele_definitions.csv

ImputedGenotypes.py draws from *alignmentoutput_dict2* to populate its core dictionary, *imputedgenotypes_dict*, in which data are ultimately evaluated for sample allele definitions and genotype imputation. For each sample, *imputedgenotypes_dict* compiles four subdictionaries assigned to each ranked sequence (candidate allele): subdictionary 1 records ‘allele_name’ (fasta defline), ‘chr+build’, ‘locusID’, ‘coordinates’, and ‘alignment’; subdictionary 2 records ‘allele_type’ (*e.g*., wild-type, mutant) and ‘allele_specs’ (‘specifications’, *e.g*., likely deletion, insertion, substitution, indel), subdictionary 3 records guide RNA sequence(s) with match position in reference sequence, and subdictionary 4 records DNA test sequence(s) with match position in reference sequence. Finally, imputed genotype—based on ranked allele type and specification for alleles with >10% adjusted frequency—is assigned to each sample ID (*e.g*., homozygous wild-type, homozygous deletion, heterozygous deletion, multi-allelic, etc. (**Table 3**))

ImputedGenotypes.py displays its findings in two text files, *allele_definitions.txt* and *imputed_genotypes.txt*; for each sample ID, *allele_definitions.txt* reports imputed genotype, followed by ‘alleles’ (up to ten sequences ranked by relative frequency) identified from read 1 (R1) and read 2 (R2) sequences (**Fig. 3d, Supp. Fig. 7**). Three text blocks report frequency metrics, allele specifications, and alignments: (1) ‘*Allele’* reports sequence name (fasta defline containing sample ID and frequency metrics) and allele specifications (definition as ‘wild-type’ or ‘mutant’ relative to reference sequence (BLASTN ‘hit’), and if ‘mutant’, further resolution as ‘likely deletion | insertion | substitution | complex indel’, including number of bp altered by the mutation (*e.g*., ‘8 bp’); (2) ‘*Locus’* reports details of alignment database ‘hit’, defined by BLASTN database content (typically, locus identifier and coordinates); (3) ‘*Alignment’* reports allele sequence relative to reference ‘hit’ sequence, with midline (pipe (‘|’) *vs.* gap) reporting matched *vs.* unmatched nucleotide positions. If DNA sub-sequences were provided for annotation (*optional inputs #6 & 7*), ImputedGenotypes.py maps the position(s) of these sequence(s) above (if guide RNA) or below (if test sequence) each alignment in which the sequence(s) were identified in an allele or its reference, facilitating interpretation of indels as plausible consequences of Cas9-directed mutagenesis, and/or assessment of test sequences for presence *vs.* ablation. Ranked sequences that occurred at adjusted frequency <10% are demarcated from other alignments with text highlighting their scarceness as likely artefacts (not representing genetic source sequences): ‘*>>>>> remaining alleles occur at frequency <10% <<<<<’*.

Like *allele_definitions.txt*, the file *imputed_genotypes.txt* reports allele definitions and alignments for ranked alleles, but reports samples based on imputed genotype class. In other words, a ‘homozygous deletion’ cohort is reported before a ‘homozygous insertion’ cohort, in turn reported before ‘heterozygous’ cohorts, followed by a ‘homozygous wild-type’ cohort and finally, a cohort for which insufficient evidence was recovered to impute genotype (‘unclear or multi-allelic, insufficient representation of allele(s)’) (**Table 3**). This format provides an organized list of identified mutant clones, useful for further experimental processing and long-term storage. In addition to their summation in *allele_definitions.txt* and *imputed_genotypes.txt*, allele data compiled in *imputedgenotypes_dict* are transferred to a pandas dataframe for output in *allele_definitions.csv*, for convenient user access to raw data (**Supp. Fig. 7**).

#### allele_evidence.pdf (optional)

ImputedGenotypes.py visually reports R1 and R2 ranked sequences in frequency plots printed to *allele_evidence.pdf* (**Fig. 3d, Supp. Fig. 7**), an optional output file that highlights the ranked sequence subset representing alleles most likely to account for sample genotype. For each sample, ranked sequence abundance is rendered as (1) raw frequency (% total reads), (2) % top 10 reads, (3) % reads filtered for reads occurring at >1% raw frequency, (4) % reads filtered for reads occurring at >10% raw frequency. A sizeable fraction of reads that occur at <1% frequency in a fastq file are attributable to template differences introduced by sequencing or PCR artefacts^27^, justifying their exclusion in plots (3) and (4) and the frequency recalibration of more abundant sequences. Generation of these plots can be time-intensive (*e.g.*, ∼2 min. per pdf page depending on system resources), and this code passage is therefore optional in ImputedGenotypes.py; after initial user input and just before script operations begin, a user is prompted to specify whether to include (‘Y’) or bypass (‘N’) frequency plot generation and assembly into a pdf file.

#### population_summary.txt

ImputedGenotypes.py chiefly hypothesizes genotypes for individual samples demultiplexed from a potentially diverse library population, but the program also reports aggregate population properties in *population_summary.txt* (**Supp. Fig. 7**). In ‘Synopsis of Interpretations: Allele Definitions & Genotype Imputations’, the script catalogs *i)* the fraction of samples for which a genotype was imputed, *ii)* overall genotype properties represented in the sample population (*e.g.*, % samples diploid (1-2 prominent alleles inferred) *vs.* % multiploid (>2 prominent alleles inferred), % homozygous wild-type *vs.* homozygous mutant (subsetted for deletion, insertion, substitution, complex indel), % heterozygous (wt + mutant), subsetted as above, % heterozygous (mutant + mutant), etc.), and *iii)* overall alleles represented (*e.g.*, % wild-type alleles, % mutant alleles (deletion, insertion, substitution, complex indel). In ‘Synopsis of Reads Lost to Analysis’, the script earmarks ranked reads for which there were *i)* no hits, or *ii)* multiple hits, in the reference database (for sequences with ‘no hits’, a user may wish to use BLAST online to identify non-target sequence that was detected as amplified from sample source; for sequences with multiple hits, a user may choose to recast (constrain) the reference database to focus target alignment, and/or may choose to redesign primers or PCR conditions to improve specificity in future amplicon libraries for the locus in question).

#### script_metrics.txt

Finally, ImputedGenotypes.py logs script operation parameters in *script_metrics.txt*, preserving *i)* operating system information (name, platform, RAM (GB), physical CPU/effective CPU, Python executable), *ii)* user-defined variables (output_directory, fastq_directory, blastn_path, db_path, db_prefix, guideRNA_seq, extant_seq), fastq file properties (*e.g.*, Illumina run ID(s), # of fastq files processed and their size and read distribution), *iii)* file output information (output directory, files and their sizes), and *iv)* script operation times (*e.g.*, start time, fasta processing time, alignments processing time, imputation processing time, frequency plots compilation time, etc.) (**Supp. Fig. 7**).

##### CollatedMotifs.py: identification of altered TFBS in individual mutant clones

DNA sequence-selective transcriptional regulatory factors (TFs) interact with genomic response elements (alternatively denoted as enhancers or *cis*-regulatory modules), chromosomal regions that confer transcriptional regulation, each containing transcription factor binding sites (TFBS) for distinct combinations of TFs. ENCODE catalogues genome-wide occupancy^28^ positions for such TFBS clusters within genomes, but few have been validated as functional response elements. Cas9 editing routinely yields mixed allelic mutation at target loci (*e.g*., variable insertion *vs.* deletion, indel length across edited cells). For editing efforts targeted to putative response elements, widely available pattern-matching tools enable prediction of TFBSs in a query sequence based on matches to position frequency matrices of known TFs. We developed CollatedMotifs.py, a script that automates identification and comparison of TFBS motifs between sample-specific alleles and a user-supplied reference sequence.

### User inputs and program dependencies

CollatedMotifs.py prompts users for ten required values—nine absolute paths to directories, files, or executables, plus one prefix for alignment database files—entered as text at Jupyter notebook or CLI prompts (**Fig. 4, Table 4**).

**Table 4.**
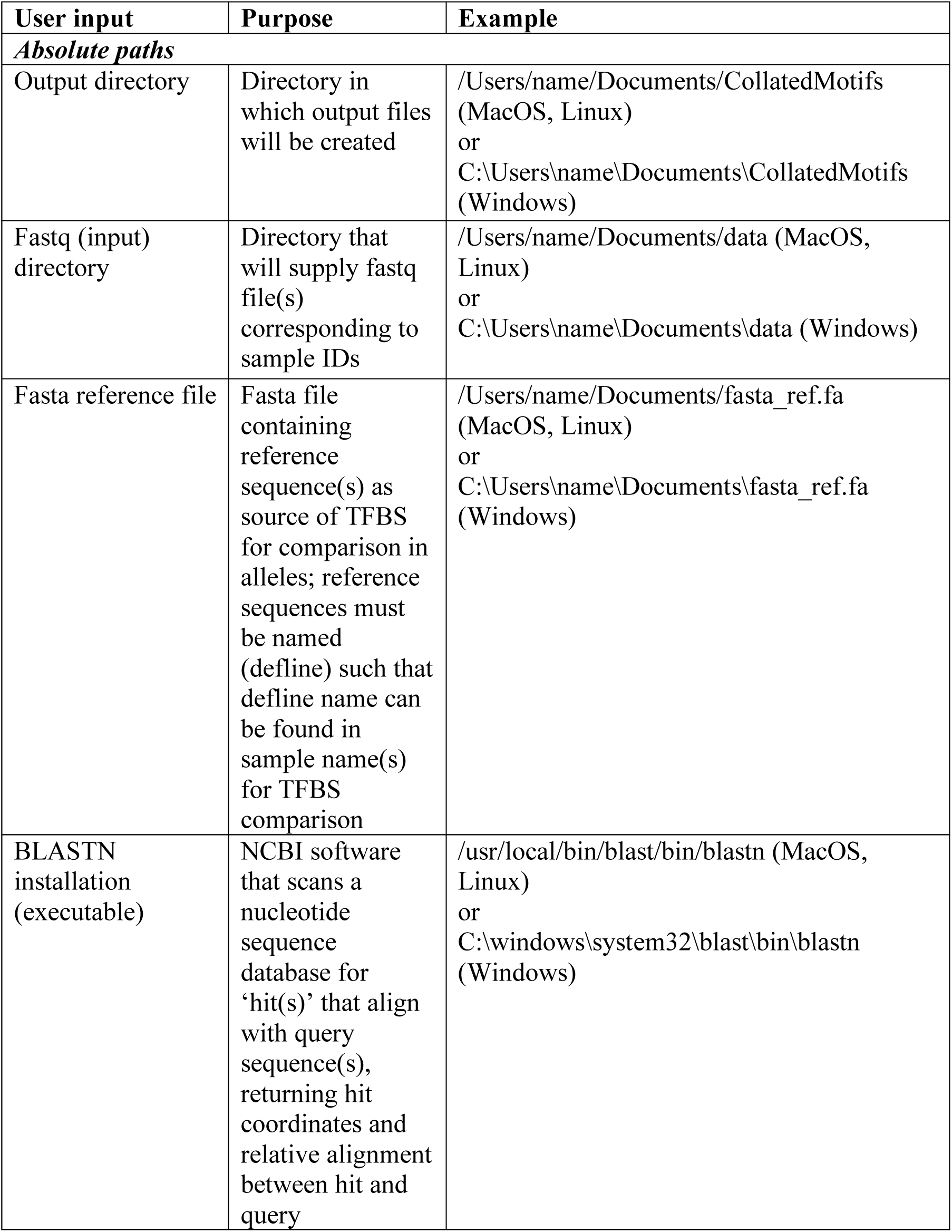
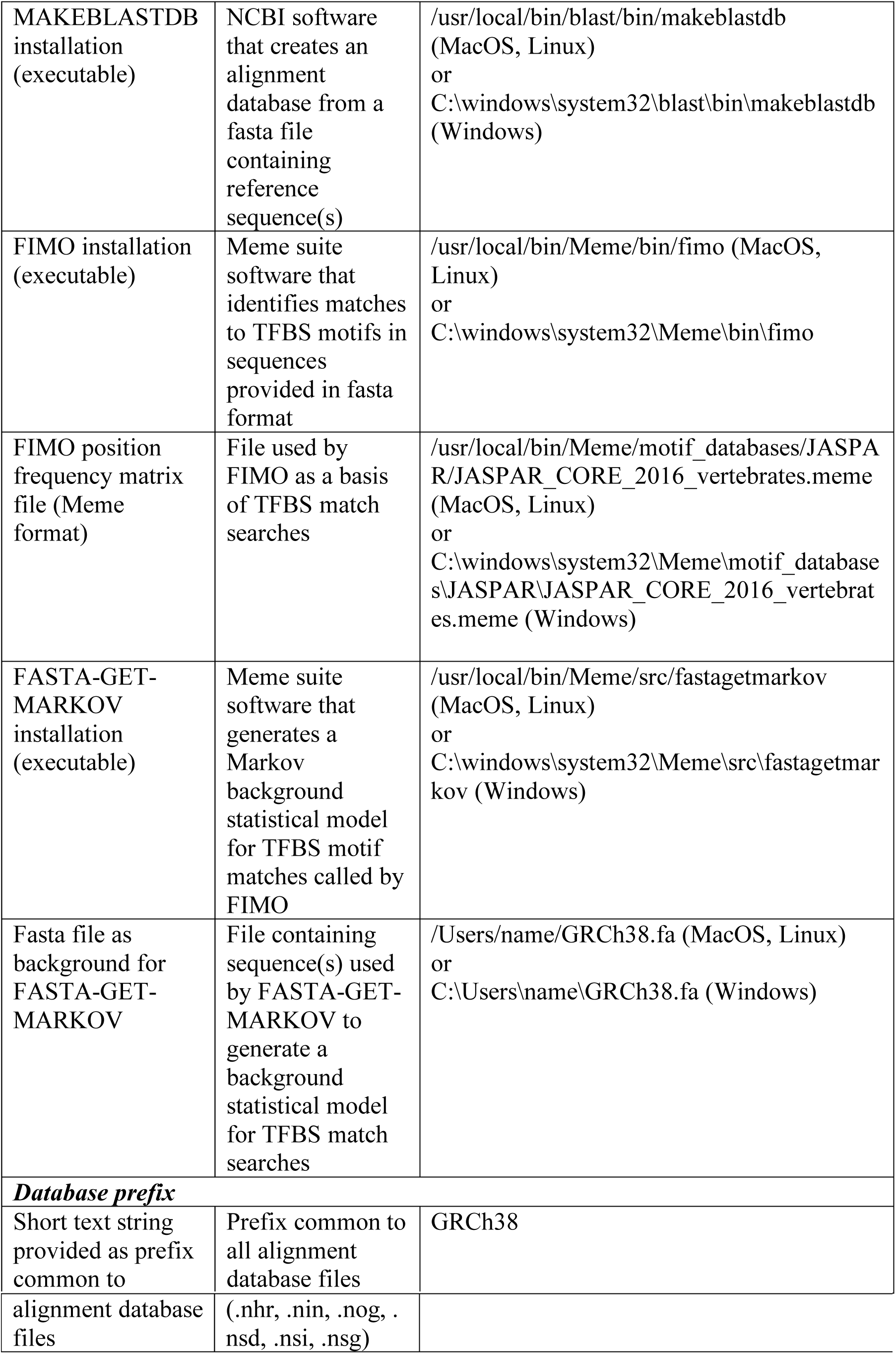
CollatedMotifs.py inputs. Note that in Windows OS, Meme suite programs (FIMO and FASTA-GET-MARKOV) require virtualization and CollatedMotifs.py must be run from within a hypervisor (*e.g.*, Oracle VirtualBox; Open Virtualization Format file available at DOI 10.5281/zenodo.3406862).

#### 1-Absolute path to output directory

Users are first prompted to enter the location of a directory for output sub-directories and files (absolute path to target destination, empty of directories and files) (**Fig. 4a, input #1; Supp. Fig. 3c-1**). The directory can either pre-exist (as long as it is empty), or will be created by the script if it does not yet exist. Three sub-directories (*alignments_database, fimo_out* and *fimo_out_ref*) and five files will ultimately be generated in this directory as script output (**Table 5**).

**Table 5.**
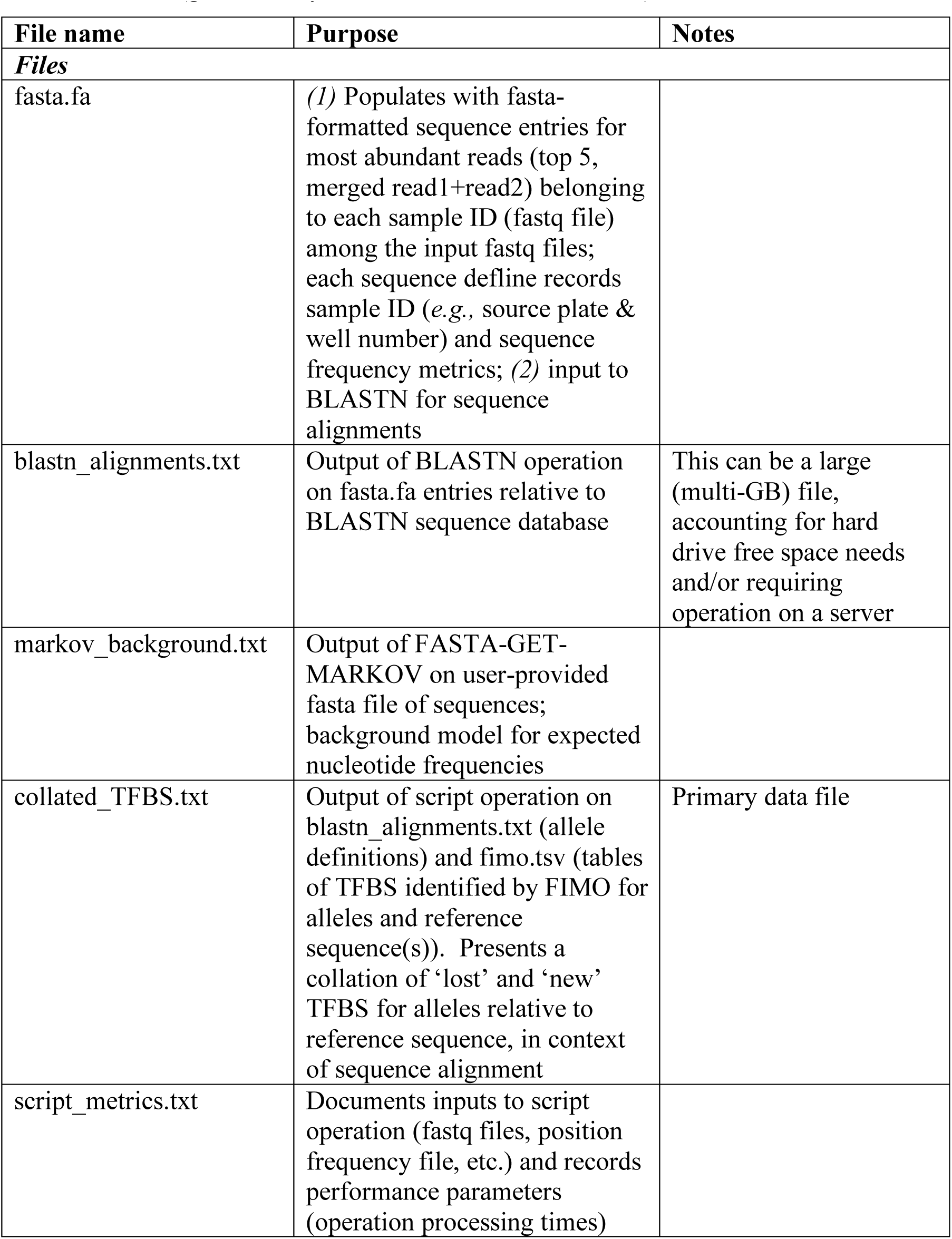
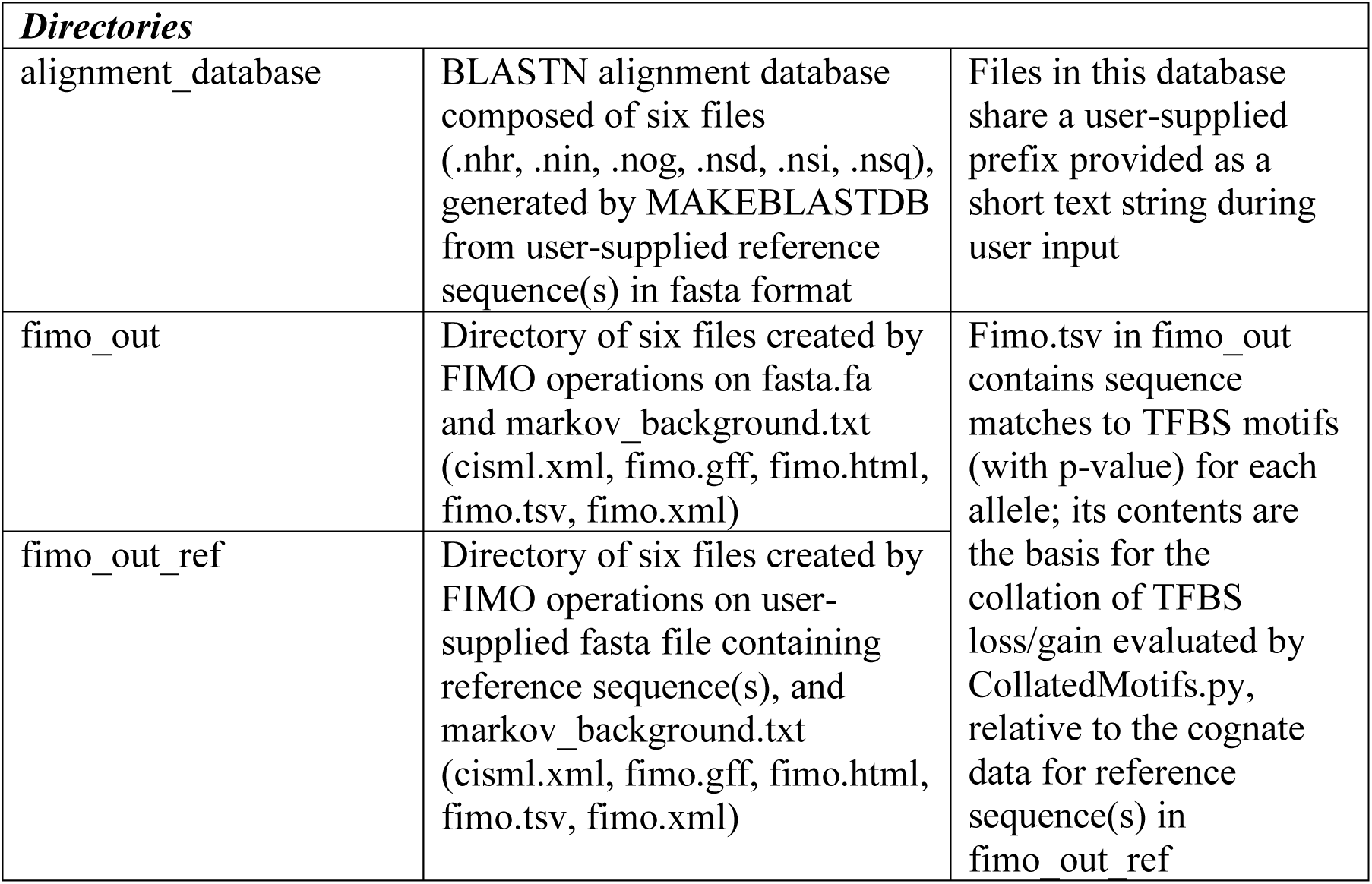
File outputs of CollatedMotifs.py. The script generates 5 separate files, plus three directories (generated by MAKEBLASTDB and FIMO).

**Table 6.**
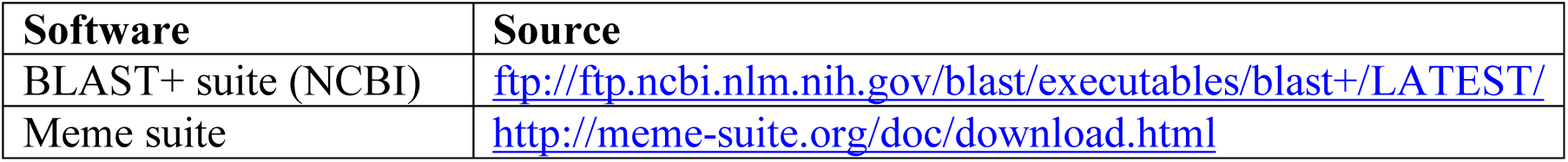
Public sources for software dependencies.

| **Software** | **Source** |
| --- | --- |
| BLAST+ suite (NCBI) | ftp://ftp.ncbi.nlm.nih.gov/blast/executables/blast+/LATEST/ |
| Meme suite | http://meme-suite.org/doc/download.html |

#### 2-Absolute path to fastq files

The script then requests location of the source file directory—a directory populated with sample-specific fastq files containing reads to be processed for sequence content and matches to TFBS motifs (**Fig. 4a, input #2; Supp. Fig. 3c-2**).

#### 3-Absolute path to reference fasta file

CollatedMotifs.py relies on user-supplied reference sequence(s) to evaluate read (allele) sequence properties, specifically to (1) run sequence alignments and (2) compare TFBS. Users supply reference sequence(s) (≥1) in NCBI fasta format in a single fasta-formatted text file (**Fig. 4a, input #3; Supp. Fig. 3c-3**). The span of each user-supplied reference sequence should correspond to the full genomic amplicon prepared for deep sequencing (borders defined by 5’-most complementarity of primers to target sequence); each associated fasta defline should be designated by a *useful name or string* that can be matched in entirety in fastq sample name(s) scheduled to be compared to the fasta reference sequence in question (for example, for sample fastq filenames containing the shared prefix ‘DG-1’ (*e.g.*, DG-1-A01, DG-1-A02, … DG-1-H12), the fasta defline for the reference sequence assigned to DG-1 reads should appear as, ‘>DG-1’). CollatedMotifs.py provides these sequences to MAKEBLASTDB to generate a custom sequence database for alignments, and also to FIMO, to generate reference-specific TFBS lists for allele comparisons.

#### 4-Absolute path to BLASTN executable

CollatedMotifs.py aligns the top 5 reads (abundance defined by frequency) from each fastq file to a reference sequence database (*alignments_database*) using BLASTN^20^ (available for download with BLAST+ suite) (**Fig. 4a**, **input #4; Supp. Fig. 3c-4**).

#### 5-Absolute path to MAKEBLASTDB executable

BLASTN requires a local sequence database for alignment operations, a set of six files generated from a single fasta file containing one or more entries (see corresponding entry in ImputedGenotypes.py description for further detail)^26^. Whereas a user of ImputedGenotypes.py prepares a genome sequence database using the CLI program MAKEBLASTDB in advance of script operation (**Fig. 4a, input #5; Supp. Fig. 3c-5**), users of CollatedMotifs.py provide the absolute path to the MAKEBLASTDB executable (within BLAST+ suite); the script invokes MAKEBLASTDB to generate a database from the sequences in the user-supplied *reference fasta file* (*input #3*).

#### 6-Prefix common to the six files that compose the alignment reference database

The alignment reference database comprises six files with a common prefix; the script requests a user-defined (custom) prefix to be assigned to these files by MAKEBLASTDB (**Fig. 4a**, **input #6; Supp. Fig. 3c-6**).

#### 7-Absolute path to FIMO executable

To identify matches to TFBS motifs, CollatedMotifs.py invokes FIMO (‘Find Individual Motif Occurrences’), a program available in the MEME suite of motif-based sequence analysis tools (Meme 5.0.5 download available at http://meme-suite.org/doc/download.html, FIMO background available at http://meme-suite.org/doc/fimo-tutorial.html) (**Fig. 4a**, **input #7; Supp. Fig. 3c-7**). Users supply the absolute path to the local installation of the FIMO executable.

Note that in Windows OS, Meme suite programs require virtualization, and CollatedMotifs.py must be run from within a hypervisor (*e.g.*, Oracle VirtualBox; Open Virtualization Format file available at DOI 10.5281/zenodo.3406862).

#### 8-Absolute path to FIMO motif file

The TFBS search program FIMO uses a plain-text file containing position frequency matrices for one or more TFs (Meme format), the basis of TFBS identification in user-supplied reference sequences and fastq-supplied (sample ‘query’) sequences. Users can download a directory of motif database files at http://meme-suite.org/doc/download.html (**Fig. 4a**, **input #8; Supp. Fig. 3c-8**). Dozens of files listing position frequency matrices experimentally defined for TFs in eubacteria, archaea, and eukaryotic groups are available in the *motif_databases* directory of a Meme suite download. We used *JASPAR/JASPAR_CORE_2016_vertebrates.meme*, from the 2016 (6^th^) release of the public database JASPAR^29^, as the motif reference file in the example case (containing position frequency matrices for 519 vertebrate TFs).

#### 9-Absolute path to FASTA-GET-MARKOV executable

FIMO requires a background model from which to assess statistical significance of sequence matches to position frequency matrices; CollatedMotifs.py invokes FASTA-GET-MARKOV, a program available within Meme suite download, to generate a background model for FIMO operations on evaluated sequences. FASTA-GET-MARKOV generates a background Markov model from a user-supplied *reference fasta file* (*input #10*) (further description at http://meme-suite.org/doc/fasta-get-markov.html) (**Fig. 4a, input #9; Supp. Fig. 3c-9**).

#### 10-Absolute path to file supplied to FASTA-GET-MARKOV for background model generation

FASTA-GET-MARKOV produces a background model file from a user-supplied, fasta-formatted nucleotide sequence file (**Fig. 4a, input #10; Supp. Fig. 3c-10**; also see *input #9*). We used the full human genome sequence (GRCh38.p13 Primary Assembly) as the fasta reference file supplied to FASTA-GET-MARKOV for background model generation (RefSeq Accession ID: GCF_000001405.39; filename: GCF_000001405.39_GRCh38.p13_genomic.fna; https://www.ncbi.nlm.nih.gov/assembly/GCF_000001405.39).

##### Other dependencies

Generation of read frequency statistics requires installation of Python NumPy and SciPy libraries.

### Operations and output files

Like ImputedGenotypes.py, CollatedMotifs.py reports hypothesized alleles for demultiplexed NGS datasets, but its sequence alignments are populated with matches to TFBS motifs—specifically, with TFBS lost or gained in an allele sequence relative to a user-provided reference sequence. Four dictionaries organize the key data operations that report TFBS collations in the visual context of alignments. Its key outputs prepare data files used by BLASTN and FIMO, and supply visually accessible evidence for allele definitions and associated TFBS comparisons (**Supp. Fig. 8**).

#### Alignment_database and markov_background.txt

Unlike ImputedGenotypes.py, CollatedMotifs.py generates the BLASTN alignment database inline, generating a spartan database derived solely from the sequences provided in the reference fasta file (*input #3*), with six file names prefixed by a custom string (*input #6*). A background Markov file is generated from user-defined sequences (*input #10*), to be provided to FIMO during TFBS match operations. MAKEBLASTDB and FASTA-GET-MARKOV are invoked using the Python System Command function.

#### Fasta.fa

The script proceeds to operations that overlap with the fastq→fasta steps in ImputedGenotypes.py, linking frequency metrics to ranked sequences in *fasta.fa* (**Fig. 3a-c, Fig. 4b-c**). CollatedMotifs.py differs from ImputedGenotypes.py in that (for PE sequencing) read 1 (R1) and read 2 (R2) sequences for each sample are *merged* based on common cluster ID in R1 and R2 fastq files (*i.e*., R1 and R2 reads are not tracked independently, in contrast to ImputedGenotypes.py) (**Fig. 4b**). Merged (R1+R2) sequences are channeled into a temporary Python list and filtered for the five most abundant reads, then populated to *fasta.fa* with fasta defline linking sample ID and frequency metrics to sequences.

#### Blastn_alignments.txt

CollatedMotifs.py passes *fasta.fa* to BLASTN, which generates alignments to sequence content in *alignment_database* (**Fig. 4c**). Alignments in *blastn_alignments.txt* are iterated through a series of Python list objects to filter for ‘no hits’ or ‘multiple hits’, ultimately yielding a dictionary that contains unique sample IDs (keys) linked to tuples (values) comprising the suite of sample ID-derived sequence(s) that aligned to unique loci in *alignment_database*.

#### Fimo_out and fimo_out_ref

The script then advances to identification of matches to TFBS motifs in DNA sequences, invoking FIMO. FIMO separately queries two fasta files*—(1)* the reference sequence file (*input #3*) and *(2) fasta.fa* generated by CollatedMotifs.py; FIMO evaluates sequences in these files for TFBS matches to motifs in the user-supplied position frequency matrix (*input #8*). For each of the two fasta files, CollatedMotifs.py directs five FIMO default output files (*cisml.xml, fimo.gff, fimo.html, fimo.tsv, fimo.xml*) to one of two script-generated subdirectories, *ref_fimo_out* or *fimo_out* (**Fig. 3c**). The five FIMO files present TFBS identification outputs in distinct formats; only *fimo.tsv* is accessed by CollatedMotifs.py, and its contents are read into reference- or allele-respective dictionaries, *dict_ref_TFBS* and *dict_allele_TFBS.* By default, FIMO reports TFBS matches at a p-value threshold of 0.0001 (1e-4), but users can adjust this threshold by adding the flag *--thresh* with a revised value to the script’s FIMO operation call (details for this and other flags can be found at http://meme-suite.org/doc/fimo.html). *Collated_TFBS.txt*—CollatedMotifs.py uses Python dictionary objects to complete its distinctive feature, collation of TFBSs for reference sequence(s) and putative alleles. For each sample-associated, ranked allele in *dict_allele_TFBS*, CollatedMotifs.py determines the appropriate reference sequence with which to pose a comparison in *dict_ref_TFBS.* A compilation dictionary, *dict_allele_TFBS_synopsis*, assembles each sample ID (key) linked to a dictionary (value) containing ranked alleles that each point to further subdictionaries (**Fig. 4**): *i)* ‘TFs’ summarizes the transcription factors with TFBS identified for each allele, *ii)* ‘gained’ and *iii)* ‘lost’ list TFBS that are novel or absent in the allele relative to ‘all_sites’ in *dict_ref_TFBS_synopsis* for the corresponding reference sequence . For each sample ID in *dict_allele_TFBS_synopsis, collated_TFBS.txt* reports a visual mapping of *i)* TFBSs new to each allele above the alignment (retrieved from *alignmentsoutput_dict2*), and *ii)* TFBSs lost from each allele below the alignment (*e.g.*, ‘new TFBS’ and ‘lost TFBS’) (**Fig. 4d**; **Supp. Fig. 8**).

#### Script_metrics.txt

Like ImputedGenotypes.py, CollatedMotifs.py logs script operation parameters in *script_metrics.txt*, specifically preserving *i)* operating system information, *ii)* user-defined variables, *iii)* fastq file properties, *iv)* position frequency matrix file properties (TF metadata), *v)* file output information, and *vi)* script operation times (*e.g.*, start time, MAKEBLASTDB and FASTA-GET-MARKOV processing time, fasta processing time, alignments processing time, FIMO processing time, etc.) (**Supp. Fig. 8**).

##### Use case: Cas9-edited disruptions of glucocorticoid receptor-bound loci near a glucocorticoid-regulated gene, FKBP5

We used the 96×96 barcoded primers and the three computational tools described here to identify and characterize mutants among thousands of Cas9-treated clones in the human adenocarcinoma cell line A549, specifically seeking disruptions of glucocorticoid receptor-occupied regions (GORs) near the glucocorticoid-responsive gene *FKBP5* (Gencode v22 gene ENSG00000096060.13). *FKBP5* is one of the most highly glucocorticoid-induced genes in many systems examined; in A549 cells, the *FKBP5* gene body is characterized by promoter-proximal and intronic^30^ GORs (**Fig. 5a**). GC resistance in humans—associated with recurring lifetime vulnerability to major depressive disorder (MDD) and other brain diseases—is potentially associated with higher induced levels of *FKBP5*^31, 32^. Several GORs proximal to *FKBP5* house GR binding sites (GBS) with high evolutionary conservation across 100 vertebrates examined, suggesting distant emergence of these sequences in the vertebrate lineage and long-term negative selection against changes to these GBS over up to 400 million years (**Supp. Fig. 9**).

We examined consequences of individual disruptions of eight GORs in a 1.5 Mb genomic region (GRCh38/chr6:34,950,000-36,450,000), in which *FKBP5* occurs as the only dex-responsive gene within a putative topological domain comprising ∼400 kb (chr6:35,339,500-35,740,000, also comprising the genes *PPARD, FANCE, RPL10A, TEAD3, TULP1, ARMC12*; *FKBP5* mRNA is induced ∼15-fold within 4 h of dexamethasone exposure (100 nM), the only gene body affected >1.5-fold in this region) (**Fig. 5c** and data not shown). In brief, Cas9 sgRNA sequences were cloned downstream of a U6 promoter in a puromycin-selectable vector expressing *Streptococcus pyogenes* Cas9, producing functional Cas9 RNPs *in vivo* when transfected into cells (RRID:Addgene_62988, procedure detailed in **Supp. Methods**) (**Fig. 5b**); after puromycin selection, single cells from Cas9-treated cell populations were FACS-isolated into individual wells of ninety-six 96-well plates for arrayed clonal expansion and amplicon barcoding (**Supp. Methods, Supp. Fig. 10**). Amplicons from 96-well plates were pooled and sequenced on an Illumina MiSeq with excellent (>99%) barcode pair detection across 9,216 samples (**Supp. Fig. 11, Supp. Fig. 12**). A single list of 96 plate:barcode relationships was prepared in plain text format to present as [Data] input to SampleSheet.py; SampleSheet.py rendered the Sample Sheet used for sequencing and demultiplexing in 0.05 sec (<2 min total user interaction with script).

After sequencing and demultiplexing reads into 9,216×2 (18,432) fastq files containing sample read content, fastq files were submitted to ImputedGenotypes.py in batches corresponding to locus and sgRNA (*i.e.*, individual editing scenarios, 384-480 samples/768-960 fastq files per batch) (**Fig. 5b**). On a laptop machine (Mac OS) with 16 GB RAM and 4 physical CPU, allele definitions and imputed genotypes were completed for each batch and returned in *allele_definitions.txt* and *imputed_genotypes.txt* within 1.2 min (mean), with population statistics completed in *population_summary.txt* in <20 sec (total genotype imputation and text file reports completed within 2 min); visual evidence in the form of frequency plots (*allele_evidence.pdf*) was completed within 6 h. Altogether, genotypes for 9,216 samples were complete within 35 min processing time.

We selected clones based on mutant genotypes at target GORs, and evaluated consequences to *FKBP5* regulation by RT-qPCR (**Supp. Methods**). We batch-processed fastq files in CollatedMotifs.py as for ImputedGenotypes.py; *collated_TFBS.txt* files earmarking matches for up to 519 vertebrate TFs, mapped over inferred alleles for 384-480 in each batch, were complete within 3 min. Altogether, files documenting TFBS for 9,216 samples were complete within 60 min processing time. We found that independently derived clones altered at GR-occupied regions (GORs) proximal to the glucocorticoid-responsive gene *FKBP5* experience distinct regulatory consequences to *FKBP5* regulation, correlated to differential loss and/or gain of binding sites for transcription factor binding site motif match(es).

In one example, a mutant with bi-allelic deletions at GOR -*26.65 kb* appeared unaffected for *FKBP5* induction at 1 nM and 100 nM dexamethasone (**Fig. 5c**, **clone #1**), whereas another mutant homozygous for a single-bp insertion (+1 bp) at the same GOR showed nearly ablated *FKBP5* induction (**Fig. 5c**, **clone #2**); closer analysis revealed that the mutant with ablated *FKBP5* induction harbored an apparent gain-of-function (increased pre-dex *FKBP5* transcript levels relative to wild-type), attributed to homozygous +1 bp insertion (**Fig. 5d**). Further evaluation with CollatedMotifs.py revealed that although both clone #1 and #2 lost the native GOR summit GBS as a consequence of Cas9 editing (**Fig. 5e**, maroon arrows (‘lost GBS’) in top (clone #1) and lower (clone #2) panels), the homozygous insertion in clone #2 uniquely reconstituted a novel GBS (**Fig. 5e**, red arrow (‘new GBS’) in lower panel). Moreover, clone #2 acquired sequence matches to Sox2 and DMRT3 binding motifs consequent to the edit within this GOR (**Fig. 5e**, lower panel). Identification of the ‘new Sox2’ site (**Fig. 5e**, blue arrow in lower panel) associated with an *FKBP5* transcriptional phenotype is particularly interesting, as Sox2—a TF typically associated with stemness—is ectopically expressed in many lung carcinomas, including A549^33, 34^. These results highlight that different Cas9 edits can be associated with distinct transcription regulatory consequences, potentially illuminated by mutation-specific TFBS alterations that could render distinct functionalities (*e.g.*, amorphic, neomorphic or inconsequential outcomes) to response elements under evaluation. Similarly, in other examples, small indels that successfully ablated the GBS native to the GOR summit commonly introduced a novel alternative GBS, potentially obviating the utility of the mutant clone for interpretation of regulatory consequences (**Supp. Fig. 13, Supp. Fig. 14**).

These results underscore the value of routine monitoring of altered regulatory motifs within candidate response elements indel-edited by Cas9, as indels engender distinct binding site alterations that may relate to distinct regulatory outcomes. Automated collation of lost and gained TFBS may be generally useful to inform selection of clones for analysis and/or to guide hypotheses for further study.

## Test data availability

The list of sample:barcode assignments, fastq files, GRCh38 reference genome, fasta file of reference sequences, and TFBS file with position frequency matrices underlying these examples are available at Zenodo (10.5281/zenodo.3406862) as test data, along with sample output files; users can recapitulate generation of the Sample Sheet that demultiplexed reads from these clones as a test of SampleSheet.py, and for a subset of fastq files (corresponding to Cas9 edits targeted to GOR+86.85 kb) can recapitulate generation of the imputed genotypes and associated files as a test of ImputedGenotypes.py, and can recapitulate comparison of TFBS motifs as a test of CollatedMotifs.py.

## Conclusions

### Other applications

We have described three Python programs (https://github.com/YamamotoLabUCSF), which will be of value for researchers who prepare amplicons for targeted SBS on Illumina® platforms: SampleSheet.py, ImputedGenotypes.py, CollatedMotifs.py. We presented the scripts in the context of a Cas9-editing workflow targeting candidate genomic response elements, but their uses are not limited to Cas9-editing scenarios; rather, they are applicable to any scenario calling for locus-specific assignment of allele definitions and genotype imputations to individual members of a potentially diverse population (for example, sequencing of single or multiple loci amplified from cell lines, tumor biopsies, cell-free DNA samples, viral passages, or individuals in a population). Illumina® SBS is broadly amenable to paired-end sequencing of amplicons that canvas alleles with larger indel size variation than those described here (**Supp. Fig. 15**). We envision that SampleSheet.py and the 96×96 i5/i7 barcoded primers may be of broadest utility to users who sequence pooled, PCR-amplified material from large populations of discrete entities and wish to back-track sequence properties to their sources; that ImputedGenotypes.py may be useful for those who need rapid distillation of allele definitions and hypothesized genotype(s) for samples of known biological origin (*i.e.*, with reference sequences available for alignment); and that CollatedMotifs.py may be of utility for users interested in an overview of TFBS differences resulting from genetic differences in experimental samples relative to a reference sequence, potentially aiding understanding of molecular phenotypes (hypothesis generation) or prioritization of clone choice/selection for further experimental analysis. Open source program files, annotated Jupyter notebooks, and Open Virtualization Format file for all code are available for download, enabling users to edit and tailor for customized goals, preferences and applications.

**Availability and requirements**

**Project name:** SampleSheet.py

**Project home page:** YamamotoLabUCSF/SampleSheet

https://github.com/YamamotoLabUCSF/SampleSheet

**Operating system(s):** Platform independent

**Programming Language:** Python

**Other requirements:** Python 3.7 or higher

Python package installations: PrettyTable

(CLI format; not required in Jupyter notebook format)

**License:** GNU General Public License

**Any restrictions to use by non-academics:** No restrictions

**Project name:** ImputedGenotypes.py

**Project home page:** YamamotoLabUCSF/ImputedGenotypes

https://github.com/YamamotoLabUCSF/ImputedGenotypes

**Operating system(s):** Platform independent

**Programming Language:** Python

**Other requirements:** Python 3 or higher, BLASTN

Installations: BLASTN

Python package installations: SciPy, NumPy, psutil, fpdf, PyPDF2

**License:** GNU General Public License

**Any restrictions to use by non-academics:** No restrictions

**Project name:** CollatedMotifs.py

**Project home page:** YamamotoLabUCSF/CollatedMotifs

https://github.com/YamamotoLabUCSF/CollatedMotifs

**Operating system(s):** Linux platform-dependent (Mac OS, virtual machine)

**Programming Language:** Python

**Other requirements:** Python 3 or higher, BLASTN, MAKEBLASTDB, FIMO, FASTA-GET-MARKOV

Installations: BLASTN, MAKEBLASTDB (BLAST+ suite);

FIMO, FASTA-GET-MARKOV (Meme suite)

Python package installations: SciPy, NumPy, psutil

**License:** GNU General Public License

**Any restrictions to use by non-academics:** No restrictions

## List of abbreviations

CLI: command-line interface
CRISPR: clustered regularly interspaced short palindromic repeats dex, dexamethasone
FACS: flow-assisted cell sorting
GBS: GR binding site
GC: glucocorticoid
GOR: GR-occupied region
GR: glucocorticoid receptor
GRE: glucocorticoid response element
i5/i7: generic Illumina® index primer identifiers
MCS: MiSeq Controller Software
NGS: next generation sequencing
PE: paired-end (dual indexed) sequencing
P5/P7: Illumina® flow cell oligonucleotide adapters
SBS: sequencing by synthesis
SE: single-end (single indexed) sequencing
TF: transcription factor
TFBS: transcription factor binding site

## Declarations

**Ethics approval and consent to participate:** Not applicable.

**Consent for publication:** Not applicable.

## Availability of data and material

Sample datasets analyzed in the current study are available under the following DOI: 10.5281/zenodo.3406862

## Authors’ contributions

KTE authored all code and deposited resources in GitHub and Zenodo repositories, performed Cas9 mutagenesis experiments and regulatory analysis, generated and analyzed sequencing datasets, and authored manuscript. MTK tested code, suggested updates for improvement, and assembled the Open Virtualization Format file (Linux virtual machine). DM performed GR ChIP-seq and peak calling analysis. MA tested code and documented amplicon size-dependent recoveries by two DNA clean-up approaches. HA tested amplicon length recoveries. KRY secured funding support, advised, and edited manuscript. All authors read and approved the final manuscript.

## Acknowledgments

We thank Mark Stenglein (Colorado State University, Fort Collins, CO) and Joseph DeRisi (University of California, San Francisco) for design and validation of 96×96 read1 and read2 barcode adaptors and sequences; all members of the Yamamoto laboratory for support, discussion and feedback; Sarah Elmes (Laboratory of Cell Analysis, UCSF; NIH P30CA082103) for expert FACS assistance and training; Jason Fernandes for initially introducing and sharing barcoded adaptors and sequences; Michel Tassetto for MiSeq background and Raul Andino lab (UCSF) for shared MiSeq use; Illumina® Technical Support for expert equipment troubleshooting and maintenance; Albertas Navickas and Hani Goodarzi lab (UCSF) for 384-well qPCR access; City College of San Francisco, in particular Douglas Putnam, Greg Boyd, Aaron Brick, and Jonathan Potter for programming coursework; contributors to Stack Exchange, Stack Overflow and related forums for freely sharing programming practices. Research support was from NIH CA020535 (to KRY).

