## Supplemental Material for "Definition of alleles and altered regulatory motifs across Cas9-edited cell populations"

#### Supplementary Figures

#### S1. 96x96 array of Illumina® sequencing platform-compatible i7 and i5 barcode primers.

(a) 192 primers (ninety-six "i7/index 1" and ninety-six "i5/index 2") are easily ordered in 96-well plate format from commercial oligonucleotide suppliers (*e.g.,* IDT, Coralville, IA), and can be used to label amplicons in up to 9,216 unique combinations for dual-indexed (paired-end) sequencing;

(b) up to 9,216 samples prepared for 96x96 dual barcoding can be arrayed in up to ninety-six 96-well plates or up to twenty-four 384-well plates (for example, see **Supp. Fig. 7**);

(c) i7 and i5 barcode sequences identified according to position in a 96-well plate when ordered from the .csv template provided (**Supp. Excel file**); 8-bp barcode sequence indicated 5'→3' as it appears in the primer (i7 or i5), or 5'→3' as it appears in the reverse complement (rev comp) frequently recovered as the index sequence in an Illumina® sequencing workflow.

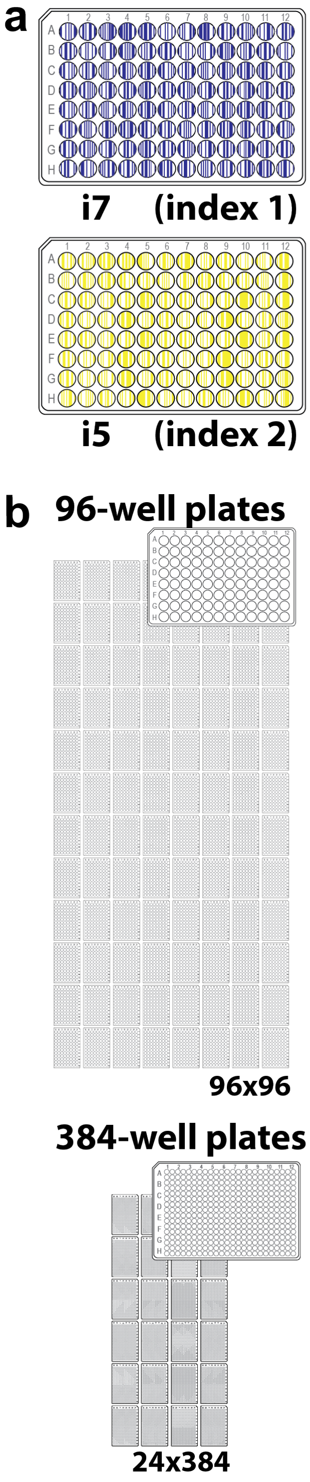


c

| i7 | | | | i5 | | | |
|---|---|---|---|---|---|---|---|
| well | name | i7 | rev. comp. | well | name | i5 | rev. comp. |
| A1 | read2_bc_1_TGACGTAC | TGACGTAC | GTACGTCA | A1 | read1_bc_1_GAGGTAGT | GAGGTAGT | CTCCATCA |
| A2 | read2_bc_2_TAACTGCA | TAACTGCA | TGCAGTTA | A2 | read1_bc_2_GCTTAACT | GCTTAACT | CGAATTGA |
| A3 | read2_bc_3_TCCACAGT | TCCACAGT | ACTGTGGA | A3 | read1_bc_3_GCAATTCT | GCAATTCT | CGTTAAGA |
| A4 | read2_bc_4_TCTTAACC | TCTTAACC | GGTTAAGA | A4 | read1_bc_4_TCCTCACT | TCCTCACT | AGGAGTGA |
| A5 | read2_bc_5_TAGTGTGA | TAGTGTGA | TCACACTA | A5 | read1_bc_5_AGTTAGCT | AGTTAGCT | TCAATCGA |
| A6 | read2_bc_6_TACCTCTA | TACCTCTA | TAGAGGTA | A6 | read1_bc_6_TCATGGCT | TCATGGCT | AGTACCGA |
| A7 | read2_bc_7_TATGTCGC | TATGTCGC | GCGACATA | A7 | read1_bc_7_AGCCTAAT | AGCCTAAT | TCGGATTA |
| A8 | read2_bc_8_TGCATGTA | TGCATGTA | TACATGCA | A8 | read1_bc_8_AACACTCT | AACACTCT | TTGTGAGA |
| A9 | read2_bc_9_TCATCATC | TCATCATC | GATGATGA | A9 | read1_bc_9_AGAGCTCT | AGAGCTCT | TCTCGAGA |
| A10 | read2_bc_10_TGCACACA | TGCACACA | TGTGTGCA | A10 | read1_bc_10_GGCCATAT | GGCCATAT | CCGGTATA |
| A11 | read2_bc_11_TGTAGCGA | TGTAGCGA | TCGCTACA | A11 | read1_bc_11_TAGAACGT | TAGAACGT | ATCTTGCA |
| A12 | read2_bc_12_TCTAGCTT | TCTAGCTT | AAGCTAGA | A12 | read1_bc_12_GGTTGAAT | GGTTGAAT | CCAACTTA |
| B1 | read2_bc_13_TGGTCCTA | TGGTCCTA | TAGGACCA | B1 | read1_bc_13_GATCCTAT | GATCCTAT | CTAGGATA |
| B2 | read2_bc_14_TCCAACGA | TCCAACGA | TCGTTGGA | B2 | read1_bc_14_TGGTCTCT | TGGTCTCT | ACCAGAGA |
| B3 | read2_bc_15_TAGGACAC | TAGGACAC | GTGTCCTA | B3 | read1_bc_15_TCCAGTCT | TCCAGTCT | AGGTCAGA |
| B4 | read2_bc_16_TATACGGA | TATACGGA | TCCGTATA | B4 | read1_bc_16_TAGCGTAT | TAGCGTAT | ATCGCATA |
| B5 | read2_bc_17_TAAGCGTA | TAAGCGTA | TACGCTTA | B5 | read1_bc_17_GCGATCAT | GCGATCAT | CGCTAGTA |
| B6 | read2_bc_18_TCGTTGTA | TCGTTGTA | TACAACGA | B6 | read1_bc_18_AGCTACAT | AGCTACAT | TCGATGTA |
| B7 | read2_bc_19_TCTCCTTC | TCTCCTTC | GAAGGAGA | B7 | read1_bc_19_ACTCATCT | ACTCATCT | TGAGTAGA |
| B8 | read2_bc_20_TGATGCTT | TGATGCTT | AAGCATCA | B8 | read1_bc_20_TTGCGAGT | TTGCGAGT | AACGCTCA |
| B9 | read2_bc_21_TGATCTCA | TGATCTCA | TGAGATCA | B9 | read1_bc_21_AAGCGACT | AAGCGACT | TTCGCTGA |
| B10 | read2_bc_22_TGTCTAGA | TGTCTAGA | TCTAGACA | B10 | read1_bc_22_TGCACGAT | TGCACGAT | ACGTGCTA |
| B11 | read2_bc_23_TATGAGCA | TATGAGCA | TGCTCATA | B11 | read1_bc_23_TATGCGGT | TATGCGGT | ATACGCCA |
| B12 | read2_bc_24_TGGAGTTA | TGGAGTTA | TAACTCCA | B12 | read1_bc_24_TCGTCGAT | TCGTCGAT | AGCAGCTA |
| C1 | read2_bc_25_TGTGTGTC | TGTGTGTC | GACACACA | C1 | read1_bc_25_ACACTTGT | ACACTTGT | TGTGAACA |
| C2 | read2_bc_26_TGCTACAA | TGCTACAA | TTGTAGCA | C2 | read1_bc_26_TACGGCAT | TACGGCAT | ATGCCGTA |
| C3 | read2_bc_27_TCGTAGAC | TCGTAGAC | GTCTACGA | C3 | read1_bc_27_TCTCGTGT | TCTCGTGT | AGAGCACA |
| C4 | read2_bc_28_TGTAATGC | TGTAATGC | GCATTACA | C4 | read1_bc_28_GCAGATGT | GCAGATGT | CGTCTACA |
| C5 | read2_bc_29_TATGGAAC | TATGGAAC | GTTCCATA | C5 | read1_bc_29_TCCGTTGT | TCCGTTGT | AGGCAACA |
| C6 | read2_bc_30_TGGTATTC | TGGTATTC | GAATACCA | C6 | read1_bc_30_TTCGCAGT | TTCGCAGT | AAGCGTCA |
| C7 | read2_bc_31_TCAAGTGA | TCAAGTGA | TCACTTGA | C7 | read1_bc_31_GGAATAGT | GGAATAGT | CCTTATCA |
| C8 | read2_bc_32_TCTTGTGT | TCTTGTGT | ACACAAGA | C8 | read1_bc_32_GCAACAAT | GCAACAAT | CGTTGTTA |
| C9 | read2_bc_33_TCTGTAGT | TCTGTAGT | ACTACAGA | C9 | read1_bc_33_ACTGCGAT | ACTGCGAT | TGACGCTA |
| C10 | read2_bc_34_TCCTGAGA | TCCTGAGA | TCTCAGGA | C10 | read1_bc_34_TGAGTACT | TGAGTACT | ACTCATGA |
| C11 | read2_bc_35_TGCACGAT | TGCACGAT | ATCGTGCA | C11 | read1_bc_35_ACACGAAT | ACACGAAT | TGTGCTTA |
| C12 | read2_bc_36_TACGTGGA | TACGTGGA | TCCACGTA | C12 | read1_bc_36_GTGGCAAT | GTGGCAAT | CACCGTTA |
| D1 | read2_bc_37_TCATGGAT | TCATGGAT | ATCCATGA | D1 | read1_bc_37_GTCATCCT | GTCATCCT | CAGTAGGA |
| D2 | read2_bc_38_TCTGCGAA | TCTGCGAA | TTCGCAGA | D2 | read1_bc_38_ACCGTACT | ACCGTACT | TGGCATGA |
| D3 | read2_bc_39_TGAGACGA | TGAGACGA | TCGTCTCA | D3 | read1_bc_39_GTACTACT | GTACTACT | CATGATGA |
| D4 | read2_bc_40_TCGTAAGT | TCGTAAGT | ACTTACGA | D4 | read1_bc_40_TAGGTCCT | TAGGTCCT | ATCCAGGA |
| D5 | read2_bc_41_TAATCCGA | TAATCCGA | TCGGATTA | D5 | read1_bc_41_GTTACTGT | GTTACTGT | CAATGACA |
| D6 | read2_bc_42_TAGACCAA | TAGACCAA | TTGGTCTA | D6 | read1_bc_42_GCCTTAGT | GCCTTAGT | CGGAATCA |
| D7 | read2_bc_43_TCCGCTAT | TCCGCTAT | ATAGCGGA | D7 | read1_bc_43_AACCATGT | AACCATGT | TTGGTACA |
| D8 | read2_bc_44_TGGTTACA | TGGTTACA | TGTAACCA | D8 | read1_bc_44_GATGACAT | GATGACAT | CTACTGTA |
| D9 | read2_bc_45_TAGCCTGT | TAGCCTGT | ACAGGCTA | D9 | read1_bc_45_AGAAGACT | AGAAGACT | TCTTCTGA |
| D10 | read2_bc_46_TGCGACTT | TGCGACTT | AAGTCGCA | D10 | read1_bc_46_TCAAGCAT | TCAAGCAT | AGTTCGTA |
| D11 | read2_bc_47_TCGAACTC | TCGAACTC | GAGTTCGA | D11 | read1_bc_47_TGGATGCT | TGGATGCT | ACCTACGA |
| D12 | read2_bc_48_TGAAGAGT | TGAAGAGT | ACTCTTCA | D12 | read1_bc_48_ACTTGCCT | ACTTGCCT | TGAACGGA |
| E1 | read2_bc_49_TAGGTCTT | TAGGTCTT | AAGACCTA | E1 | read1_bc_49_GTTGTGAT | GTTGTGAT | CAACACTA |
| E2 | read2_bc_50_TGGATGGT | TGGATGGT | ACCATCCA | E2 | read1_bc_50_TATCTCGT | TATCTCGT | ATAGAGCA |
| E3 | read2_bc_51_TACCGTTC | TACCGTTC | GAACGGTA | E3 | read1_bc_51_TGTAAGGT | TGTAAGGT | ACATTCCA |
| E4 | read2_bc_52_TCTCACTA | TCTCACTA | TAGTGAGA | E4 | read1_bc_52_ACGTTGGT | ACGTTGGT | TGCAACCA |
| E5 | read2_bc_53_TGTTGGAC | TGTTGGAC | GTCCAACA | E5 | read1_bc_53_AGGAGGAT | AGGAGGAT | TCCTCCTA |
| E6 | read2_bc_54_TCGCCATA | TCGCCATA | TATGGCGA | E6 | read1_bc_54_GAGCAGAT | GAGCAGAT | CTCGTCTA |
| E7 | read2_bc_55_TGATAAGC | TGATAAGC | GCTTATCA | E7 | read1_bc_55_TCTCTACT | TCTCTACT | AGAGATGA |
| E8 | read2_bc_56_TCAGTCAC | TCAGTCAC | GTGACTGA | E8 | read1_bc_56_GGATTCAT | GGATTCAT | CCTAAGTA |
| E9 | read2_bc_57_TATCACGT | TATCACGT | ACGTGATA | E9 | read1_bc_57_GACCTTCT | GACCTTCT | CTGGAAGA |
| E10 | read2_bc_58_TAAGAACC | TAAGAACC | GGTTCTTA | E10 | read1_bc_58_ACGGTTAT | ACGGTTAT | TGCCAATA |
| E11 | read2_bc_59_TGCTGATC | TGCTGATC | GATCAGCA | E11 | read1_bc_59_AGATGCGT | AGATGCGT | TCTACGCA |
| E12 | read2_bc_60_TATTGTCC | TATTGTCC | GGACAATA | E12 | read1_bc_60_TGACACCT | TGACACCT | ACTGTGGA |
| F1 | read2_bc_61_TCAGAGTT | TCAGAGTT | AACTCTGA | F1 | read1_bc_61_ACTAACGT | ACTAACGT | TGATTGCA |
| F2 | read2_bc_62_TAAGGTCA | TAAGGTCA | TGACCTTA | F2 | read1_bc_62_GAATGCCT | GAATGCCT | CTTACGGA |
| F3 | read2_bc_63_TCACTTCC | TCACTTCC | GGAAGTGA | F3 | read1_bc_63_GTGTACGT | GTGTACGT | CACATGCA |
| F4 | read2_bc_64_TCCTTCTT | TCCTTCTT | AAGAAGGA | F4 | read1_bc_64_AAGTCAGT | AAGTCAGT | TTCAGTCA |
| F5 | read2_bc_65_TAGATTGC | TAGATTGC | GCAATCTA | F5 | read1_bc_65_TTCCTGGT | TTCCTGGT | AAGGACCA |
| F6 | read2_bc_66_TGACTCGT | TGACTCGT | ACGAGTCA | F6 | read1_bc_66_ACCACCAT | ACCACCAT | TGGTGGTA |
| F7 | read2_bc_67_TGGCATAA | TGGCATAA | TTATGCCA | F7 | read1_bc_67_GAGTCCAT | GAGTCCAT | CTCAGGTA |
| F8 | read2_bc_68_TCTATGGC | TCTATGGC | GCCATAGA | F8 | read1_bc_68_AGCTCTGT | AGCTCTGT | TCGAGACA |
| F9 | read2_bc_69_TCCGATTA | TCCGATTA | TAATCGGA | F9 | read1_bc_69_GGTCTTGT | GGTCTTGT | CCAGAACA |
| F10 | read2_bc_70_TAACGAGA | TAACGAGA | TCTCGTTA | F10 | read1_bc_70_AATGTGCT | AATGTGCT | TTACACGA |
| F11 | read2_bc_71_TACAGACC | TACAGACC | GGTCTGTA | F11 | read1_bc_71_GTCTGGAT | GTCTGGAT | CAGACCTA |
| F12 | read2_bc_72_TACCGGAT | TACCGGAT | ATCCGGTA | F12 | read1_bc_72_GCGTATAT | GCGTATAT | CGCATATA |
| G1 | read2_bc_73_TGTCGCAT | TGTCGCAT | ATGCGACA | G1 | read1_bc_73_TGACTGAT | TGACTGAT | ACTGACTA |
| G2 | read2_bc_74_TGCGTTCA | TGCGTTCA | TGAACGCA | G2 | read1_bc_74_TGTCCAGT | TGTCCAGT | ACAGGTCA |
| G3 | read2_bc_75_TCATACCA | TCATACCA | TGGTATGA | G3 | read1_bc_75_TTGTGCCT | TTGTGCCT | AACACGGA |
| G4 | read2_bc_76_TCTGTCCA | TCTGTCCA | TGGACAGA | G4 | read1_bc_76_ACAGACCT | ACAGACCT | TGTCTGGA |
| G5 | read2_bc_77_TCTTGCAA | TCTTGCAA | TTGCAAGA | G5 | read1_bc_77_GATCAAGT | GATCAAGT | CTAGTTCA |
| G6 | read2_bc_78_TACCTAGT | TACCTAGT | ACTAGGTA | G6 | read1_bc_78_GATAGGAT | GATAGGAT | CTATCCTA |
| G7 | read2_bc_79_TATCGGTA | TATCGGTA | TACCGATA | G7 | read1_bc_79_GTAGTCGT | GTAGTCGT | CATCAGCA |
| G8 | read2_bc_80_TCACCAAT | TCACCAAT | ATTGGTGA | G8 | read1_bc_80_TCACCTAT | TCACCTAT | AGTGGATA |
| G9 | read2_bc_81_TCGAGGAA | TCGAGGAA | TTCCTCGA | G9 | read1_bc_81_AGTGGTAT | AGTGGTAT | TCACCATA |
| G10 | read2_bc_82_TACCATGA | TACCATGA | TCATGGTA | G10 | read1_bc_82_GCTAGAGT | GCTAGAGT | CGATCTCA |
| G11 | read2_bc_83_TGTGGATA | TGTGGATA | TATCCACA | G11 | read1_bc_83_TTAACGCT | TTAACGCT | AATTGCGA |
| G12 | read2_bc_84_TACAAGTC | TACAAGTC | GACTTGTA | G12 | read1_bc_84_TCAGGAGT | TCAGGAGT | AGTCCTCA |
| H1 | read2_bc_85_TACTCCAT | TACTCCAT | ATGGAGTA | H1 | read1_bc_85_AACTGGCT | AACTGGCT | TTGACCGA |
| H2 | read2_bc_86_TATCTGAC | TATCTGAC | GTCAGATA | H2 | read1_bc_86_TACAGAGT | TACAGAGT | ATGTCTCA |
| H3 | read2_bc_87_TAGTGCTC | TAGTGCTC | GAGCACTA | H3 | read1_bc_87_TTGCATCT | TTGCATCT | AACGTAGA |
| H4 | read2_bc_88_TGAATGAC | TGAATGAC | GTCATTCA | H4 | read1_bc_88_GAACGTGT | GAACGTGT | CTTGCACA |
| H5 | read2_bc_89_TCTTAGGA | TCTTAGGA | TCCTAAGA | H5 | read1_bc_89_AAGTACCT | AAGTACCT | TTCATGGA |
| H6 | read2_bc_90_TACGCATC | TACGCATC | GATGCGTA | H6 | read1_bc_90_GGTACCAT | GGTACCAT | CCATGGTA |
| H7 | read2_bc_91_TGGAAGCA | TGGAAGCA | TGCTTCCA | H7 | read1_bc_91_AATCGCAT | AATCGCAT | TTAGCGTA |
| H8 | read2_bc_92_TCTGACAT | TCTGACAT | ATGTCAGA | H8 | read1_bc_92_GGTAATCT | GGTAATCT | CCATTAGA |
| H9 | read2_bc_93_TGAGCAAC | TGAGCAAC | GTTGCTCA | H9 | read1_bc_93_GATTCGCT | GATTCGCT | CTAAGCGA |
| H10 | read2_bc_94_TGGATATC | TGGATATC | GATATCCA | H10 | read1_bc_94_AGCATCGT | AGCATCGT | TCGTAGCA |
| H11 | read2_bc_95_TCGCTAAC | TCGCTAAC | GTTAGCGA | H11 | read1_bc_95_GTTCACCT | GTTCACCT | CAAGTGGA |
| H12 | read2_bc_96_TGTACCTC | TGTACCTC | GAGGTACA | H12 | read1_bc_96_AGGCAAGT | AGGCAAGT | TCCGTTCA |

**S2. Overview of customized Illumina® Sample Sheet sections populated by SampleSheet.py.**

The Illumina® Sample Sheet is indispensable for accurate demultiplexing of pooled reads to individual, sample-specific fastq files. Four sections of the Illumina® Sample Sheet are customized by user input at the SampleSheet.py interface: *InvestigatorName, ProjectName, [Reads]* (single-end/SE or paired-end/PE; cycle #(s) for read1, read2), and contents of the *[Data]* table. *Date* is auto-generated using system's current calendar date.

Should users wish to customize default key values, Illumina® publications can be consulted for further details and metadata key:value options; default parameters may be readily customized by editing the appropriate field content of the SampleSheet *.csv file output, or contents of the script itself (*e.g.*, Jupyter notebook section labeled, 'IV. Create Sample Sheet and populate with content').

**[Header]**
InvestigatorName, ***user-supplied***
ProjectName, ***user-supplied***
Date, ***auto-generated***
Workflow, GenerateFASTQ
Application, FASTQonly
Description, Sequencing
Chemistry, Amplicon

**[Reads]**
***user-supplied***

**[Settings]**
ReverseComplement, 0
Adapter, CTGTCTCTTATACACATCT

**[Data]**
Sample_ID, Sample_Name, i7_Index_ID, index, i5_Index_ID, index2
***user-supplied***

**S3. Command-line and Jupyter notebook interface (console) views of user prompts**
*(examples shown are from Jupyter notebook interface)*
**(a) SampleSheet.py;**
**(b) ImputedGenotypes.py;**
**(c) CollatedMotifs.py.**

### a SampleSheet.py

## 1

```
--------------------------------------------
Illumina Indexed Sequencing Workflow (A or B)
--------------------------------------------

Illumina Indexed Sequencing for dual-indexed (Paired End) runs uses one of two different Workflows (A or B),
defined by whether (A) both index sequences (i7 and i5) are sequenced by primers that anneal to the 'Read 1' strand,
or (B) i7 index is sequenced by a primer that anneals to the 'Read 1' strand, whereas i5 index is sequenced by a
primer that anneals to the 'Read 2' strand. Awareness of the different Workflows for sequencing the indices is
critical, because Workflow determines whether the 5'->3' index sequence (i5 or i7) is returned as a 'forward'
sequence that reads just like the index sequence as it occurs in indexing primers during library preparation,
or whether the 5'->3' index sequence is returned as a 'reverse complement' relative to the index sequence as it
occurs in indexing primers during library preparation. Fundamentally, Workflow determines the 5'->3' nucleotide
sequences entered for i7 and i5 indices in an Illumina Sample Sheet, essential for faithful demultiplexing of
samples.

As of June 2019, Illumina sequencing instruments use the two dual-indexed Workflows as follows:

    Workflow A: MiSeq, NovaSeq 6000, HiSeq 2500, HiSeq 2000
    -------------------------------------------------------
    --> i7 index sequence is recovered as 'reverse complement' relative to i7 sequence
        as it occurs in primers used for library construction.
    --> i5 index sequence is recovered as it occurs in primers used for library construction.

    Workflow B: iSeq100, MiniSeq, NextSeq, HiSeqX, HiSeq 4000, HiSeq 3000
    ----------------------------------------------------------------------
    --> i7 index sequence is recovered as 'reverse complement' relative to i7 sequence
        as it occurs in primers used for library construction.
    --> i5 index sequence is recovered as 'reverse complement' relative to i5 sequence
        as it occurs in primers used for library construction.

    Please confirm the Workflow appropriate for your sequencing application.

    Note, if your application is strictly single-indexed (Single End run using only i7 indices), choose Workflow 'A'.

    Enter 'A' or 'B' to specify the Workflow, and therefore the index sequence orientations,
    appropriate for your Sample Sheet:
```

## 2

```
-----------------------------------------------------------------------------
Sample Sheet file name and location (absolute path to future .csv filename)
-----------------------------------------------------------------------------

Enter the name of the .csv file you'd like to create as your Sample Sheet, with an absolute path to its location.

The .csv file should not exist yet -- it will be created as an output of this script.

To create the .csv file in a directory where you can find it, enter an *absolute path*
to where you would like this file to be created.

Example: if your target file name is 'SampleSheet.csv', and you'd like to create that file
in a directory that is accessed with an absolute path of /Users/myname/Illumina/SampleSheet.csv,
enter '/Users/myname/Illumina/SampleSheet.csv' (Mac) or 'C:\Users\myname\Illumina\SampleSheet.csv' (PC)
at the command line prompt. Replace 'myname' with the appropriate intervening directory identifiers.
Do *not* flank your entry with quotation marks (') at the command-line.

Alternatively, simply enter a target file name (e.g., 'SampleSheet.csv') and run this script from
within a directory where you'd like to output this file.

     -----> File name and path:
```

#### a SampleSheet.py *(continued)*

**3**

```
----------------------------------------------------------------
Sample Sheet inputs: [Header], [Reads], and [Data] specifications
----------------------------------------------------------------

.......................................................
[Header] details: specify InvestigatorName & ProjectName

To specify InvestigatorName & ProjectName, enter text for each directly at the command line,
separated by a comma ('InvestigatorName, ProjectName').

When both text entries are entered, press ‘enter’ again to proceed in the script.
To skip text entries for these fields, simply press ‘enter’ until the next prompt appears (if you skip
entry now, 'NA' will be entered in these fields in the output file).

Example: if your InvestigatorName is 'Dorothy Gale' and ProjectName is 'Sequences', enter
'Dorothy Gale, Sequences'.


    -----> [Header] details. InvestigatorName & ProjectName:
```

**4**

```
..........................................................................................
[Reads] details: specify whether sequencing is Single-End or Paired-End, and the number of cycles

To specify Single-End vs. Paired-End format and number of cycles, enter text directly at the command line,
on a single line. Indicate Single-End (SE) or Paired-End (PE), followed by the number of cycles for each read.
Separate values by comma(s).

When text is entered, press ‘enter’ again to proceed in the script.

Examples:
If you are performing a Paired-End run with 151 cycles in reads 1 & 2, enter
'PE, 151, 151'.

If you are performing a Single-End run with 151 cycles in read 1, enter
'SE, 151'.
```

**5**

```
......................................................................
[Data] details: specify relationships between sample names and barcodes.

You will now be asked to enter a text-only table of plate names with their corresponding
i7 barcode range and i5 barcode, for dual-indexed (Paired-End) sequencing.

Specifics:
    * Each line corresponds to samples barcoded across a single 96-well plate.
    * Each line is expanded in the output Sample Sheet based on the 'plate name' and specified number of
        barcoded sample wells; individual wells are assigned unique sample IDs in the Sample Sheet based on their
        unique combination of 'plate name' and 'i7 barcode' well ID.
        * 'Plate name' identifies a single 96-well plate identifier and must be unique.
            * Any letter, digit, and punctuation characters are acceptable in names,
              excluding underscores ('_') and semicolons (';'), which must *not* be used.
        * 'i7 barcode' identifies an individual well
          (entry will be a numeric range, any # range up to '1-96')
        * 'i5 barcode' identifies all wells in a single plate
          (entry will be a single number, any # in '1' to '96'.)
        * For i7 and i5 barcode identifiers, use only the *range* (e.g., '1-96') or *number* (e.g., '1'-'96')
          that corresponds to a given index. Refer to i5 and i7 96-well plate sequences
          (displayed earlier as console PLATEVIEWS), if needed.
        * Fields (plate name, i7 index range, i5 index) are comma-separated.
        * You may manually enter or paste up to 96 lines that specify sample name-barcode relationships.
        * End each plate description line with a semi-colon (';'), except for the final line.

Example: imagine that you have 310 samples barcoded across four 96-well plates (some plates containing 96 samples,
some plates containing fewer than 96 samples). Plate 1 used unique i7 barcodes '1-96' ('i7A01-i7H12') across 96
samples + i5 barcode '4' ('i5A05') for all 96 samples; Plate 2 used the same i7 barcode range + i5 barcode '5'
('i5A05') for its 96 samples, Plate 3 used unique barcodes '1-50' ('i7A01-i7E02') across 50 samples + i5 barcode '2'
('i5A02') for all 50 samples; Plate 4 used unique barcodes '1-68' ('i7A01-i7F08') across 68 samples + i5 barcode '34'
('i5C10') for all 68 samples. You would enter text, line by line at the command line, that resembles this:

        DG-1, 1-96, 4;
        DG-2, 1-96, 5;
        DG-3, 1-50, 2;
        DG-4, 1-68, 34

When you're done entering plates and their indices, press 'Enter' again to proceed in the script.


    -----> [Data] details:
```

### b ImputedGenotypes.py

#### *List vs. Prompt*

```
-----------------------------------------------------------------------
User-specified input: choice of coached prompts vs. single-list entry
-----------------------------------------------------------------------

Values for the user-specified input indicated above can be entered at individually coached command-line prompts
(default), or as a single list of variables provided in a single command-line entry without coached prompts.

To proceed with input at individual command-line PROMPTS, type 'Prompt' and press Enter;
To proceed with input provided as a single LIST in one command-line entry, type 'List' and press Enter:
```

**1**

```
---------------------------------------------
Location of OUTPUT DIRECTORY for output files
---------------------------------------------

The script generates 8 separate files, all in the directory you indicate here.  It is important that this directory
either not exist prior to running the script, or if it does exist, it must be *empty* of any files with the names
to be created below.  These files are:

    1. fasta.fa

    2. blastn_alignments.txt
        (output of blastn operation on fasta.fa)

    3. allele_definitions.txt
        (output of script operation on blastn_alignments.txt, samples returned in order of processing)

    4. allele_evidence.pdf
        (output of script operation on blastn_alignments.txt, plot of calculated read/allele frequencies)

    5. imputed_genotypes.txt
        (output of script operation on blastn_alignments.txt, samples returned in order of genotype imputation)

    6. population_summary.txt
        (output of script operation on imputed_genotypes.txt)

    7. allele_definitions.csv
        (allele metrics (frequency representations) and definitions for each sample, in spreadsheet format)

    8. script_metrics.txt
        (summary/analysis of script operation metrics)

        Notes:
        * These files do not exist before the script is run. The files are made by the script.
        * The primary data outputs for genotypes are found in:
            allele_definitions.txt, allele_evidence.pdf, imputed_genotypes.txt & population_summary.txt


At this prompt, indicate an absolute path to a ** directory ** that will be created by the script as the location
for output files.  This directory should not exist yet -- it will be created as an output of this script, and will
be populated with the file outputs of this specific instance of the script operation.

Use only forward slashes ('/') as directory separators, regardless of operating system (Mac or Windows).

Example: if you'd like to create a directory ('ImputedGenotypes') in an existing directory ('Illumina'), accessed
with absolute path of '/Users/myname/Illumina/ImputedGenotypes' (Mac) or 'C:\Users\myname\Illumina\ImputedGenotypes'
(Windows), enter '/Users/myname/Illumina/ImputedGenotypes' at the command line prompt. Replace 'myname' with the
appropriate intervening directory identifiers. Do *not* flank your entry with quotation marks (') at the command
line.

Alternatively, simply enter a desired directory name (e.g., 'ImputedGenotypes') and run this script from
within a directory where you'd like to create this new directory.

    -----> Output directory name and path:
```

### b ImputedGenotypes.py *(continued)*

## 2

```
-----------------------------------------------------------------------------
Location of INPUT FILES (single directory containing demutiplexed fastq files)
-----------------------------------------------------------------------------

You will now be asked to enter the path to the directory containing the fastq files
to be processed as ImputedGenotypes.py input.

Use only forward slashes ('/') as directory separators.

Example: if your fastq input files are named file1.fastq, file2.fastq, etc. and are found in a directory
named 'Sequences' with absolute path of '/Users/myname/Sequences' (Mac) or 'C:\Users\myname\Sequences' (PC),
enter '/Users/myname/Sequences' at the command line prompt.

When you're done entering the fastq file location, press 'Enter' again to proceed in the script.


    -----> Directory name and path:
```

## 3

```
-----------------------------
Location of BLASTN EXECUTABLE
-----------------------------

This script uses BLASTN (NCBI) to align reads from your fastq files to a reference sequence database
(such as a genome database or sequence database).
Please indicate the absolute path to the BLASTN executable.

Use only forward slashes ('/') as directory separators.

Example: if your BLASTN executable is found at absolute path /Users/myname/blastn, type '/Users/myname/blastn'
and press Enter.


    -----> Path to BLASTN executable:
```

## 4

```
---------------------------------------------
Location of BLASTN ALIGNMENT DATABASE DIRECTORY
---------------------------------------------

Because this script uses BLASTN (NCBI) to align reads from your fastq files a reference sequence database,
an alignment reference database is needed. This reference database consists of a single directory containing
six files (.nhr, .nin, .nog, .nsd, .nsi, .nsg) (generated by the program MAKEBLASTDB (NCBI) from a file
containing sequences in fasta format, or downloaded from NCBI as an existing database).

Please indicate the absolute path to the directory you are using as your reference sequence database.

Use only forward slashes ('/') as directory separators.

Example: if your reference sequence database is found at absolute path /Users/myname/database, type
'/Users/myname/database' and press Enter.


    -----> Path to BLASTN alignment reference sequence database:
```

## 5

```
---------------------------------------------
PREFIX common to BLASTN ALIGNMENT DATABASE FILES
---------------------------------------------

A BLASTN reference sequence database consists of six files in a single directory, with each of the six
files sharing a common prefix (usually determined by the name of the fasta file provided to MAKEBLASTDB
during database generation).

Please indicate the common prefix for files of the reference sequence database.


    -----> Prefix for alignment reference sequence database files:
```

#### **b ImputedGenotypes.py** *(continued)*

```
----------------------------------------------------------------
Optional: Nucleotide sequence(s) to identify in output alignments
----------------------------------------------------------------

Some applications of 'allele definition' and 'genotype imputation' may call for identification of the presence
or absence of a specific anticipated sub-sequence (few nucleotides), and/or for the mapping of the location of
a sub-sequence if present in the sequence alignment.

ImputedGenotypes.py allows for the optional testing of sub-sequences.

If you would like to specify subsequences, type 'Yes' and press Enter.
Otherwise, if you do not wish to specify subsequences, type 'No' and press Enter.

    -----> 'Yes' or 'No' to sub-sequence specification:

    ----------------------------------------------------------------------------------------------------
    Nucleotide sequence(s): guide RNA annealing sites and/or test for presence/absence of sub-sequence
    ----------------------------------------------------------------------------------------------------
```

**6**
```
    .............................................................
    ***** guide RNA details: specify guide RNA sequence(s) *****

    To specify guide RNA sequence(s), enter text for each directly at the command line,
    separated by a comma ('x,y').

    Please specify guide RNA sequence(s) [excluding PAM]:

    When text entries are entered, press 'enter' again to proceed in the script.
    To skip text entries for these fields, simply press 'enter' until the next prompt appears.

    Examples:
        If your single guide RNA sequence is 'ATCCAGTTCTCCAGTCTCCC', enter: 'ATCCAGTTCTCCAGTCTCCC'.
        If you have two guide RNA sequences and they are 'ATCCAGTTCTCCAGTCTCCC' and 'GCGAGCTCGTGTCTGTGACG',
        enter: 'ATCCAGTTCTCCAGTCTCCC, GCGAGCTCGTGTCTGTGACG'.

        -----> guide RNA sequences:
```

**7**
```
    ..........................................................................................
    ***** query DNA sequence(s): specify sequence(s) to test for presence or absence *****

    To specify query DNA sequence(s), enter text for each directly at the command line,
    separated by a comma ('x,y').

    Please specify short DNA sequence to test for presence vs. ablation.

    When text entries are entered, press 'enter' again to proceed in the script.
    To skip text entries for these fields, simply press 'enter' until the next prompt appears.

    Examples:
        If your single query sequence is 'TACTCAATATCGATC', enter: 'TACTCAATATCGATC'.
        If you have two query sequences and they are 'TACTCAATATCGATC' and 'CGGGAGCCCGAG', enter:
        'TACTCAATATCGATC, CGGGAGCCCGAG'.

        -----> query DNA sequences:
```

### b ImputedGenotypes.py *(continued)*

#### *List vs. Prompt*

```
----------------------------------
User-specified input (list format)
----------------------------------
Please paste individual input values directly at the command line prompts, specifying the following 7 values
in the specified order.

Press 'Enter' twice to complete.

    1-Location of OUTPUT DIRECTORY for output files
    2-Location of INPUT FILES (directory containing fastq files)
    3-Location of BLASTN EXECUTABLE
    4-Location of BLASTN ALIGNMENT DATABASE DIRECTORY
    5-Prefix common to BLASTN sequence database files
    6-Optional guide RNA sequence(s) to identify in output alignments
    7-Optional sub-sequence(s) to identify in output alignments
```

#### *Include or Bypass frequency plot generation (allele_evidence.pdf)*

```
ImputedGenotypes.py is ready to process fastq files. Before script operations begin, please indicate whether visual
plots of allele frequencies should be rendered and delivered in an output file, allele_evidence.pdf.

Note that production of allele_evidence.pdf can require hours of processing time, although the output timing of
key text files with allele definitions and genotype imputations (e.g., allele_definitions.txt, imputed_genotypes.txt,
allele_definitions.csv, population_summary.txt) will not be affected.

To PROCEED with script operations that INCLUDE allele_evidence.pdf, type 'Y';

To BYPASS script operations that generate allele_evidence.pdf, type 'N':
```

### C CollatedMotifs.py

***List vs. Prompt***

```
-----> List or Prompt:
```

```
----------------------------------------------------------------------
User-specified input: choice of coached prompts vs. single list entry
----------------------------------------------------------------------

Values for the user-specified input indicated above can be entered at individually coached command-line prompts
(default), or as a single list of variables provided in a single command-line entry without coached prompts.

To proceed with input at individual command-line PROMPTS, type 'Prompt' and press Enter;
To proceed with input provided as a single LIST in one command-line entry, type 'List' and press Enter:
```

## 1

```
---------------------------------------------
Location of OUTPUT DIRECTORY for output files
---------------------------------------------

This script produces 5 output files in the user-specified output directory, plus three directories:
two directories and subsidiary files created by FIMO (fimo_out and fimo_out_ref) and one directory
and subsidiary files created by MAKEBLASTDB (alignment_database).

CollatedMotifs.py output files include:

    1. fasta.fa

    2. blastn_alignments.txt
        (output of BLASTN operation on fasta.fa)

    3. markov_background.txt
        (output of FASTA-GET-MARKOV operation on user-supplied fasta reference file)

    4. collated_TFBS.txt
        (output of script operation on FIMO-generated .tsv files in fimo_out and fimo_out_ref)

    5. script_metrics.txt (summary/analysis of script operation metrics [metadata])

        Note:
        * These files do not exist before the script is run. The files are made by the script.
        * The primary data outputs for TFBS comparisons are found in collated_TFBS.txt

At this prompt, indicate an absolute path to a ** directory ** that will be created by the script as the location
for output files.  This directory should not exist yet -- it will be created as an output of this script, and will
be populated with the file outputs of this specific instance of the script operation.

Example: if you'd like to create a directory ('CollatedMotifs') in an existing directory ('Illumina'), accessed
with absolute path of '/Users/myname/Illumina/CollatedMotifs' (Mac) or 'C:\Users\myname\Illumina\CollatedMotifs'
(Windows), enter '/Users/myname/Illumina/CollatedMotifs' at the command line prompt. Replace 'myname' with the
appropriate intervening directory identifiers. Do *not* flank your entry with quotation marks (') at the
command-line.

Alternatively, simply enter a desired directory name (e.g., 'CollatedMotifs') and run this script from
within a directory where you'd like to create this new directory.


    -----> Output directory name and path:
```

## 2

```
----------------------------------------------------------------------------
Location of INPUT FILES (single directory containing demutiplexed fastq files)
----------------------------------------------------------------------------

You will now be asked to enter the path to the directory containing the fastq files
to be processed as CollatedMotifs.py input.

Example: if your fastq input files are named file1.fastq, file2.fastq, etc. and are found in a directory
named 'Sequences' with absolute path of '/Users/myname/Sequences' (Mac) or 'C:\Users\myname\Sequences' (PC),
enter '/Users/myname/Sequences' at the command line prompt.

When you're done entering the fastq file location, press 'enter' again to proceed in the script.


    -----> Directory name and path:
```

#### C CollatedMotifs.py *(continued)*

### 3

```
----------------------------------------
Location of FIMO REFERENCE SEQUENCES FILE
----------------------------------------

This script aligns and compares your top sample read sequence(s) to a defined reference sequence,
as its basis for determining distinct vs. common TFBS motifs.
Please indicate the absolute path to a fasta file containing reference sequences.

Example: if you have three sample names relating to three different reference sequences, enter these
sequences in fasta format, saved in a single text file. Each fasta entry definition line (defline)
should be named such that the defline name matches a unique descriptor in fastq file names.

    >Sample1
    GATCGACTAGAGCGAGCATTCATCATATCACGAGTAGCATCGACGTGCACGATCGATCGTAGCTAGCTAGTCATGCATGCATGCTAGATTCGAGCATGCATGCTAC
    >Sample2
    AGTAGCTGTGATGCTAGTCATCTAGCTAGCAGCGTAGCTAGCGATCGATCTAGAGCCGATCGATCGAGCATCTAGCTATCAGCGGCGGGATCATCTATCTACGGG
    >Sample3
    CGATGCAGCGCGATCGAGCGCGATCGATATTAGCATGCGCAGCTAGCTAGCTGGCGATCGATGCATGCTAGCTGTGTCAGTCGACGATCACACGATCACACTGTGTG

When you're done entering the list of reference sequences, press 'enter' again to proceed in the script.
```

### 4

```
-----------------------------
Location of BLASTN EXECUTABLE
-----------------------------

This script uses BLASTN (NCBI) to align reads from your fastq files to a reference sequence database.
Please indicate the absolute path to the BLASTN executable.

Example: if your BLASTN executable is found at absolute path /Users/myname/blastn, type '/Users/myname
and press Enter.

    -----> Path to BLASTN executable:
```

### 5

```
---------------------------------
Location of MAKEBLASTDB EXECUTABLE
---------------------------------

Because this script uses BLASTN (NCBI) to align reads from your fastq files to a reference sequence database,
a compatible reference sequence database is required. This script uses MAKEBLASTDB (NCBI) to generate
a reference sequence database from the reference sequences in the fasta file you provided earlier.

Please indicate the absolute path to the MAKEBLASTDB executable.

Example: if your MAKEBLASTDB executable is found at absolute path /Users/myname/makeblastdb,
type '/Users/myname/makeblastdb' and press Enter.

    -----> Path to MAKEBLASTDB executable:
```

### 6

```
-------------------------------------------
Prefix for files in BLASTN ALIGNMENT DATABASE
-------------------------------------------

Because this script uses BLASTN (NCBI) and an alignment reference database, a common prefix identifier for the six
database files generated by MAKEBLASTDB is needed.

Please indicate a prefix to assign to each of the database files.

Example: if your alignment reference was generated by MAKEBLASTDB from a fasta file called hg38.fa,
the alignment database files will have been assigned the prefix 'hg38'; you would type 'hg38'
and press Enter.

    -----> Prefix for alignment reference sequence database files:
```

#### c CollatedMotifs.py *(continued)*

**7**

```
--------------------------
Location of FIMO EXECUTABLE
--------------------------

This script uses FIMO from the MEME suite of sequence analysis tools as its basis for determining distinct vs.
common TFBSs.
Please indicate the absolute path to the FIMO installation.

Example: if your FIMO executable is found at absolute path /Users/myname/fimo, type '/Users/myname/fimo'
and press Enter.

    -----> Path to FIMO executable:
```

**8**

```
----------------------------
Location of FIMO MOTIFS FILE
----------------------------

This script uses FIMO from the meme suite of sequence analysis tools as its basis for determining distinct vs.
common TFBS motifs.
Please indicate the absolute path to the FIMO motifs file (containing position frequency matrix/matrices).

When you're done entering the location of the motifs file, press Enter.

    -----> Path to FIMO motifs file:
```

**9**

```
--------------------------------------------
Location of FIMO FASTA-GET-MARKOV EXECUTABLE
--------------------------------------------

This script uses FIMO from the MEME suite of sequence analysis tools as its basis for determining distinct vs.
common TFBSs.
Please indicate an absolute path to the location of the FASTA-GET-MARKOV executable.

When you're done entering the location of the executable, press Enter.

    -----> Path to FASTA-GET-MARKOV executable:
```

**10**

```
-----------------------------------------------------------
Location of FIMO FASTA-GET-MARKOV BACKGROUND REFERENCE FILE
-----------------------------------------------------------

This script uses FIMO from the MEME suite of sequence analysis tools as its basis for determining distinct vs.
common TFBSs.
Please indicate an absolute path to the location of the fasta file you will use as your background reference
(on which FASTA-GET-MARKOV will operate to generate a markov background file).

When you're done entering the location of the reference sequence, press Enter.

    -----> Path to background reference file:
```

***List format***

```
---------------------------------
User-specified input (list format)
---------------------------------

Please paste individual input values directly at the interpreter prompts, specifying the following 10 values
in the specified order.

Press 'Enter' twice to complete.

    1-Location of OUTPUT DIRECTORY for output files
    2-Location of INPUT FILES (directory containing fastq files)
    3-Location of REFERENCE FASTA FILE
    4-Location of BLASTN EXECUTABLE
    5-Location of MAKEBLASTDB EXECUTABLE
    6-Prefix to assign to BLASTN sequence database files
    7-Location of FIMO EXECUTABLE
    8-Location of POSITION FREQUENCY MATRIX FILE
    9-Location of FASTA-GET-MARKOV EXECUTABLE
    10-Location of MARKOV BACKGROUND FILE
```

**S4. Illumina® Sequencing by Synthesis (SBS) workflows A & B: distinct orders of operation determine whether i7 & i5 indices are read in the same orientation as the sequenced read or in antiparallel fashion (reverse complement).**

Dual-indexed (paired-end) DNA sequencing on Illumina® instruments can use one of two Workflows (A or B), defined based on whether re-synthesis of complementary DNA template strands (for read 2) takes place before (*Workflow A)* or after (*Workflow B*) i5 (index 2) sequencing. Awareness of the Workflow customary to a sequencing application is important in Sample Sheet [Data] entry, because Workflow determines whether an index sequence is provided to demultiplexing software as the 'forward' sequence or its 'reverse complement'. In the MiSeq application reported as an Example Case Use (**Fig. 5**), we used the Illumina® MiSeq (MiSeq Reagent Kit v2: PE, 2x150 bp)—an instrument that applies *Workflow A*. In the Sample Sheet [Data] table, we therefore chose 'Workflow A' at the appropriate SampleSheet.py console prompt, and the script automatically populated i7 barcode sequences as *reverse complements* relative to barcode sequences as they appear in the i7 primers used in library construction, and i5 barcode sequences as *forward* orientations relative to barcode sequences as they appear in the i5 primers used in library construction (**Supp. Fig. 1c, Supp. Fig. 3b**).

(a) As of June 2019, Illumina® MiSeq, NovaSeq 6000, HiSeq 2500, and HiSeq 2000 instruments use Workflow A, where i7 index sequence is recovered as 'reverse complement' relative to i7 sequence as it occurs in primers used for library construction and i5 index sequence is recovered as it occurs in primers used for library construction;

(b) iSeq100, MiniSeq, NextSeq, HiSeqX, HiSeq 4000, and HiSeq 3000 instruments use Workflow B, where both i7 and i5 index sequences are recovered as 'reverse complements' relative to i7 and i5 sequences as they occur in primers used for library construction.
*See* ***Supp. Fig. 5b*** *for definitions of color-coded amplicon subregions.*

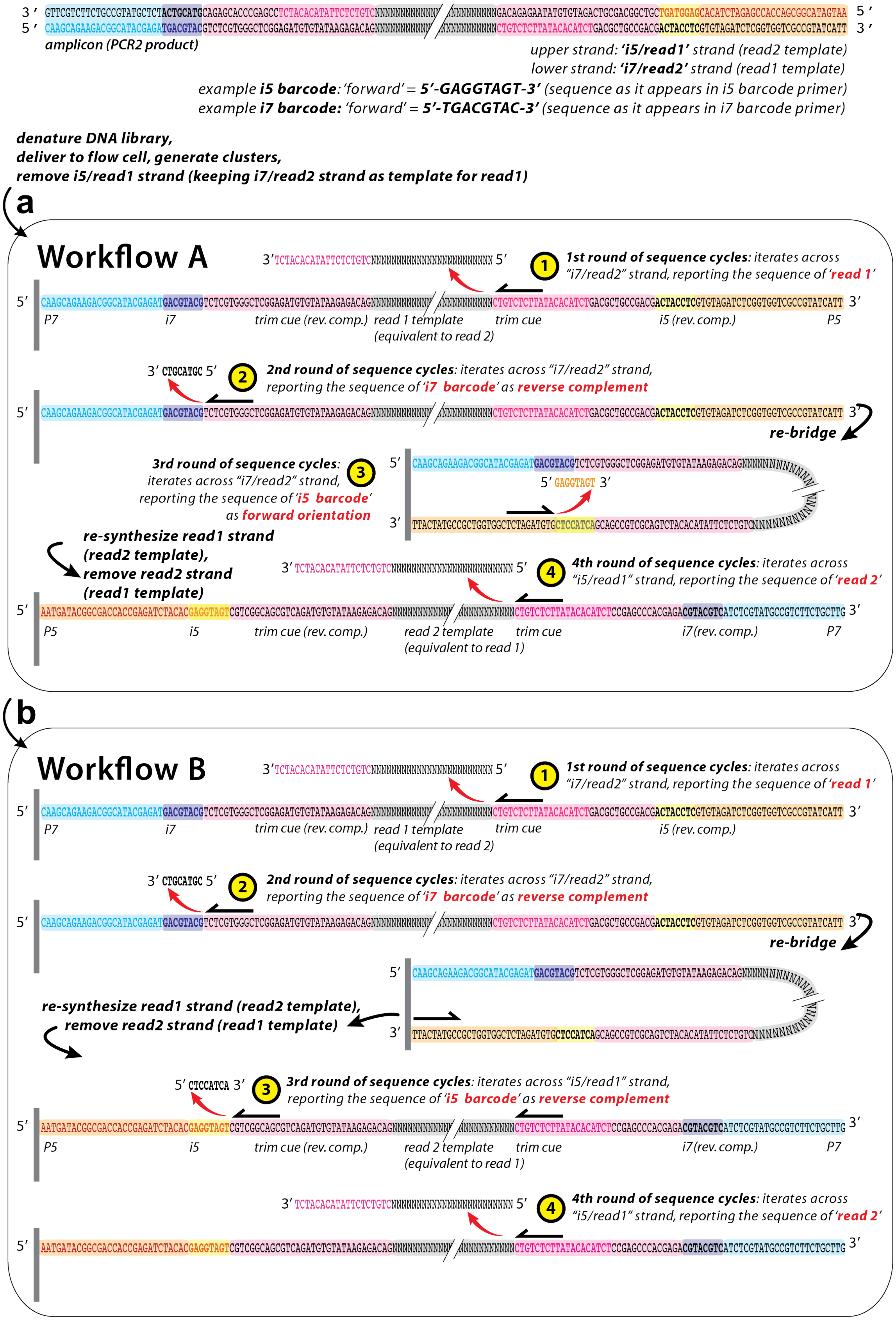
amplicon (PCR2 product)
upper strand: 'i5/read1' strand (read2 template)
lower strand: 'i7/read2' strand (read1 template)
example i5 barcode: 'forward' = 5'-GAGGTAGT-3' (sequence as it appears in i5 barcode primer)
example i7 barcode: 'forward' = 5'-TGACGTAC-3' (sequence as it appears in i7 barcode primer)
denature DNA library,
deliver to flow cell, generate clusters,
remove i5/read1 strand (keeping i7/read2 strand as template for read1)
a
Workflow A
1st round of sequence cycles: iterates across "i7/read2" strand, reporting the sequence of 'read 1'
P7
i7
trim cue (rev. comp.)
read 1 template (equivalent to read 2)
trim cue
i5 (rev. comp.)
P5
2nd round of sequence cycles: iterates across "i7/read2" strand, reporting the sequence of 'i7 barcode' as reverse complement
re-bridge
3rd round of sequence cycles: iterates across "i7/read2" strand, reporting the sequence of 'i5 barcode' as forward orientation
re-synthesize read1 strand (read2 template), remove read2 strand (read1 template)
4th round of sequence cycles: iterates across "i5/read1" strand, reporting the sequence of 'read 2'
P5
i5
trim cue (rev. comp.)
read 2 template (equivalent to read 1)
trim cue
i7 (rev. comp.)
P7
b
Workflow B
1st round of sequence cycles: iterates across "i7/read2" strand, reporting the sequence of 'read 1'
P7
i7
trim cue (rev. comp.)
read 1 template (equivalent to read 2)
trim cue
i5 (rev. comp.)
2nd round of sequence cycles: iterates across "i7/read2" strand, reporting the sequence of 'i7 barcode' as reverse complement
re-bridge
re-synthesize read1 strand (read2 template), remove read2 strand (read1 template)
3rd round of sequence cycles: iterates across "i5/read1" strand, reporting the sequence of 'i5 barcode' as reverse complement
P5
i5
trim cue (rev. comp.)
read 2 template (equivalent to read 1)
trim cue
i7 (rev. comp.)
P7
4th round of sequence cycles: iterates across "i5/read1" strand, reporting the sequence of 'read 2'

**S5. Overview of PCR1 & 2 for amplicon sequencing.**
Library preparation for Illumina® sequencing often involves *i)* fragmentation of a DNA or cDNA sample to 100-1000 bp, with 5' and 3' (P5 & P7) adapter ligation, or *ii)* combined fragmentation and ligation ('tagmentation'; Nextera workflows), followed by PCR-amplification and clean-up. In amplicon sequencing as described here, sequencing libraries are prepared by two-step PCR (PCR1 + PCR2).
(a) **PCR1** produces the target amplicon (future read 1 and read2 zones) with flanking sequence that introduces Nextera adaptor sequences ('trim cues'). PCR1 requires custom-designed oligos specific to a target locus, with specific sample source as DNA template (*e.g.,* genomic DNA lysate);
(b) **PCR2** uses a small amount of PCR1 amplicon as input, and i5 and i7 barcode oligos (96x96 possibilities for dual indexing) as primers that add P5 and i5 index on one end of the product amplicon, and P7 and i7 index on the other end of the product amplicon.
PCR2 product can be pooled from multiple independent samples, and after SPRI clean-up from unincorporated primers, the pooled amplicons represent a library ready to be quantified and applied to an Illumina® flow cell for deep sequencing. Denatured library templates anneal to surface-bound oligos complementary to the P7 & P5 library adapters, and are locally amplified into physically separate clusters by bridge amplification; after sequencing, data processing includes demultiplexing of barcoded reads, trimming of adaptor sequences from reads ('trim cues') if adaptor template was encroached during sequencing, and documentation of quality scores for each sequenced nucleotide, all output to individual fastq files (Illumina® Pub. No. 770-2012-008-B (2017)).

Note regarding P5, P7, & Nextera adapter sequences (in concordance with Illumina® Document #1000000002694 v10, Feb. 2019): *Oligonucleotide sequences © 2018 Illumina®, Inc. All rights reserved. Derivative works created by Illumina® customers are authorized for use with Illumina® instruments and products only. All other uses are strictly prohibited.*

# a

#### PCR 1

**'read2 oligo'** [aka 'rev']
(sequence contiguous with 5'->3' bases that occur in this oligo is future read2/future read1 template)

5 ' – TCGGAGATGTGTATAAGAGACAG TTACTGCTGTGCACTCTCTTTCC –3 '
*Nextera read2 addition* *target-specific sequence (example is for FKBP5 GOR +86 kb)*

**'read1 oligo'** [aka 'fwd']
(sequence contiguous with 5'->3' bases that occur in this oligo is future read1/future read2 template)

5 ' – TCAGATGTGTATAAGAGACAG ACTTAAACTGGAGCTCTGACTTATTGTTC –3 '
*Nextera read1 addition* *target-specific sequence (example is for FKBP5 GOR +86 kb)*

***amplicon product:***

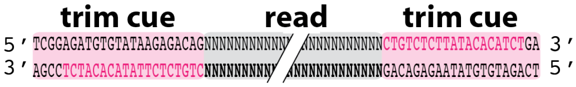


*expected size (bp):*

*+23 bp [target amplicon length] +21 bp = [target amplicon length] +44 bp*

# b

#### PCR 2

**'i7 oligo'** [aka 'read2_bc_1_TGACGTAC']
(sequence contiguous with 5'->3' bases that occur in this oligo is future read2/future read1 template)

5 ' – CAAGCAGAAGACGGCATACGAGATTGACGTACGTCTCGTGGGCTCGGAGATGTGTATAAGAGACAG –3 '
P7 i7
*P7 & i7 addition* *PCR1 amplicon product overlap*

**'i5 oligo'** [aka 'read1_bc_1_GAGGTAGT']
(sequence contiguous with 5'->3' bases that occur in this oligo is future read1/future read2 template)

5 ' – AATGATACGGCGACCACCGAGATCTACACGAGGTAGTTCGTCGGCAGCGTCAGATGTGTATAAGAGACAG –3 '
P5 i5
*P5 & i5 addition* *PCR1 amplicon product overlap*

***amplicon product:***

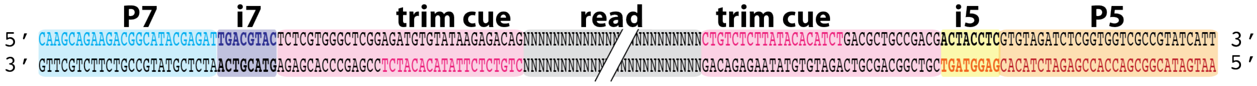


*expected size (bp):*

*+42 bp [PCR1 amplicon length] +48 bp = [PCR1 amplicon length] +90 bp*

*( [target amplicon length]+134 bp )*

**S6. Command line interface 'plateviews' of barcode sequences (i7 & i5).**

CLI Plateviews of i7 & i5 primer sets available to user of SampleSheet.py, with PrettyTable installation. The integers 1-96 above the barcode ID (*e.g.,* i7B03) designate barcode identifiers, and are the barcode labels specified by a user as input when prompted. For example, the integer '15' entered at the console specifies 'i7B03' (barcode sequence 'GTGTCCTA' in Workflow A). SampleSheet.py anticipates i7 barcodes to specify up to 96 individual wells of a plate, and i5 barcodes to specify individual plates (up to 96 separate 96-well plates). Sequences displayed (and rendered in Sample Sheet [Data] fields) correspond to 'forward' or 'reverse complement' orientations, defined by Workflow A or B.

***Command line interface:***

***<u>Workflow A: i5 'forward', i7 'reverse complement'</u>***

Please note, each 8-bp barcode sequence as displayed in this table is the sequence to be used in a Workflow A Sample Sheet barcode field.
The displayed sequence is the reverse complement of the barcode sequence as it occurs in the i7 primer.

| | 1 | 2 | 3 | 4 | 5 | 6 | 7 | 8 | 9 | 10 | 11 | 12 |
|---|---|---|---|---|---|---|---|---|---|---|---|---|
| A | 1<br>i7A01<br>GTACGTCA | 2<br>i7A02<br>TGCAGTTA | 3<br>i7A03<br>ACTGTGGA | 4<br>i7A04<br>GGTTAAGA | 5<br>i7A05<br>TCACACTA | 6<br>i7A06<br>TAGAGGTA | 7<br>i7A07<br>GCGACATA | 8<br>i7A08<br>TACATGCA | 9<br>i7A09<br>GATGATGA | 10<br>i7A10<br>TGTGTGCA | 11<br>i7A11<br>TCGCTACA | 12<br>i7A12<br>AAGCTAGA |
| B | 13<br>i7B01<br>TAGGACCA | 14<br>i7B02<br>TCGTTGGA | 15<br>i7B03<br>GTGTCCTA | 16<br>i7B04<br>TCCGTATA | 17<br>i7B05<br>TACGCTTA | 18<br>i7B06<br>TACAACGA | 19<br>i7B07<br>GAAGGAGA | 20<br>i7B08<br>AAGCATCA | 21<br>i7B09<br>TGAGATCA | 22<br>i7B10<br>TCTAGACA | 23<br>i7B11<br>TGCTCATA | 24<br>i7B12<br>TAACTCCA |
| C | 25<br>i7C01<br>GACACACA | 26<br>i7C02<br>TTGTAGCA | 27<br>i7C03<br>GTCTACGA | 28<br>i7C04<br>GCATTACA | 29<br>i7C05<br>GTTCCATA | 30<br>i7C06<br>GAATACCA | 31<br>i7C07<br>TCACTTGA | 32<br>i7C08<br>ACACAAGA | 33<br>i7C09<br>ACTACAGA | 34<br>i7C10<br>TCTCAGGA | 35<br>i7C11<br>ATCGTGCA | 36<br>i7C12<br>TCCACGTA |
| D | 37<br>i7D01<br>ATCCATGA | 38<br>i7D02<br>TTCGCAGA | 39<br>i7D03<br>TCGTCTCA | 40<br>i7D04<br>ACTTACGA | 41<br>i7D05<br>TCGGATTA | 42<br>i7D06<br>TTGGTCTA | 43<br>i7D07<br>ATAGCGGA | 44<br>i7D08<br>TGTAACCA | 45<br>i7D09<br>ACAGGCTA | 46<br>i7D10<br>AAGTCGCA | 47<br>i7D11<br>GAGTTCGA | 48<br>i7D12<br>ACTCTTCA |
| E | 49<br>i7E01<br>AAGACCTA | 50<br>i7E02<br>ACCATCCA | 51<br>i7E03<br>GAACGGTA | 52<br>i7E04<br>TAGTGAGA | 53<br>i7E05<br>GTCCAACA | 54<br>i7E06<br>TATGGCGA | 55<br>i7E07<br>GCTTATCA | 56<br>i7E08<br>GTGACTGA | 57<br>i7E09<br>ACGTGATA | 58<br>i7E10<br>GGTTCTTA | 59<br>i7E11<br>GATCAGCA | 60<br>i7E12<br>GGACAATA |
| F | 61<br>i7F01<br>AACTCTGA | 62<br>i7F02<br>TGACCTTA | 63<br>i7F03<br>GGAAGTGA | 64<br>i7F04<br>AAGAAGGA | 65<br>i7F05<br>GCAATCTA | 66<br>i7F06<br>ACGAGTCA | 67<br>i7F07<br>TTATGCCA | 68<br>i7F08<br>GCCATAGA | 69<br>i7F09<br>TAATCGGA | 70<br>i7F10<br>TCTCGTTA | 71<br>i7F11<br>GGTCTGTA | 72<br>i7F12<br>ATCCGGTA |
| G | 73<br>i7G01<br>ATGCGACA | 74<br>i7G02<br>TGAACGCA | 75<br>i7G03<br>TGGTATGA | 76<br>i7G04<br>TGGACAGA | 77<br>i7G05<br>TTGCAAGA | 78<br>i7G06<br>ACTAGGTA | 79<br>i7G07<br>TACCGATA | 80<br>i7G08<br>ATTGGTGA | 81<br>i7G09<br>TTCCTCGA | 82<br>i7G10<br>TCATGGTA | 83<br>i7G11<br>TATCCACA | 84<br>i7G12<br>GACTTGTA |
| H | 85<br>i7H01<br>ATGGAGTA | 86<br>i7H02<br>GTCAGATA | 87<br>i7H03<br>GAGCACTA | 88<br>i7H04<br>GTCATTCA | 89<br>i7H05<br>TCCTAAGA | 90<br>i7H06<br>GATGCGTA | 91<br>i7H07<br>TGCTTCCA | 92<br>i7H08<br>ATGTCAGA | 93<br>i7H09<br>GTTGCTCA | 94<br>i7H10<br>GATATCCA | 95<br>i7H11<br>GTTAGCGA | 96<br>i7H12<br>GAGGTACA |

Please note, each 8-bp barcode sequence as displayed in this table is the sequence to be used in a Workflow A Sample Sheet barcode field.
The displayed sequence is identical to the barcode sequence as it occurs in the i5 primer.

| | 1 | 2 | 3 | 4 | 5 | 6 | 7 | 8 | 9 | 10 | 11 | 12 |
|---|---|---|---|---|---|---|---|---|---|---|---|---|
| A | 1<br>i5A01<br>GAGGTAGT | 2<br>i5A02<br>GCTTAACT | 3<br>i5A03<br>GCAATTCT | 4<br>i5A04<br>TCCTCACT | 5<br>i5A05<br>AGTTAGCT | 6<br>i5A06<br>TCATGGCT | 7<br>i5A07<br>AGCCTAAT | 8<br>i5A08<br>AACACTCT | 9<br>i5A09<br>AGAGCTCT | 10<br>i5A10<br>GGCCATAT | 11<br>i5A11<br>TAGAACGT | 12<br>i5A12<br>GGTTGAAT |
| B | 13<br>i5B01<br>GATCCTAT | 14<br>i5B02<br>TGGTCTCT | 15<br>i5B03<br>TCCAGTCT | 16<br>i5B04<br>TAGCGTAT | 17<br>i5B05<br>GCGATCAT | 18<br>i5B06<br>AGCTACAT | 19<br>i5B07<br>ACTCATCT | 20<br>i5B08<br>TTGCGAGT | 21<br>i5B09<br>AAGCGACT | 22<br>i5B10<br>TGCACGAT | 23<br>i5B11<br>TATGCGGT | 24<br>i5B12<br>TCGTCGAT |
| C | 25<br>i5C01<br>ACACTTGT | 26<br>i5C02<br>TACGGCAT | 27<br>i5C03<br>TCTCGTGT | 28<br>i5C04<br>GCAGATGT | 29<br>i5C05<br>TCCGTTGT | 30<br>i5C06<br>TTCGCAGT | 31<br>i5C07<br>GGAATAGT | 32<br>i5C08<br>GCAACAAT | 33<br>i5C09<br>ACTGCGAT | 34<br>i5C10<br>TGAGTACT | 35<br>i5C11<br>ACACGAAT | 36<br>i5C12<br>GTGGCAAT |
| D | 37<br>i5D01<br>GTCATCCT | 38<br>i5D02<br>ACCGTACT | 39<br>i5D03<br>GTACTACT | 40<br>i5D04<br>TAGGTCCT | 41<br>i5D05<br>GTTACTGT | 42<br>i5D06<br>GCCTTAGT | 43<br>i5D07<br>AACCATGT | 44<br>i5D08<br>GATGACAT | 45<br>i5D09<br>AGAAGACT | 46<br>i5D10<br>TCAAGCAT | 47<br>i5D11<br>TGGATGCT | 48<br>i5D12<br>ACTTGCCT |
| E | 49<br>i5E01<br>GTTGTGAT | 50<br>i5E02<br>TATCTCGT | 51<br>i5E03<br>TGTAAGGT | 52<br>i5E04<br>ACGTTGGT | 53<br>i5E05<br>AGGAGGAT | 54<br>i5E06<br>GAGCAGAT | 55<br>i5E07<br>TCTCTACT | 56<br>i5E08<br>GGATTCAT | 57<br>i5E09<br>GACCTTCT | 58<br>i5E10<br>ACGGTTAT | 59<br>i5E11<br>AGATGCGT | 60<br>i5E12<br>TGACACCT |
| F | 61<br>i5F01<br>ACTAACGT | 62<br>i5F02<br>GAATGCCT | 63<br>i5F03<br>GTGTACGT | 64<br>i5F04<br>AAGTCAGT | 65<br>i5F05<br>TTCCTGGT | 66<br>i5F06<br>ACCACCAT | 67<br>i5F07<br>GAGTCCAT | 68<br>i5F08<br>AGCTCTGT | 69<br>i5F09<br>GGTCTTGT | 70<br>i5F10<br>AATGTGCT | 71<br>i5F11<br>GTCTGGAT | 72<br>i5F12<br>GCGTATAT |
| G | 73<br>i5G01<br>TGACTGAT | 74<br>i5G02<br>TGTCCAGT | 75<br>i5G03<br>TTGTGCCT | 76<br>i5G04<br>ACAGACCT | 77<br>i5G05<br>GATCAAGT | 78<br>i5G06<br>GATAGGAT | 79<br>i5G07<br>GTAGTCGT | 80<br>i5G08<br>TCACCTAT | 81<br>i5G09<br>AGTGGTAT | 82<br>i5G10<br>GCTAGAGT | 83<br>i5G11<br>TTAACGCT | 84<br>i5G12<br>TCAGGAGT |
| H | 85<br>i5H01<br>AACTGGCT | 86<br>i5H02<br>TACAGAGT | 87<br>i5H03<br>TTGCATCT | 88<br>i5H04<br>GAACGTGT | 89<br>i5H05<br>AAGTACCT | 90<br>i5H06<br>GGTACCAT | 91<br>i5H07<br>AATCGCAT | 92<br>i5H08<br>GGTAATCT | 93<br>i5H09<br>GATTCGCT | 94<br>i5H10<br>AGCATCGT | 95<br>i5H11<br>GTTCACCT | 96<br>i5H12<br>AGGCAAGT |

### *Workflow B: i5 'reverse complement', i7 'reverse complement'*

Please note, each 8-bp barcode sequence as displayed in this table is the sequence to be used in a Workflow B Sample Sheet barcode field. The displayed sequence is the reverse complement of the barcode sequence as it occurs in the i7 primer.

| | 1 | 2 | 3 | 4 | 5 | 6 | 7 | 8 | 9 | 10 | 11 | 12 |
|---|---|---|---|---|---|---|---|---|---|---|---|---|
| A | 1<br>i7A01<br>GTACGTCA | 2<br>i7A02<br>TGCAGTTA | 3<br>i7A03<br>ACTGTGGA | 4<br>i7A04<br>GGTTAAGA | 5<br>i7A05<br>TCACACTA | 6<br>i7A06<br>TAGAGGTA | 7<br>i7A07<br>GCGACATA | 8<br>i7A08<br>TACATGCA | 9<br>i7A09<br>GATGATGA | 10<br>i7A10<br>TGTGTGCA | 11<br>i7A11<br>TCGCTACA | 12<br>i7A12<br>AAGCTAGA |
| B | 13<br>i7B01<br>TAGGACCA | 14<br>i7B02<br>TCGTTGGA | 15<br>i7B03<br>GTGTCCTA | 16<br>i7B04<br>TCCGTATA | 17<br>i7B05<br>TACGCTTA | 18<br>i7B06<br>TACAACGA | 19<br>i7B07<br>GAAGGAGA | 20<br>i7B08<br>AAGCATCA | 21<br>i7B09<br>TGAGATCA | 22<br>i7B10<br>TCTAGACA | 23<br>i7B11<br>TGCTCATA | 24<br>i7B12<br>TAACTCCA |
| C | 25<br>i7C01<br>GACACACA | 26<br>i7C02<br>TTGTAGCA | 27<br>i7C03<br>GTCTACGA | 28<br>i7C04<br>GCATTACA | 29<br>i7C05<br>GTTCCATA | 30<br>i7C06<br>GAATACCA | 31<br>i7C07<br>TCACTTGA | 32<br>i7C08<br>ACACAAGA | 33<br>i7C09<br>ACTACAGA | 34<br>i7C10<br>TCTCAGGA | 35<br>i7C11<br>ATCGTGCA | 36<br>i7C12<br>TCCACGTA |
| D | 37<br>i7D01<br>ATCCATGA | 38<br>i7D02<br>TTCGCAGA | 39<br>i7D03<br>TCGTCTCA | 40<br>i7D04<br>ACTTACGA | 41<br>i7D05<br>TCGGATTA | 42<br>i7D06<br>TTGGTCTA | 43<br>i7D07<br>ATAGCGGA | 44<br>i7D08<br>TGTAACCA | 45<br>i7D09<br>ACAGGCTA | 46<br>i7D10<br>AAGTCGCA | 47<br>i7D11<br>GAGTTCGA | 48<br>i7D12<br>ACTCTTCA |
| E | 49<br>i7E01<br>AAGACCTA | 50<br>i7E02<br>ACCATCCA | 51<br>i7E03<br>GAACGGTA | 52<br>i7E04<br>TAGTGAGA | 53<br>i7E05<br>GTCCAACA | 54<br>i7E06<br>TATGGCGA | 55<br>i7E07<br>GCTTATCA | 56<br>i7E08<br>GTGACTGA | 57<br>i7E09<br>ACGTGATA | 58<br>i7E10<br>GGTTCTTA | 59<br>i7E11<br>GATCAGCA | 60<br>i7E12<br>GGACAATA |
| F | 61<br>i7F01<br>AACTCTGA | 62<br>i7F02<br>TGACCTTA | 63<br>i7F03<br>GGAAGTGA | 64<br>i7F04<br>AAGAAGGA | 65<br>i7F05<br>GCAATCTA | 66<br>i7F06<br>ACGAGTCA | 67<br>i7F07<br>TTATGCCA | 68<br>i7F08<br>GCCATAGA | 69<br>i7F09<br>TAATCGGA | 70<br>i7F10<br>TCTCGTTA | 71<br>i7F11<br>GGTCTGTA | 72<br>i7F12<br>ATCCGGTA |
| G | 73<br>i7G01<br>ATGCGACA | 74<br>i7G02<br>TGAACGCA | 75<br>i7G03<br>TGGTATGA | 76<br>i7G04<br>TGGACAGA | 77<br>i7G05<br>TTGCAAGA | 78<br>i7G06<br>ACTAGGTA | 79<br>i7G07<br>TACCGATA | 80<br>i7G08<br>ATTGGTGA | 81<br>i7G09<br>TTCCTCGA | 82<br>i7G10<br>TCATGGTA | 83<br>i7G11<br>TATCCACA | 84<br>i7G12<br>GACTTGTA |
| H | 85<br>i7H01<br>ATGGAGTA | 86<br>i7H02<br>GTCAGATA | 87<br>i7H03<br>GAGCACTA | 88<br>i7H04<br>GTCATTCA | 89<br>i7H05<br>TCCTAAGA | 90<br>i7H06<br>GATGCGTA | 91<br>i7H07<br>TGCTTCCA | 92<br>i7H08<br>ATGTCAGA | 93<br>i7H09<br>GTTGCTCA | 94<br>i7H10<br>GATATCCA | 95<br>i7H11<br>GTTAGCGA | 96<br>i7H12<br>GAGGTACA |

Please note, each 8-bp barcode sequence as displayed in this table is the sequence to be used in a Workflow B Sample Sheet barcode field. The displayed sequence is the reverse complement of the barcode sequence as it occurs in the i5 primer.

| | 1 | 2 | 3 | 4 | 5 | 6 | 7 | 8 | 9 | 10 | 11 | 12 |
|---|---|---|---|---|---|---|---|---|---|---|---|---|
| A | 1<br>i5A01<br>CTCCATCA | 2<br>i5A02<br>CGAATTGA | 3<br>i5A03<br>CGTTAAGA | 4<br>i5A04<br>AGGAGTGA | 5<br>i5A05<br>TCAATCGA | 6<br>i5A06<br>AGTACCGA | 7<br>i5A07<br>TCGGATTA | 8<br>i5A08<br>TTGTGAGA | 9<br>i5A09<br>TCTCGAGA | 10<br>i5A10<br>CCGGTATA | 11<br>i5A11<br>ATCTTGCA | 12<br>i5A12<br>CCAACTTA |
| B | 13<br>i5B01<br>CTAGGATA | 14<br>i5B02<br>ACCAGAGA | 15<br>i5B03<br>AGGTCAGA | 16<br>i5B04<br>ATCGCATA | 17<br>i5B05<br>CGCTAGTA | 18<br>i5B06<br>TCGATGTA | 19<br>i5B07<br>TGAGTAGA | 20<br>i5B08<br>AACGCTCA | 21<br>i5B09<br>TTCGCTGA | 22<br>i5B10<br>ACGTGCTA | 23<br>i5B11<br>ATACGCCA | 24<br>i5B12<br>AGCAGCTA |
| C | 25<br>i5C01<br>TGTGAACA | 26<br>i5C02<br>ATGCCGTA | 27<br>i5C03<br>AGAGCACA | 28<br>i5C04<br>CGTCTACA | 29<br>i5C05<br>AGGCAACA | 30<br>i5C06<br>AAGCGTCA | 31<br>i5C07<br>CCTTATCA | 32<br>i5C08<br>CGTTGTTA | 33<br>i5C09<br>TGACGCTA | 34<br>i5C10<br>ACTCATGA | 35<br>i5C11<br>TGTGCTTA | 36<br>i5C12<br>CACCGTTA |
| D | 37<br>i5D01<br>CAGTAGGA | 38<br>i5D02<br>TGGCATGA | 39<br>i5D03<br>CATGATGA | 40<br>i5D04<br>ATCCAGGA | 41<br>i5D05<br>CAATGACA | 42<br>i5D06<br>CGGAATCA | 43<br>i5D07<br>TTGGTACA | 44<br>i5D08<br>CTACTGTA | 45<br>i5D09<br>TCTTCTGA | 46<br>i5D10<br>AGTTCGTA | 47<br>i5D11<br>ACCTACGA | 48<br>i5D12<br>TGAACGGA |
| E | 49<br>i5E01<br>CAACACTA | 50<br>i5E02<br>ATAGAGCA | 51<br>i5E03<br>ACATTCCA | 52<br>i5E04<br>TGCAACCA | 53<br>i5E05<br>TCCTCCTA | 54<br>i5E06<br>CTCGTCTA | 55<br>i5E07<br>AGAGATGA | 56<br>i5E08<br>CCTAAGTA | 57<br>i5E09<br>CTGGAAGA | 58<br>i5E10<br>TGCCAATA | 59<br>i5E11<br>TCTACGCA | 60<br>i5E12<br>ACTGTGGA |
| F | 61<br>i5F01<br>TGATTGCA | 62<br>i5F02<br>CTTACGGA | 63<br>i5F03<br>CACATGCA | 64<br>i5F04<br>TTCAGTCA | 65<br>i5F05<br>AAGGACCA | 66<br>i5F06<br>TGGTGGTA | 67<br>i5F07<br>CTCAGGTA | 68<br>i5F08<br>TCGAGACA | 69<br>i5F09<br>CCAGAACA | 70<br>i5F10<br>TTACACGA | 71<br>i5F11<br>CAGACCTA | 72<br>i5F12<br>CGCATATA |
| G | 73<br>i5G01<br>ACTGACTA | 74<br>i5G02<br>ACAGGTCA | 75<br>i5G03<br>AACACGGA | 76<br>i5G04<br>TGTCTGGA | 77<br>i5G05<br>CTAGTTCA | 78<br>i5G06<br>CTATCCTA | 79<br>i5G07<br>CATCAGCA | 80<br>i5G08<br>AGTGGATA | 81<br>i5G09<br>TCACCATA | 82<br>i5G10<br>CGATCTCA | 83<br>i5G11<br>AATTGCGA | 84<br>i5G12<br>AGTCCTCA |
| H | 85<br>i5H01<br>TTGACCGA | 86<br>i5H02<br>ATGTCTCA | 87<br>i5H03<br>AACGTAGA | 88<br>i5H04<br>CTTGCACA | 89<br>i5H05<br>TTCATGGA | 90<br>i5H06<br>CCATGGTA | 91<br>i5H07<br>TTAGCGTA | 92<br>i5H08<br>CCATTAGA | 93<br>i5H09<br>CTAAGCGA | 94<br>i5H10<br>TCGTAGCA | 95<br>i5H11<br>CAAGTGGA | 96<br>i5H12<br>TCCGTTCA |

***Jupyter notebook interface:***

***Workflow A: i5 'forward', i7 'reverse complement'***

| | 1 | 2 | 3 | 4 | 5 | 6 | 7 | 8 | 9 | 10 | 11 | 12 |
|---|---|---|---|---|---|---|---|---|---|---|---|---|
| A | 1<br>i7A01<br>GTACGTCA | 2<br>i7A02<br>TGCAGTTA | 3<br>i7A03<br>ACTGTGGA | 4<br>i7A04<br>GGTTAAGA | 5<br>i7A05<br>TCACACTA | 6<br>i7A06<br>TAGAGGTA | 7<br>i7A07<br>GCGACATA | 8<br>i7A08<br>TACATGCA | 9<br>i7A09<br>GATGATGA | 10<br>i7A10<br>TGTGTGCA | 11<br>i7A11<br>TCGCTACA | 12<br>i7A12<br>AAGCTAGA |
| B | 13<br>i7B01<br>TAGGACCA | 14<br>i7B02<br>TCGTTGGA | 15<br>i7B03<br>GTGTCCTA | 16<br>i7B04<br>TCCGTATA | 17<br>i7B05<br>TACGCTTA | 18<br>i7B06<br>TACAACGA | 19<br>i7B07<br>GAAGGAGA | 20<br>i7B08<br>AAGCATCA | 21<br>i7B09<br>TGAGATCA | 22<br>i7B10<br>TCTAGACA | 23<br>i7B11<br>TGCTCATA | 24<br>i7B12<br>TAACTCCA |
| C | 25<br>i7C01<br>GACACACA | 26<br>i7C02<br>TTGTAGCA | 27<br>i7C03<br>GTCTACGA | 28<br>i7C04<br>GCATTACA | 29<br>i7C05<br>GTTCCATA | 30<br>i7C06<br>GAATACCA | 31<br>i7C07<br>TCACTTGA | 32<br>i7C08<br>ACACAAGA | 33<br>i7C09<br>ACTACAGA | 34<br>i7C10<br>TCTCAGGA | 35<br>i7C11<br>ATCGTGCA | 36<br>i7C12<br>TCCACGTA |
| D | 37<br>i7D01<br>ATCCATGA | 38<br>i7D02<br>TTCGCAGA | 39<br>i7D03<br>TCGTCTCA | 40<br>i7D04<br>ACTTACGA | 41<br>i7D05<br>TCGGATTA | 42<br>i7D06<br>TTGGTCTA | 43<br>i7D07<br>ATAGCGGA | 44<br>i7D08<br>TGTAACCA | 45<br>i7D09<br>ACAGGCTA | 46<br>i7D10<br>AAGTCGCA | 47<br>i7D11<br>GAGTTCGA | 48<br>i7D12<br>ACTCTTCA |
| E | 49<br>i7E01<br>AAGACCTA | 50<br>i7E02<br>ACCATCCA | 51<br>i7E03<br>GAACGGTA | 52<br>i7E04<br>TAGTGAGA | 53<br>i7E05<br>GTCCAACA | 54<br>i7E06<br>TATGGCGA | 55<br>i7E07<br>GCTTATCA | 56<br>i7E08<br>GTGACTGA | 57<br>i7E09<br>ACGTGATA | 58<br>i7E10<br>GGTTCTTA | 59<br>i7E11<br>GATCAGCA | 60<br>i7E12<br>GGACAATA |
| F | 61<br>i7F01<br>AACTCTGA | 62<br>i7F02<br>TGACCTTA | 63<br>i7F03<br>GGAAGTGA | 64<br>i7F04<br>AAGAAGGA | 65<br>i7F05<br>GCAATCTA | 66<br>i7F06<br>ACGAGTCA | 67<br>i7F07<br>TTATGCCA | 68<br>i7F08<br>GCCATAGA | 69<br>i7F09<br>TAATCGGA | 70<br>i7F10<br>TCTCGTTA | 71<br>i7F11<br>GGTCTGTA | 72<br>i7F12<br>ATCCGGTA |
| G | 73<br>i7G01<br>ATGCGACA | 74<br>i7G02<br>TGAACGCA | 75<br>i7G03<br>TGGTATGA | 76<br>i7G04<br>TGGACAGA | 77<br>i7G05<br>TTGCAAGA | 78<br>i7G06<br>ACTAGGTA | 79<br>i7G07<br>TACCGATA | 80<br>i7G08<br>ATTGGTGA | 81<br>i7G09<br>TTCCTCGA | 82<br>i7G10<br>TCATGGTA | 83<br>i7G11<br>TATCCACA | 84<br>i7G12<br>GACTTGTA |
| H | 85<br>i7H01<br>ATGGAGTA | 86<br>i7H02<br>GTCAGATA | 87<br>i7H03<br>GAGCACTA | 88<br>i7H04<br>GTCATTCA | 89<br>i7H05<br>TCCTAAGA | 90<br>i7H06<br>GATGCGTA | 91<br>i7H07<br>TGCTTCCA | 92<br>i7H08<br>ATGTCAGA | 93<br>i7H09<br>GTTGCTCA | 94<br>i7H10<br>GATATCCA | 95<br>i7H11<br>GTTAGCGA | 96<br>i7H12<br>GAGGTACA |

| | 1 | 2 | 3 | 4 | 5 | 6 | 7 | 8 | 9 | 10 | 11 | 12 |
|---|---|---|---|---|---|---|---|---|---|---|---|---|
| A | 1<br>i5A01<br>GAGGTAGT | 2<br>i5A02<br>GCTTAACT | 3<br>i5A03<br>GCAATTCT | 4<br>i5A04<br>TCCTCACT | 5<br>i5A05<br>AGTTAGCT | 6<br>i5A06<br>TCATGGCT | 7<br>i5A07<br>AGCCTAAT | 8<br>i5A08<br>AACACTCT | 9<br>i5A09<br>AGAGCTCT | 10<br>i5A10<br>GGCCATAT | 11<br>i5A11<br>TAGAACGT | 12<br>i5A12<br>GGTTGAAT |
| B | 13<br>i5B01<br>GATCCTAT | 14<br>i5B02<br>TGGTCTCT | 15<br>i5B03<br>TCCAGTCT | 16<br>i5B04<br>TAGCGTAT | 17<br>i5B05<br>GCGATCAT | 18<br>i5B06<br>AGCTACAT | 19<br>i5B07<br>ACTCATCT | 20<br>i5B08<br>TTGCGAGT | 21<br>i5B09<br>AAGCGACT | 22<br>i5B10<br>TGCACGAT | 23<br>i5B11<br>TATGCGGT | 24<br>i5B12<br>TCGTCGAT |
| C | 25<br>i5C01<br>ACACTTGT | 26<br>i5C02<br>TACGGCAT | 27<br>i5C03<br>TCTCGTGT | 28<br>i5C04<br>GCAGATGT | 29<br>i5C05<br>TCCGTTGT | 30<br>i5C06<br>TTCGCAGT | 31<br>i5C07<br>GGAATAGT | 32<br>i5C08<br>GCAACAAT | 33<br>i5C09<br>ACTGCGAT | 34<br>i5C10<br>TGAGTACT | 35<br>i5C11<br>ACACGAAT | 36<br>i5C12<br>GTGGCAAT |
| D | 37<br>i5D01<br>GTCATCCT | 38<br>i5D02<br>ACCGTACT | 39<br>i5D03<br>GTACTACT | 40<br>i5D04<br>TAGGTCCT | 41<br>i5D05<br>GTTACTGT | 42<br>i5D06<br>GCCTTAGT | 43<br>i5D07<br>AACCATGT | 44<br>i5D08<br>GATGACAT | 45<br>i5D09<br>AGAAGACT | 46<br>i5D10<br>TCAAGCAT | 47<br>i5D11<br>TGGATGCT | 48<br>i5D12<br>ACTTGCCT |
| E | 49<br>i5E01<br>GTTGTGAT | 50<br>i5E02<br>TATCTCGT | 51<br>i5E03<br>TGTAAGGT | 52<br>i5E04<br>ACGTTGGT | 53<br>i5E05<br>AGGAGGAT | 54<br>i5E06<br>GAGCAGAT | 55<br>i5E07<br>TCTCTACT | 56<br>i5E08<br>GGATTCAT | 57<br>i5E09<br>GACCTTCT | 58<br>i5E10<br>ACGGTTAT | 59<br>i5E11<br>AGATGCGT | 60<br>i5E12<br>TGACACCT |
| F | 61<br>i5F01<br>ACTAACGT | 62<br>i5F02<br>GAATGCCT | 63<br>i5F03<br>GTGTACGT | 64<br>i5F04<br>AAGTCAGT | 65<br>i5F05<br>TTCCTGGT | 66<br>i5F06<br>ACCACCAT | 67<br>i5F07<br>GAGTCCAT | 68<br>i5F08<br>AGCTCTGT | 69<br>i5F09<br>GGTCTTGT | 70<br>i5F10<br>AATGTGCT | 71<br>i5F11<br>GTCTGGAT | 72<br>i5F12<br>GCGTATAT |
| G | 73<br>i5G01<br>TGACTGAT | 74<br>i5G02<br>TGTCCAGT | 75<br>i5G03<br>TTGTGCCT | 76<br>i5G04<br>ACAGACCT | 77<br>i5G05<br>GATCAAGT | 78<br>i5G06<br>GATAGGAT | 79<br>i5G07<br>GTAGTCGT | 80<br>i5G08<br>TCACCTAT | 81<br>i5G09<br>AGTGGTAT | 82<br>i5G10<br>GCTAGAGT | 83<br>i5G11<br>TTAACGCT | 84<br>i5G12<br>TCAGGAGT |
| H | 85<br>i5H01<br>AACTGGCT | 86<br>i5H02<br>TACAGAGT | 87<br>i5H03<br>TTGCATCT | 88<br>i5H04<br>GAACGTGT | 89<br>i5H05<br>AAGTACCT | 90<br>i5H06<br>GGTACCAT | 91<br>i5H07<br>AATCGCAT | 92<br>i5H08<br>GGTAATCT | 93<br>i5H09<br>GATTCGCT | 94<br>i5H10<br>AGCATCGT | 95<br>i5H11<br>GTTCACCT | 96<br>i5H12<br>AGGCAAGT |

#### ***Workflow B: i5 ‘reverse complement’, i7 ‘reverse complement’***

| | 1 | 2 | 3 | 4 | 5 | 6 | 7 | 8 | 9 | 10 | 11 | 12 |
|---|---|---|---|---|---|---|---|---|---|---|---|---|
| **A** | 1<br>i7A01<br>GTACGTCA | 2<br>i7A02<br>TGCAGTTA | 3<br>i7A03<br>ACTGTGGA | 4<br>i7A04<br>GGTTAAGA | 5<br>i7A05<br>TCACACTA | 6<br>i7A06<br>TAGAGGTA | 7<br>i7A07<br>GCGACATA | 8<br>i7A08<br>TACATGCA | 9<br>i7A09<br>GATGATGA | 10<br>i7A10<br>TGTGTGCA | 11<br>i7A11<br>TCGCTACA | 12<br>i7A12<br>AAGCTAGA |
| **B** | 13<br>i7B01<br>TAGGACCA | 14<br>i7B02<br>TCGTTGGA | 15<br>i7B03<br>GTGTCCTA | 16<br>i7B04<br>TCCGTATA | 17<br>i7B05<br>TACGCTTA | 18<br>i7B06<br>TACAACGA | 19<br>i7B07<br>GAAGGAGA | 20<br>i7B08<br>AAGCATCA | 21<br>i7B09<br>TGAGATCA | 22<br>i7B10<br>TCTAGACA | 23<br>i7B11<br>TGCTCATA | 24<br>i7B12<br>TAACTCCA |
| **C** | 25<br>i7C01<br>GACACACA | 26<br>i7C02<br>TTGTAGCA | 27<br>i7C03<br>GTCTACGA | 28<br>i7C04<br>GCATTACA | 29<br>i7C05<br>GTTCCATA | 30<br>i7C06<br>GAATACCA | 31<br>i7C07<br>TCACTTGA | 32<br>i7C08<br>ACACAAGA | 33<br>i7C09<br>ACTACAGA | 34<br>i7C10<br>TCTCAGGA | 35<br>i7C11<br>ATCGTGCA | 36<br>i7C12<br>TCCACGTA |
| **D** | 37<br>i7D01<br>ATCCATGA | 38<br>i7D02<br>TTCGCAGA | 39<br>i7D03<br>TCGTCTCA | 40<br>i7D04<br>ACTTACGA | 41<br>i7D05<br>TCGGATTA | 42<br>i7D06<br>TTGGTCTA | 43<br>i7D07<br>ATAGCGGA | 44<br>i7D08<br>TGTAACCA | 45<br>i7D09<br>ACAGGCTA | 46<br>i7D10<br>AAGTCGCA | 47<br>i7D11<br>GAGTTCGA | 48<br>i7D12<br>ACTCTTCA |
| **E** | 49<br>i7E01<br>AAGACCTA | 50<br>i7E02<br>ACCATCCA | 51<br>i7E03<br>GAACGGTA | 52<br>i7E04<br>TAGTGAGA | 53<br>i7E05<br>GTCCAACA | 54<br>i7E06<br>TATGGCGA | 55<br>i7E07<br>GCTTATCA | 56<br>i7E08<br>GTGACTGA | 57<br>i7E09<br>ACGTGATA | 58<br>i7E10<br>GGTTCTTA | 59<br>i7E11<br>GATCAGCA | 60<br>i7E12<br>GGACAATA |
| **F** | 61<br>i7F01<br>AACTCTGA | 62<br>i7F02<br>TGACCTTA | 63<br>i7F03<br>GGAAGTGA | 64<br>i7F04<br>AAGAAGGA | 65<br>i7F05<br>GCAATCTA | 66<br>i7F06<br>ACGAGTCA | 67<br>i7F07<br>TTATGCCA | 68<br>i7F08<br>GCCATAGA | 69<br>i7F09<br>TAATCGGA | 70<br>i7F10<br>TCTCGTTA | 71<br>i7F11<br>GGTCTGTA | 72<br>i7F12<br>ATCCGGTA |
| **G** | 73<br>i7G01<br>ATGCGACA | 74<br>i7G02<br>TGAACGCA | 75<br>i7G03<br>TGGTATGA | 76<br>i7G04<br>TGGACAGA | 77<br>i7G05<br>TTGCAAGA | 78<br>i7G06<br>ACTAGGTA | 79<br>i7G07<br>TACCGATA | 80<br>i7G08<br>ATTGGTGA | 81<br>i7G09<br>TTCCTCGA | 82<br>i7G10<br>TCATGGTA | 83<br>i7G11<br>TATCCACA | 84<br>i7G12<br>GACTTGTA |
| **H** | 85<br>i7H01<br>ATGGAGTA | 86<br>i7H02<br>GTCAGATA | 87<br>i7H03<br>GAGCACTA | 88<br>i7H04<br>GTCATTCA | 89<br>i7H05<br>TCCTAAGA | 90<br>i7H06<br>GATGCGTA | 91<br>i7H07<br>TGCTTCCA | 92<br>i7H08<br>ATGTCAGA | 93<br>i7H09<br>GTTGCTCA | 94<br>i7H10<br>GATATCCA | 95<br>i7H11<br>GTTAGCGA | 96<br>i7H12<br>GAGGTACA |

| | 1 | 2 | 3 | 4 | 5 | 6 | 7 | 8 | 9 | 10 | 11 | 12 |
|---|---|---|---|---|---|---|---|---|---|---|---|---|
| **A** | 1<br>i5A01<br>CTCCATCA | 2<br>i5A02<br>CGAATTGA | 3<br>i5A03<br>CGTTAAGA | 4<br>i5A04<br>AGGAGTGA | 5<br>i5A05<br>TCAATCGA | 6<br>i5A06<br>AGTACCGA | 7<br>i5A07<br>TCGGATTA | 8<br>i5A08<br>TTGTGAGA | 9<br>i5A09<br>TCTCGAGA | 10<br>i5A10<br>CCGGTATA | 11<br>i5A11<br>ATCTTGCA | 12<br>i5A12<br>CCAACTTA |
| **B** | 13<br>i5B01<br>CTAGGATA | 14<br>i5B02<br>ACCAGAGA | 15<br>i5B03<br>AGGTCAGA | 16<br>i5B04<br>ATCGCATA | 17<br>i5B05<br>CGCTAGTA | 18<br>i5B06<br>TCGATGTA | 19<br>i5B07<br>TGAGTAGA | 20<br>i5B08<br>AACGCTCA | 21<br>i5B09<br>TTCGCTGA | 22<br>i5B10<br>ACGTGCTA | 23<br>i5B11<br>ATACGCCA | 24<br>i5B12<br>AGCAGCTA |
| **C** | 25<br>i5C01<br>TGTGAACA | 26<br>i5C02<br>ATGCCGTA | 27<br>i5C03<br>AGAGCACA | 28<br>i5C04<br>CGTCTACA | 29<br>i5C05<br>AGGCAACA | 30<br>i5C06<br>AAGCGTCA | 31<br>i5C07<br>CCTTATCA | 32<br>i5C08<br>CGTTGTTA | 33<br>i5C09<br>TGACGCTA | 34<br>i5C10<br>ACTCATGA | 35<br>i5C11<br>TGTGCTTA | 36<br>i5C12<br>CACCGTTA |
| **D** | 37<br>i5D01<br>CAGTAGGA | 38<br>i5D02<br>TGGCATGA | 39<br>i5D03<br>CATGATGA | 40<br>i5D04<br>ATCCAGGA | 41<br>i5D05<br>CAATGACA | 42 Tex<br>i5D06<br>CGGAATCA | 43<br>i5D07<br>TTGGTACA | 44<br>i5D08<br>CTACTGTA | 45<br>i5D09<br>TCTTCTGA | 46<br>i5D10<br>AGTTCGTA | 47<br>i5D11<br>ACCTACGA | 48<br>i5D12<br>TGAACGGA |
| **E** | 49<br>i5E01<br>CAACACTA | 50<br>i5E02<br>ATAGAGCA | 51<br>i5E03<br>ACATTCCA | 52<br>i5E04<br>TGCAACCA | 53<br>i5E05<br>TCCTCCTA | 54<br>i5E06<br>CTCGTCTA | 55<br>i5E07<br>AGAGATGA | 56<br>i5E08<br>CCTAAGTA | 57<br>i5E09<br>CTGGAAGA | 58<br>i5E10<br>TGCCAATA | 59<br>i5E11<br>TCTACGCA | 60<br>i5E12<br>ACTGTGGA |
| **F** | 61<br>i5F01<br>TGATTGCA | 62<br>i5F02<br>CTTACGGA | 63<br>i5F03<br>CACATGCA | 64<br>i5F04<br>TTCAGTCA | 65<br>i5F05<br>AAGGACCA | 66<br>i5F06<br>TGGTGGTA | 67<br>i5F07<br>CTCAGGTA | 68<br>i5F08<br>TCGAGACA | 69<br>i5F09<br>CCAGAACA | 70<br>i5F10<br>TTACACGA | 71<br>i5F11<br>CAGACCTA | 72<br>i5F12<br>CGCATATA |
| **G** | 73<br>i5G01<br>ACTGACTA | 74<br>i5G02<br>ACAGGTCA | 75<br>i5G03<br>AACACGGA | 76<br>i5G04<br>TGTCTGGA | 77<br>i5G05<br>CTAGTTCA | 78<br>i5G06<br>CTATCCTA | 79<br>i5G07<br>CATCAGCA | 80<br>i5G08<br>AGTGGATA | 81<br>i5G09<br>TCACCATA | 82<br>i5G10<br>CGATCTCA | 83<br>i5G11<br>AATTGCGA | 84<br>i5G12<br>AGTCCTCA |
| **H** | 85<br>i5H01<br>TTGACCGA | 86<br>i5H02<br>ATGTCTCA | 87<br>i5H03<br>AACGTAGA | 88<br>i5H04<br>CTTGCACA | 89<br>i5H05<br>TTCATGGA | 90<br>i5H06<br>CCATGGTA | 91<br>i5H07<br>TTAGCGTA | 92<br>i5H08<br>CCATTAGA | 93<br>i5H09<br>CTAAGCGA | 94<br>i5H10<br>TCGTAGCA | 95<br>i5H11<br>CAAGTGGA | 96<br>i5H12<br>TCCGTTCA |

**S7. File outputs of ImputedGenotypes.py.**

(a) Example of custom-named directory (ImputedGenotypes.py user *input* #1) populated with 8 output files; filenames are automatically prefixed by script operations with system start date. ***fasta.fa*** and ***blastn_alignments.txt*** are prepared during script operations, but key script output content is contained in the other six files.

(b) ***allele_definitions.csv*** provides comma-separated panel data compatible with spreadsheet programs such as Excel, and/or import as panel data in Python pandas for user access and manipulation. This file contains raw content of *imputedgenotypes_dict,* the primary Python data object of ImputedGenotypes.py.

(c) ***allele_definitions.txt*** reports the top 10 ranked alleles for R1 (and R2 for PE data), with allele metadata (frequency metrics, allele specifications, reference sequence database alignment location and alignment), along with (optional) mapped DNA sub-sequences (*e.g.*, Cas9 guide RNA sequence(s), test sequence(s). ***imputed_genotypes.txt*** reports the same content, but ordered based on imputed genotype cohorts.

(d) ***allele_definitions.pdf*** presents a visualization of ranked allele frequencies (raw frequency relative to all other reads, and adjusted frequencies relative to reads at various abundance thresholds),

(e) ***population_summary.txt*** reports aggregate genotype and allele representations across all samples.

(f) ***script_metrics.txt*** logs script operation metadata.

**a**

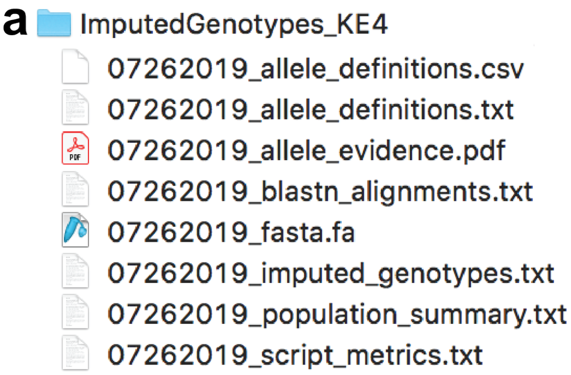


**b** *allele_definitions.csv*

| | allele | read | sample | reads | totalreads | %totalreads | %top10read | %readsfilter | %readsfilter | chr | locusID | coordinates | alignment_q | alignment_m | alignment_h | allele_type | allele_specs | imputed_genotype |
|---|---|---|---|---|---|---|---|---|---|---|---|---|---|---|---|---|---|---|
| 0 | KE4-1-A01_ | R1 | KE4-1-A01 | 163 | 389 | 41.9 | 49.85 | 51.75 | 62.93 | Homo sapier | ref\|NC_000( | 35602016-3 | query AC | \|\|\|\|\|\|\|\|\| | reference A | mutant | likely substit | \|heterozygous\| substitution + wild-type (sub/wt) |
| 1 | [illegible] | [illegible] | [illegible] | 96 | 389 | 24.68 | 29.36 | 30.48 | 37.07 | [illegible] | [illegible] | [illegible] | [illegible] | [illegible] | [illegible] | [illegible] | [illegible] | [illegible] |
| 6558 | KE4-4-G10_ | R1 | KE4-4-G10 | 530 | 1362 | 38.91 | 48.67 | 50.19 | 50.91 | Homo sapier | ref\|NC_000( | 35602016-3 | query AC | \|\|\|\|\|\|\|\|\| | reference A | mutant | likely deletic | \|heterozygous\| deletion1 + deletion2 (del1/del2) |
| 6559 | KE4-4-G10_ | R1 | KE4-4-G10 | 511 | 1362 | 37.52 | 46.92 | 48.39 | 49.09 | Homo sapier | ref\|NC_000( | 35602016-3 | query AC | \|\|\|\|\|\|\|\|\| | reference A | mutant | likely deletic | \|heterozygous\| deletion1 + deletion2 (del1/del2) |
| 6560 | KE4-4-G10_ | R1 | KE4-4-G10 | 15 | 1362 | 1.1 | 1.38 | 1.42 | None | Homo sapier | ref\|NC_000( | 35602016-3 | query AC | \|\|\|\|\|\|\|\|\| | reference A | mutant | likely deletic | \|heterozygous\| deletion1 + deletion2 (del1/del2) |
| 6561 | KE4-4-G10_ | R1 | KE4-4-G10 | 8 | 1362 | 0.59 | 0.73 | None | None | Homo sapier | ref\|NC_000( | 35602016-3 | query AC | \|\|\|\|\|\|\|\|\| | reference A | mutant | likely deletic | \|heterozygous\| deletion1 + deletion2 (del1/del2) |
| 6562 | KE4-4-G10_ | R1 | KE4-4-G10 | 5 | 1362 | 0.37 | 0.46 | None | None | Homo sapier | ref\|NC_000( | 35602016-3 | query AC | \|\|\|\|\|\|\|\|\| | reference A | mutant | likely deletic | \|heterozygous\| deletion1 + deletion2 (del1/del2) |
| 6563 | KE4-4-G10_ | R1 | KE4-4-G10 | 5 | 1362 | 0.37 | 0.46 | None | None | Homo sapier | ref\|NC_000( | 35602016-3 | query AC | \|\|\|\|\|\|\|\|\| | reference A | mutant | likely deletic | \|heterozygous\| deletion1 + deletion2 (del1/del2) |
| 6564 | KE4-4-G10_ | R1 | KE4-4-G10 | 4 | 1362 | 0.29 | 0.37 | None | None | Homo sapier | ref\|NC_000( | 35602016-3 | query AC | \|\|\|\|\|\|\|\|\| | reference A | wild-type | | \|heterozygous\| deletion1 + deletion2 (del1/del2) |
| 6565 | KE4-4-G10_ | R1 | KE4-4-G10 | 4 | 1362 | 0.29 | 0.37 | None | None | Homo sapier | ref\|NC_000( | 35602190-3 | query TT | \|\|\|\|\|\|\|\|\| | reference T | mutant | likely deletic | \|heterozygous\| deletion1 + deletion2 (del1/del2) |
| 6566 | KE4-4-G10_ | R1 | KE4-4-G10 | 4 | 1362 | 0.29 | 0.37 | None | None | Homo sapier | ref\|NC_000( | 35602019-3 | query TA | \|\|\|\|\|\|\|\|\| | reference T | mutant | likely deletic | \|heterozygous\| deletion1 + deletion2 (del1/del2) |
| 6567 | KE4-4-G10_ | R1 | KE4-4-G10 | 3 | 1362 | 0.22 | 0.28 | None | None | Homo sapier | ref\|NC_000( | 35602016-3 | query AC | \|\|\|\|\|\|\|\|\| | reference A | mutant | likely deletic | \|heterozygous\| deletion1 + deletion2 (del1/del2) |
| 6568 | KE4-4-G10_ | R2 | KE4-4-G10 | 373 | 1362 | 27.39 | 45.1 | 46.92 | 52.17 | Homo sapier | ref\|NC_000( | 35602190-3 | query TT | \|\|\|\|\|\|\|\|\| | reference T | mutant | likely deletic | \|heterozygous\| deletion1 + deletion2 (del1/del2) |
| 6569 | KE4-4-G10_ | R2 | KE4-4-G10 | 342 | 1362 | 25.11 | 41.35 | 43.02 | 47.83 | Homo sapier | ref\|NC_000( | 35602190-3 | query TT | \|\|\|\|\|\|\|\|\| | reference T | mutant | likely deletic | \|heterozygous\| deletion1 + deletion2 (del1/del2) |
| 6570 | KE4-4-G10_ | R2 | KE4-4-G10 | 49 | 1362 | 3.6 | 5.93 | 6.16 | None | Homo sapier | ref\|NC_000( | 35602190-3 | query TT | \|\|\|\|\|\|\|\|\| | reference T | mutant | likely deletic | \|heterozygous\| deletion1 + deletion2 (del1/del2) |
| 6571 | KE4-4-G10_ | R2 | KE4-4-G10 | 16 | 1362 | 1.17 | 1.93 | 2.01 | None | Homo sapier | ref\|NC_000( | 35602190-3 | query TT | \|\|\|\|\|\|\|\|\| | reference T | mutant | likely deletic | \|heterozygous\| deletion1 + deletion2 (del1/del2) |
| 6572 | KE4-4-G10_ | R2 | KE4-4-G10 | 15 | 1362 | 1.1 | 1.81 | 1.89 | None | Homo sapier | ref\|NC_000( | 35602190-3 | query TT | \|\|\|\|\|\|\|\|\| | reference T | mutant | likely deletic | \|heterozygous\| deletion1 + deletion2 (del1/del2) |
| 6573 | KE4-4-G10_ | R2 | KE4-4-G10 | 11 | 1362 | 0.81 | 1.33 | None | None | Homo sapier | ref\|NC_000( | 35602190-3 | query TT | \|\|\|\|\|\|\|\|\| | reference T | mutant | likely deletic | \|heterozygous\| deletion1 + deletion2 (del1/del2) |
| 6574 | KE4-4-G10_ | R2 | KE4-4-G10 | 7 | 1362 | 0.51 | 0.85 | None | None | Homo sapier | ref\|NC_000( | 35602190-3 | query TT | \|\|\|\|\|\|\|\|\| | reference T | mutant | likely deletic | \|heterozygous\| deletion1 + deletion2 (del1/del2) |
| 6575 | KE4-4-G10_ | R2 | KE4-4-G10 | 6 | 1362 | 0.44 | 0.73 | None | None | Homo sapier | ref\|NC_000( | 35602190-3 | query TT | \|\|\|\|\|\|\|\|\| | reference T | mutant | likely deletic | \|heterozygous\| deletion1 + deletion2 (del1/del2) |
| 6576 | KE4-4-G10_ | R2 | KE4-4-G10 | 4 | 1362 | 0.29 | 0.48 | None | None | Homo sapier | ref\|NC_000( | 35602190-3 | query TT | \|\|\|\|\|\|\|\|\| | reference T | mutant | likely deletic | \|heterozygous\| deletion1 + deletion2 (del1/del2) |
| 6577 | KE4-4-G10_ | R2 | KE4-4-G10 | 4 | 1362 | 0.29 | 0.48 | None | None | Homo sapier | ref\|NC_000( | 35602190-3 | query TT | \|\|\|\|\|\|\|\|\| | reference T | mutant | likely deletic | \|heterozygous\| deletion1 + deletion2 (del1/del2) |
| 6578 | KE4-4-G11_ | R1 | KE4-4-G11 | 1102 | 1372 | 80.32 | 94.19 | 97.52 | 100 | Homo sapier | ref\|NC_000( | 35602016-3 | query AC | \|\|\|\|\|\|\|\|\| | reference A | wild-type | | \|homozygous\| wild-type (wt/wt) |
| 6579 | KE4-4-G11_ | R1 | KE4-4-G11 | 28 | 1372 | 2.04 | 2.39 | 2.48 | None | Homo sapier | ref\|NC_000( | 35602016-3 | query AC | \|\|\|\|\|\|\|\|\| | reference A | mutant | likely deletic | \|homozygous\| wild-type (wt/wt) |
| 6580 | KE4-4-G11_ | R1 | KE4-4-G11 | 8 | 1372 | 0.58 | 0.68 | None | None | Homo sapier | ref\|NC_000( | 35602190-3 | query TT | \|\|\|\|\|\|\|\|\| | reference T | wild-type | | \|homozygous\| wild-type (wt/wt) |
| 6581 | KE4-4-G11_ | R1 | KE4-4-G11 | 8 | 1372 | 0.58 | 0.68 | None | None | Homo sapier | ref\|NC_000( | 35602019-3 | query TA | \|\|\|\|\|\|\|\|\| | reference T | wild-type | | \|homozygous\| wild-type (wt/wt) |
| 6582 | KE4-4-G11_ | R1 | KE4-4-G11 | 6 | 1372 | 0.44 | 0.51 | None | None | Homo sapier | ref\|NC_000( | 35602016-3 | query AC | \|\|\|\|\|\|\|\|\| | reference A | mutant | likely substit | \|homozygous\| wild-type (wt/wt) |
| 6583 | KE4-4-G11_ | R1 | KE4-4-G11 | 5 | 1372 | 0.36 | 0.43 | None | None | Homo sapier | ref\|NC_000( | 35602016-3 | query AC | \|\|\|\|\|\|\|\|\| | reference A | mutant | likely deletic | \|homozygous\| wild-type (wt/wt) |
| 6584 | KE4-4-G11_ | R1 | KE4-4-G11 | 4 | 1372 | 0.29 | 0.34 | None | None | Homo sapier | ref\|NC_000( | 35602016-3 | query AC | \|\|\|\|\|\|\|\|\| | reference A | mutant | likely substit | \|homozygous\| wild-type (wt/wt) |
| 6585 | KE4-4-G11_ | R1 | KE4-4-G11 | 3 | 1372 | 0.22 | 0.26 | None | None | Homo sapier | ref\|NC_000( | 35602017-3 | query CT | \|\|\|\|\|\|\|\|\| | reference C | wild-type | | \|homozygous\| wild-type (wt/wt) |
| 6586 | KE4-4-G11_ | R1 | KE4-4-G11 | 3 | 1372 | 0.22 | 0.26 | None | None | Homo sapier | ref\|NC_000( | 35602016-3 | query AC | \|\|\|\| \|\|\|\|\| | reference A | mutant | likely deletic | \|homozygous\| wild-type (wt/wt) |
| 6587 | KE4-4-G11_ | R1 | KE4-4-G11 | 3 | 1372 | 0.22 | 0.26 | None | None | Homo sapier | ref\|NC_000( | 35602016-3 | query AC | \|\|\|\|\|\|\|\|\| | reference A | mutant | likely substit | \|homozygous\| wild-type (wt/wt) |
| 6588 | KE4-4-G11_ | R2 | KE4-4-G11 | 947 | 1372 | 69.02 | 92.93 | 97.93 | 100 | Homo sapier | ref\|NC_000( | 35602190-3 | query TT | \|\|\|\|\|\|\|\|\| | reference T | wild-type | | \|homozygous\| wild-type (wt/wt) |
| 6589 | KE4-4-G11_ | R2 | KE4-4-G11 | 20 | 1372 | 1.46 | 1.96 | 2.07 | None | Homo sapier | ref\|NC_000( | 35602190-3 | query TT | \|\|\|\|\|\|\|\|\| | reference T | wild-type | | \|homozygous\| wild-type (wt/wt) |
| 6590 | KE4-4-G11_ | R2 | KE4-4-G11 | 8 | 1372 | 0.58 | 0.79 | None | None | Homo sapier | ref\|NC_000( | 35602190-3 | query TT | \|\|\|\|\|\|\|\|\| | reference T | wild-type | | \|homozygous\| wild-type (wt/wt) |

**c** *allele_definitions.txt*

```
=========
KE4-4-G10
=========
Imputed Genotype: |heterozygous| deletion1 + deletion2 (del1/del2)

   *~~~~~~~~~*
   | READ 1 |
   *~~~~~~~~~*

   Allele: KE4-4-G10 R1 [530/1362] %totalreads:38.91 percentile:100 %top10reads:48.67 %readsfilteredfor1%:50.19 %readsfilteredfor10%:50.91 | mutant, likely deletion, 4 bp
    Locus: Homo sapiens chromosome 6, GRCh38.p12 Primary Assembly, ref|NC_000006.12| 35602016-35602170

                                                                                      3'-CAAGACTTACACCGACCGTG-5' (guide sequence)
                                                                                               v
    query  ACTTAAACTGGAGCTCTGACTTATTGTTCTCTTACTGCCCTAGAGCAATTTTGTTTTGAAGAGCACAGAACACCCTGTT----ATGTGGCTGGCACATGAACTCGATGTGCTGACAGCAATTTGTACTCCGATTAAAATAGGGGGGAAAAAAGGA
           |||||||||||||||||||||||||||||||||||||||||||||||||||||||||||||||||||||||||||||||    ||||||||||||||||||||||||||||||||||||||||||||||||||||||||||||||||||||||||
reference  ACTTAAACTGGAGCTCTGACTTATTGTTCTCTTACTGCCCTAGAGCAATTTTGTTTTGAAGAGCACAGAACACCCTGTTCTGAATGTGGCTGGCACATGAACTCGATGTGCTGACAGCAATTTGTACTCCGATTAAAATAGGGGGGAAAAAAGGA
                                                                    ^^^^^^^^^^^^^^^^^
                                                                 3'-GTCTTGTGGGACAAGA-5' (test sequence)

   Allele: KE4-4-G10 R1 [511/1362] %totalreads:37.52 percentile:99 %top10reads:46.92 %readsfilteredfor1%:48.39 %readsfilteredfor10%:49.09 | mutant, likely deletion, 2 bp
    Locus: Homo sapiens chromosome 6, GRCh38.p12 Primary Assembly, ref|NC_000006.12| 35602016-35602168

                                                                                      3'-CAAGACTTACACCGACCGTG-5' (guide sequence)
                                                                                               v
    query  ACTTAAACTGGAGCTCTGACTTATTGTTCTCTTACTGCCCTAGAGCAATTTTGTTTTGAAGAGCACAGAACACCCTG--CTGAATGTGGCTGGCACATGAACTCGATGTGCTGACAGCAATTTGTACTCCGATTAAAATAGGGGGGAAAAAAG
           |||||||||||||||||||||||||||||||||||||||||||||||||||||||||||||||||||||||||||||  |||||||||||||||||||||||||||||||||||||||||||||||||||||||||||||||||||||||||||
reference  ACTTAAACTGGAGCTCTGACTTATTGTTCTCTTACTGCCCTAGAGCAATTTTGTTTTGAAGAGCACAGAACACCCTGTTCTGAATGTGGCTGGCACATGAACTCGATGTGCTGACAGCAATTTGTACTCCGATTAAAATAGGGGGGAAAAAAG
                                                                    ^^^^^^^^^^^^^^^^^
                                                                 3'-GTCTTGTGGGACAAGA-5' (test sequence)

   *~~~~~~~~~~~~~~~~~~~~~~~~~~~~~~~~~~~~~~~~~~~~~~~~~~~~~~~~~~~~*
   |  >>>>> remaining alleles occur at frequency <10% <<<<< |
   *~~~~~~~~~~~~~~~~~~~~~~~~~~~~~~~~~~~~~~~~~~~~~~~~~~~~~~~~~~~~*

   Allele: KE4-4-G10 R1 [15/1362] %totalreads:1.1 percentile:99 %top10reads:1.38 %readsfilteredfor1%:1.42 %readsfilteredfor10%:None | mutant, likely deletion, 5 bp
    Locus: Homo sapiens chromosome 6, GRCh38.p12 Primary Assembly, ref|NC_000006.12| 35602016-35602171

                                                                                      3'-CAAGACTTACACCGACCGTG-5' (guide sequence)
                                                                                               v
    query  ACTTAAACTGGAGCTCTGAC-TATTGTTCTCTTACTGCCCTAGAGCAATTTTGTTTTGAAGAGCACAGAACACCCTGTT----ATGTGGCTGGCACATGAACTCGATGTGCTGACAGCAATTTGTACTCCGATTAAAATAGGGGGGAAAAAAGGAA
           |||||||||||||||||||| ||||||||||||||||||||||||||||||||||||||||||||||||||||||||||    |||||||||||||||||||||||||||||||||||||||||||||||||||||||||||||||||||||||||
reference  ACTTAAACTGGAGCTCTGACTTATTGTTCTCTTACTGCCCTAGAGCAATTTTGTTTTGAAGAGCACAGAACACCCTGTTCTGAATGTGGCTGGCACATGAACTCGATGTGCTGACAGCAATTTGTACTCCGATTAAAATAGGGGGGAAAAAAGGAA
                                                                    ^^^^^^^^^^^^^^^^^
                                                                 3'-GTCTTGTGGGACAAGA-5' (test sequence)
```

**d** *allele_evidence.pdf*

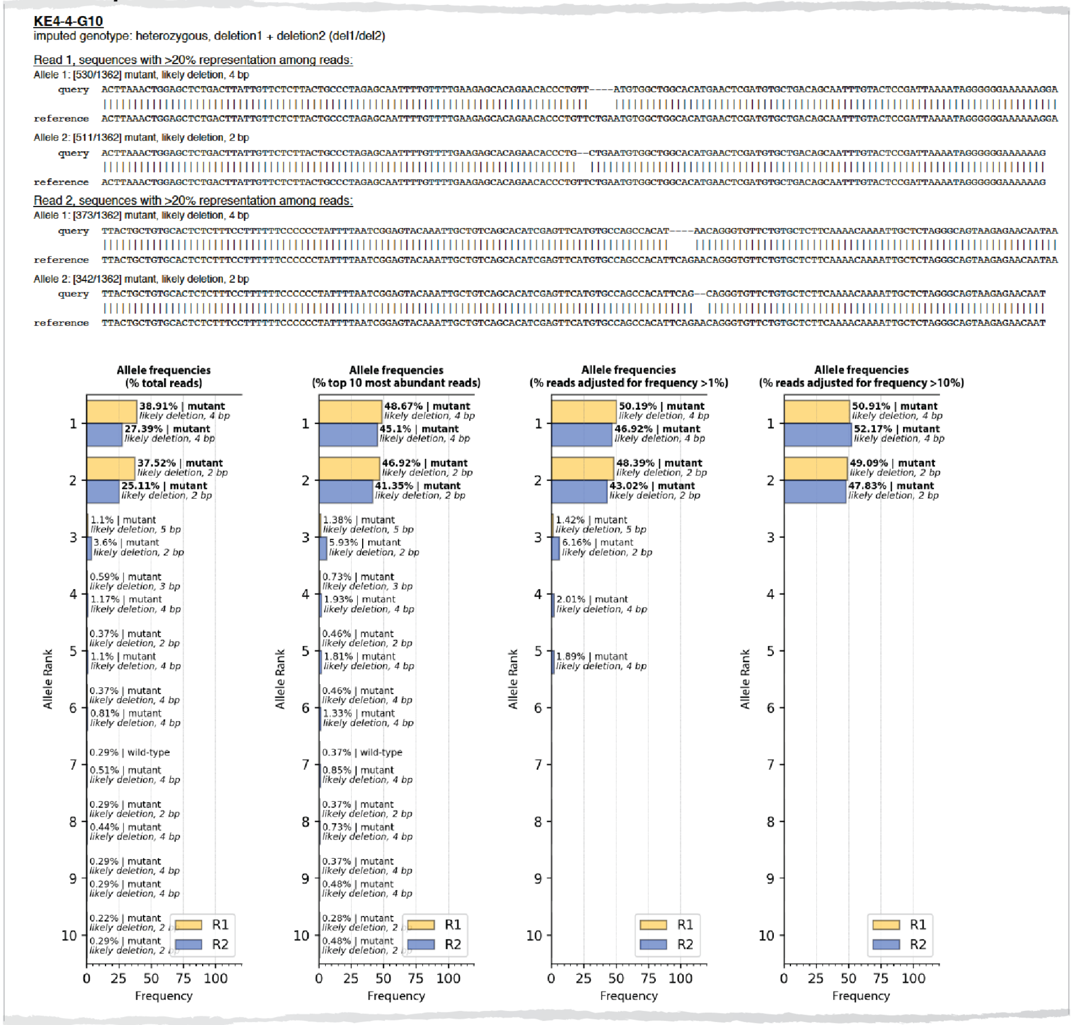

#### e *population_summary.txt*

```
ImputedGenotypes.py: Population Summary
Date: 07/26/2019

I. Synopsis of Interpretations: Allele Definitions & Genotype Imputations

    (A) Sample summary
        (i) Number of samples processed: 384
        (ii) % samples called (genotype imputed): 383 (99.74%)

    (B) Genotypes summary
        (i) % samples diploid (1-2 prominent alleles inferred): 362 (94.27%)
            (1) % homozygous wild-type (wt): 316 (82.29%)
            (2) % homozygous mutant: 12 (3.12%)
                -> % homozygous deletion: 9 (2.34%)
                -> % homozygous insertion: 1 (0.26%)
                -> % homozygous substitution: 2 (0.52%)
                -> % homozygous complex indel: 0 (0.0%)
            (3) % heterozygous (wt + mutant): 29 (7.55%)
                -> % heterozygous deletion: 12 (3.12%)
                -> % heterozygous insertion: 1 (0.26%)
                -> % heterozygous substitution: 16 (4.17%)
                -> % heterozygous complex indel: 0 (0.0%)
            (4) % heterozygous (mutant + mutant): 5 (1.3%)
                -> % heterozygous deletion + insertion: 0 (0.0%)
                -> % heterozygous deletion + substitution: 0 (0.0%)
                -> % heterozygous insertion + substitution: 0 (0.0%)
                -> % heterozygous deletion + complex indel: 0 (0.0%)
                -> % heterozygous insertion + complex indel: 0 (0.0%)
                -> % heterozygous substitution + complex indel: 0 (0.0%)
        (ii) % samples multiploid (>2 prominent alleles inferred): 5 (1.3%)

    (B) Alleles summary
        (i) % wild-type alleles: 655 (91.61% of total alleles)
        (ii) % mutant alleles: 60 (8.39% of total alleles)
            (1) % deletion alleles: 38 (5.31% of total alleles)
            (2) % insertion alleles: 1 (0.14% of total alleles)
            (3) % substitution alleles: 21 (2.94% of total alleles)
            (4) % complex indel alleles: 1 (0.14% of total alleles)
        (iii) % mutant alleles with ablated test sequence(s):
            (1) % deletion alleles:
            (2) % insertion alleles:
            (3) % substitution alleles:
            (4) % complex indel alleles:

II. Synopsis of Reads Lost to Analysis
    'Top 10' reads among samples with (A) no hits, or (B) multiple hits, in reference database

    (A) Samples with reads among the 'top 10 most abundant reads', that did not map to the reference genome
        (i) For the following sample IDs (1), NO reads among the "top 10 most abundant reads" could be mapped to the reference genome:
             KE4-1-C06
        (ii) For the following sample IDs (47), the indicated reads among the "top 10 most abundant reads" did not map to the reference genome:
             KE4-1-A08:
               R2
               KE4-1-A08 [1/6] 16.67%
               KE4-1-A08 [1/6] 16.67%
             KE4-1-B01:
               R2
               KE4-1-B01 [1/10] 10.0%
             KE4-1-B05:
               R1
               KE4-1-B05 [1/105] 0.95%
               R2
               KE4-1-B05 [1/105] 0.95%
               KE4-1-B05 [1/105] 0.95%
               KE4-1-B05 [1/105] 0.95%
             KE4-1-C02:
               R2
               KE4-1-C02 [1/171] 0.58%
               KE4-1-C02 [1/171] 0.58%
             KE4-1-C06:
               R1
               KE4-1-C06 [1/4] 25.0%
               R2
               KE4-1-C06 [1/4] 25.0%
               KE4-1-C06 [1/4] 25.0%
               KE4-1-C06 [1/4] 25.0%
```

#### f *script_metrics.txt*

```
ImputedGenotypes.py: Script Metrics
Date: 07/26/2019

Operating system information:
    name: Kirks-MBP.lan
    platform: Darwin-17.7.0-x86_64-i386-64bit
    RAM (GB): 16.0
    physical CPU/effective CPU: 4/8
    executable: /Library/Frameworks/Python.framework/Versions/3.7/Resources/Python.app/Contents/MacOS/Pythor

User-entered variables:
    output_directory: /Users/kirkehmsen/ImputedGenotypes_KE4
    fastq_directory: /Users/kirkehmsen/data/KE4
    blastn_path: /usr/local/bin/blast/bin/blastn
    db_path: /Users/kirkehmsen/blastn_database
    db_prefix: GRCh38
    guideRNA_seq: GTGCCAGCCACATTCAGAAC
    extant_seq: AGAACAGGGTGTTCTG

fastq file information:
    Illumina sequencing run ID(s): @M00582:216
    Number of fastq files processed: 768
    Size distribution of fastq files processed:
        total... 197 MB
        range... max: 0.61 MB; min: 0.00069 MB; median: 0.261 MB; mean +/- stdev: 0.257 +/- 0.132 MB
    Read distribution within fastq files to process:
        total... 569,374 reads
        range... max: 1745 reads; min: 2 reads; median: 754.0 reads; mean +/- stdev: 741.0 +/- 382.0 reads

fastq files processed (name, size (MB), reads):
    /Users/kirkehmsen/data/KE4/KE4-1-A01_S1153_L001_R1_001.fastq, 0.13484, 389
    /Users/kirkehmsen/data/KE4/KE4-1-A01_S1153_L001_R2_001.fastq, 0.13484, 389
    /Users/kirkehmsen/data/KE4/KE4-1-A02_S1154_L001_R1_001.fastq, 0.26033, 751
    /Users/kirkehmsen/data/KE4/KE4-1-A02_S1154_L001_R2_001.fastq, 0.26033, 751
    /Users/kirkehmsen/data/KE4/KE4-1-A03_S1155_L001_R1_001.fastq, 0.2347, 677
    /Users/kirkehmsen/data/KE4/KE4-1-A03_S1155_L001_R2_001.fastq, 0.2347, 677
    /Users/kirkehmsen/data/KE4/KE4-1-A04_S1156_L001_R1_001.fastq, 0.20214, 583
    /Users/kirkehmsen/data/KE4/KE4-1-A04_S1156_L001_R2_001.fastq, 0.20214, 583
    /Users/kirkehmsen/data/KE4/KE4-1-A05_S1157_L001_R1_001.fastq, 0.20627, 595
    /Users/kirkehmsen/data/KE4/KE4-1-A05_S1157_L001_R2_001.fastq, 0.20627, 595
    /Users/kirkehmsen/data/KE4/KE4-1-A06_S1158_L001_R1_001.fastq, 0.25342, 731
    /Users/kirkehmsen/data/KE4/KE4-1-A06_S1158_L001_R2_001.fastq, 0.25342, 731
    /Users/kirkehmsen/data/KE4/KE4-1-A07_S1159_L001_R1_001.fastq, 0.27492, 793
    /Users/kirkehmsen/data/KE4/KE4-1-A07_S1159_L001_R2_001.fastq, 0.27492, 793
    /Users/kirkehmsen/data/KE4/KE4-1-A08_S1160_L001_R1_001.fastq, 0.00208, 6
    /Users/kirkehmsen/data/KE4/KE4-1-A08_S1160_L001_R2_001.fastq, 0.00208, 6
    /Users/kirkehmsen/data/KE4/KE4-1-A09_S1161_L001_R1_001.fastq, 0.23332, 673
    /Users/kirkehmsen/data/KE4/KE4-1-A09_S1161_L001_R2_001.fastq, 0.23332, 673
    /Users/kirkehmsen/data/KE4/KE4-1-A10_S1162_L001_R1_001.fastq, 0.21145, 610
    /Users/kirkehmsen/data/KE4/KE4-1-A10_S1162_L001_R2_001.fastq, 0.21145, 610
    /Users/kirkehmsen/data/KE4/KE4-1-A11_S1163_L001_R1_001.fastq, 0.25657, 740
    /Users/kirkehmsen/data/KE4/KE4-1-A11_S1163_L001_R2_001.fastq, 0.25657, 740
```

```
File output information:
    Output directory: /Users/kirkehmsen/ImputedGenotypes_KE4
    Total file #: 7
    Total file output sizes:
        09182019_script_metrics.txt: 134.0 KB
        09182019_allele_definitions.txt: 7.8 MB
        09182019_fasta.fa: 1.9 MB
        09182019_allele_evidence.pdf: 111.2 MB
        09182019_blastn_alignments.txt: 108.5 MB
        09182019_population_summary.txt: 55.1 KB
        09182019_imputed_genotypes.txt: 7.8 MB

Script operation times:
    start time: 18:58:56
    fasta processing time: 0 hr|00 min|09 sec|471913 microsec
    alignments processing time: 0 hr|00 min|56 sec|204578 microsec
    imputation processing time: 0 hr|00 min|08 sec|779179 microsec
    frequency plots compilation time: 3 hr|16 min|12 sec|555270 microsec
    accessory file processing time: 0 hr|00 min|22 sec|004418 microsec
    total processing time: 3 hr|17 min|49 sec|169186 microsec
    end time: 22:16:46
```

**S8. File outputs of CollatedMotifs.py.**

(a) Example of custom-named directory (CollatedMotifs.py user *input* #1) populated with 5 output files and 3 sub-directories; filenames are automatically prefixed by script operations with system start date. ***fasta.fa, blastn_alignments.txt, markov_background.txt,*** and the three sub-directories are prepared during script operations and support script operations, but key script output content is contained in ***collated_TFBS.txt*** with script operation parameters logged in ***script_metrics.txt***.

(b) ***collated_TFBS.txt*** reports the top 5 ranked alleles for R1+R2 (for PE data), with allele frequency metrics in the '*Allele*' definition. '*Motifs*' indicates the total number of distinct TFBS identified in the allele, along with the number of distinct TFs that comprise these TFBS; '*Synopsis*' summarizes the # of lost sites and # of new TFBS relative to a reference sequence; '*Details'* summarizes the TF identities associated with the TFBS differences. *NEW motifs* are mapped above the allele alignment ('+ strand' indicating the sequence as reported, '- strand' indicating its implicit complement); *LOST motifs* are mapped below the allele alignment.

(c) ***script_metrics.txt*** logs script operation metadata.

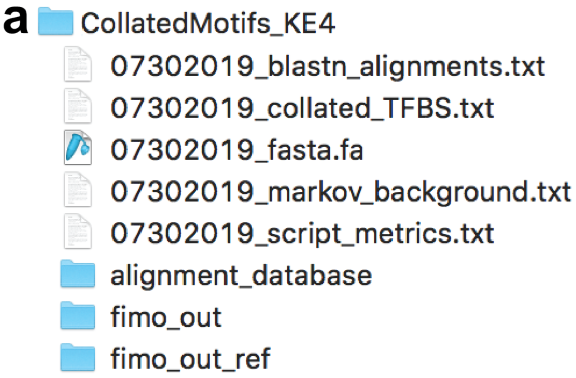


**b** *collated_TFBS.txt*

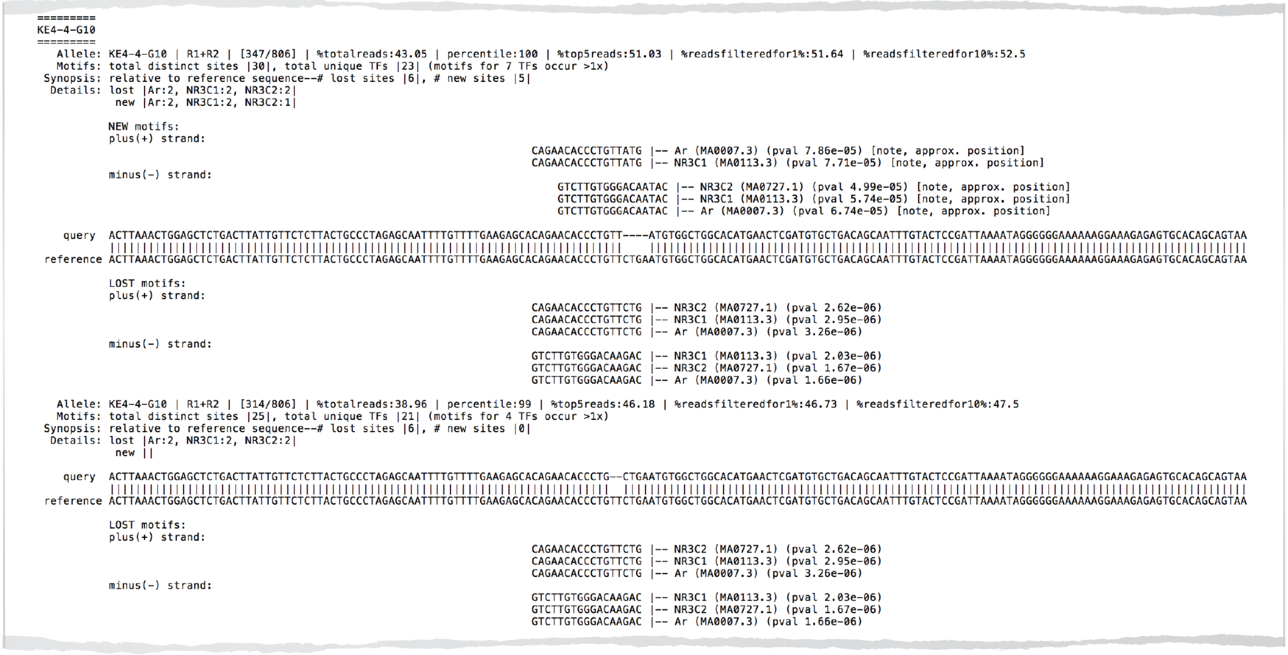

```
=========
KE4-4-G10
=========
   Allele: KE4-4-G10 | R1+R2 | [347/806] | %totalreads:43.05 | percentile:100 | %top5reads:51.03 | %readsfilteredfor1%:51.64 | %readsfilteredfor10%:52.5
   Motifs: total distinct sites |30|, total unique TFs |23| (motifs for 7 TFs occur >1x)
 Synopsis: relative to reference sequence--# lost sites |6|, # new sites |5|
  Details: lost |Ar:2, NR3C1:2, NR3C2:2|
            new |Ar:2, NR3C1:2, NR3C2:1|

           NEW motifs:
           plus(+) strand:
                                                                             CAGAACACCCTGTTATG |-- Ar (MA0007.3) (pval 7.86e-05) [note, approx. position]
                                                                             CAGAACACCCTGTTATG |-- NR3C1 (MA0113.3) (pval 7.71e-05) [note, approx. position]

           minus(-) strand:
                                                                                 GTCTTGTGGGACAATAC |-- NR3C2 (MA0727.1) (pval 4.99e-05) [note, approx. position]
                                                                                 GTCTTGTGGGACAATAC |-- NR3C1 (MA0113.3) (pval 5.74e-05) [note, approx. position]
                                                                                 GTCTTGTGGGACAATAC |-- Ar (MA0007.3) (pval 6.74e-05) [note, approx. position]

    query  ACTTAAACTGGAGCTCTGACTTATTGTTCTCTTACTGCCCTAGAGCAATTTTGTTTTGAAGAGCACAGAACACCCTGTT----ATGTGGCTGGCACATGAACTCGATGTGCTGACAGCAATTTGTACTCCGATTAAAATAGGGGGGAAAAAAGGAAAGAGAGTGCACAGCAGTAA
           |||||||||||||||||||||||||||||||||||||||||||||||||||||||||||||||||||||||||||||||    ||||||||||||||||||||||||||||||||||||||||||||||||||||||||||||||||||||||||||||||||||||||||||||
reference  ACTTAAACTGGAGCTCTGACTTATTGTTCTCTTACTGCCCTAGAGCAATTTTGTTTTGAAGAGCACAGAACACCCTGTTCTGAATGTGGCTGGCACATGAACTCGATGTGCTGACAGCAATTTGTACTCCGATTAAAATAGGGGGGAAAAAAGGAAAGAGAGTGCACAGCAGTAA

           LOST motifs:
           plus(+) strand:
                                                                             CAGAACACCCTGTTCTG |-- NR3C2 (MA0727.1) (pval 2.62e-06)
                                                                             CAGAACACCCTGTTCTG |-- NR3C1 (MA0113.3) (pval 2.95e-06)
                                                                             CAGAACACCCTGTTCTG |-- Ar (MA0007.3) (pval 3.26e-06)

           minus(-) strand:
                                                                             GTCTTGTGGGACAAGAC |-- NR3C1 (MA0113.3) (pval 2.03e-06)
                                                                             GTCTTGTGGGACAAGAC |-- NR3C2 (MA0727.1) (pval 1.67e-06)
                                                                             GTCTTGTGGGACAAGAC |-- Ar (MA0007.3) (pval 1.66e-06)

   Allele: KE4-4-G10 | R1+R2 | [314/806] | %totalreads:38.96 | percentile:99 | %top5reads:46.18 | %readsfilteredfor1%:46.73 | %readsfilteredfor10%:47.5
   Motifs: total distinct sites |25|, total unique TFs |21| (motifs for 4 TFs occur >1x)
 Synopsis: relative to reference sequence--# lost sites |6|, # new sites |0|
  Details: lost |Ar:2, NR3C1:2, NR3C2:2|
            new ||

    query  ACTTAAACTGGAGCTCTGACTTATTGTTCTCTTACTGCCCTAGAGCAATTTTGTTTTGAAGAGCACAGAACACCCTG--CTGAATGTGGCTGGCACATGAACTCGATGTGCTGACAGCAATTTGTACTCCGATTAAAATAGGGGGGAAAAAAGGAAAGAGAGTGCACAGCAGTAA
           |||||||||||||||||||||||||||||||||||||||||||||||||||||||||||||||||||||||||||||  ||||||||||||||||||||||||||||||||||||||||||||||||||||||||||||||||||||||||||||||||||||||||||||||||
reference  ACTTAAACTGGAGCTCTGACTTATTGTTCTCTTACTGCCCTAGAGCAATTTTGTTTTGAAGAGCACAGAACACCCTGTTCTGAATGTGGCTGGCACATGAACTCGATGTGCTGACAGCAATTTGTACTCCGATTAAAATAGGGGGGAAAAAAGGAAAGAGAGTGCACAGCAGTAA

           LOST motifs:
           plus(+) strand:
                                                                             CAGAACACCCTGTTCTG |-- NR3C2 (MA0727.1) (pval 2.62e-06)
                                                                             CAGAACACCCTGTTCTG |-- NR3C1 (MA0113.3) (pval 2.95e-06)
                                                                             CAGAACACCCTGTTCTG |-- Ar (MA0007.3) (pval 3.26e-06)

           minus(-) strand:
                                                                             GTCTTGTGGGACAAGAC |-- NR3C1 (MA0113.3) (pval 2.03e-06)
                                                                             GTCTTGTGGGACAAGAC |-- NR3C2 (MA0727.1) (pval 1.67e-06)
                                                                             GTCTTGTGGGACAAGAC |-- Ar (MA0007.3) (pval 1.66e-06)
```

#### C *script_metrics.txt*

```
CollatedMotifs.py: Script Metrics
Date: 07/30/2019

Operating system information:
    name: Kirks-MBP.lan
    platform: Darwin-17.7.0-x86_64-i386-64bit
    RAM (GB): 16.0
    physical CPU/effective CPU: 4/8
    executable: /Library/Frameworks/Python.framework/Versions/3.7/Resources/Python.app/Contents/MacOS/Pytho

User-entered variables:
    output_directory: /Users/kirkehmsen/CollatedMotifs_KE4
    fastq_directory: /Users/kirkehmsen/data/KE4
    fasta_ref: /Users/kirkehmsen/FKBP5_GOR+86.848kb_KE4.txt
    blastn_path: /usr/local/bin/blast/bin/blastn
    makeblastdb_path: /usr/local/bin/blast/bin/makeblastdb
    db_prefix: FKBP5_GOR+86.848kb
    fimo_path: /Users/kirkehmsen/Meme/bin/fimo
    fimo_motifs_path: /Users/kirkehmsen/Meme/motif_databases/JASPAR/JASPAR_CORE_2016_vertebrates.meme
    fasta_get_markov_path: /Users/kirkehmsen/Meme/meme-5.0.1/src/fasta-get-markov
    markov_background_file: /Users/kirkehmsen/hg38.fa

fastq file information:
    Illumina sequencing run ID(s): @M00582:216
    Number of fastq files processed: 768
    Size distribution of fastq files processed:
        total... 197 MB
        range... max: 0.61 MB; min: 0.00069 MB; median: 0.261 MB; mean +/- stdev: 0.257 +/- 0.132 MB
    Read distribution within fastq files to process:
        total... 569,374 reads
        range... max: 1745 reads; min: 2 reads; median: 754.0 reads; mean +/- stdev: 741.0 +/- 382.0 reads

fastq files processed (name, size (MB), reads):
    /Users/kirkehmsen/data/KE4/KE4-1-A01_S1153_L001_R1_001.fastq, 0.13484, 389
    /Users/kirkehmsen/data/KE4/KE4-1-A01_S1153_L001_R2_001.fastq, 0.13484, 389
    /Users/kirkehmsen/data/KE4/KE4-1-A02_S1154_L001_R1_001.fastq, 0.26033, 751
    /Users/kirkehmsen/data/KE4/KE4-1-A02_S1154_L001_R2_001.fastq, 0.26033, 751

Reference sequences provided in fasta_ref file:
    >KE4
    ACTTAAACTGGAGCTCTGACTTATTGTTCTCTTACTGCCCTAGAGCAATTTTGTTTTGAAGAGCACAGAACACCCTGTTCTGAATGTGGCTGGCACATGAACTCGATGTGCTGACAGCAATTTGTACTCCGATTAAAATAGGGGGGAAAAAAGGAAAGAGAGTGCACAGCAGTAAC

# of TFBS motifs examined: 519
Identities of TFBS motifs examined:
    ALX3      ARNT::HIF1A    ATF4        ATF7       Ahr::Arnt     Alx1       Alx4        Ar
   Arid3a       Arid3b      Arid5a       Arnt         Arntl        Arx       Ascl2       Atf1
    Atf3        Atoh1       BARHL2      BARX1         BATF3     BATF::JUN    BCL6B     BHLHE22
  BHLHE23      BHLHE40     BHLHE41       BSX      Bach1::Mafk   Barhl1       Bcl6     Bhlha15
    CDX1         CDX2        CEBPA      CEBPB         CEBPD       CEBPE      CEBPG       CENPB
    CLOCK        CREB1       CREB3     CREB3L1        CTCF        CUX1       CUX2      Creb3l2
    Creb5        Crem         Crx        DBP          DLX6       DMRT3      DUX4        DUXA
Ddit3::Cebpa     Dlx1        Dlx2       Dlx3          Dlx4       Dmbx1       Dux        E2F1
    E2F2         E2F3        E2F4       E2F6          E2F7       E2F8       EBF1        EGR1
    EGR2         EGR3        EGR4        EHF          ELF1       ELF3       ELF4        ELF5
    ELK1         ELK3        ELK4       EMX1          EMX2        EN1        EN2       EOMES
     ERF          ERG        ESR1       ESR2          ESRRB      ESX1       ETS1        ETV1
    ETV2         ETV3        ETV4       ETV5          ETV6       EVX1       EVX2    EWSR1-FLI1
    Esrra        Esrrg        FEV       FIGLA         FLI1        FOS      FOS::JUN     FOSL1
    FOSL2        FOXA1       FOXB1      FOXC1         FOXC2      FOXD1      FOXD2       FOXF2
    FOXG1        FOXH1       FOXI1      FOXL1         FOXO3      FOXO4      FOXO6       FOXP1
    FOXP2        FOXP3       Foxa2      Foxd3         Foxj2      Foxj3      Foxk1       Foxo1
 [illegible]  GATA1::TAL1    GATA2      GATA3      [illegible] [illegible] [illegible] [illegible]

 File output information:
    Output directory: /Users/kirkehmsen/CollatedMotifs_KE4
    Total file #: 5
    Total file output sizes: 320.0 bytes
    07292019_markov_background.txt: 223.0 bytes
    07292019_fasta.fa: 564.7 KB
    07292019_blastn_alignments.txt: 3.5 MB
    07292019_script_metrics.txt: 141.7 KB
    07292019_collated_TFBS.txt: 1.4 MB


 Script operation times:
    start time: 23:14:12
    makeblastdb and fasta-get-markov processing time: 0 hr|00 min|18 sec|846692 microsec
    fasta processing time: 0 hr|00 min|25 sec|742864 microsec
    alignments processing time: 0 hr|00 min|01 sec|536980 microsec
    allele definitions processing time: 0 hr|00 min|00 sec|908172 microsec
    TFBS processing time (FIMO): 0 hr|02 min|45 sec|710900 microsec
    TFBS collation processing time: 0 hr|00 min|01 sec|133114 microsec
    total processing time: 0 hr|03 min|34 sec|016575 microsec
    end time: 23:17:46
```

**S9. *FKBP5* GOR conservation across 100 vertebrates.**
PhyloP (blue) and phastCons (green) tracks for *FKBP5* (a) intronic and (b) upstream GORs defined in A549. GBS matches identified by RSAT matrix-scan or by 'short match' to generic GBS motif 5'-NNN**ACA**NNNN**GT**N**C**NN-3' in UCSC Genome Browser are indicated.

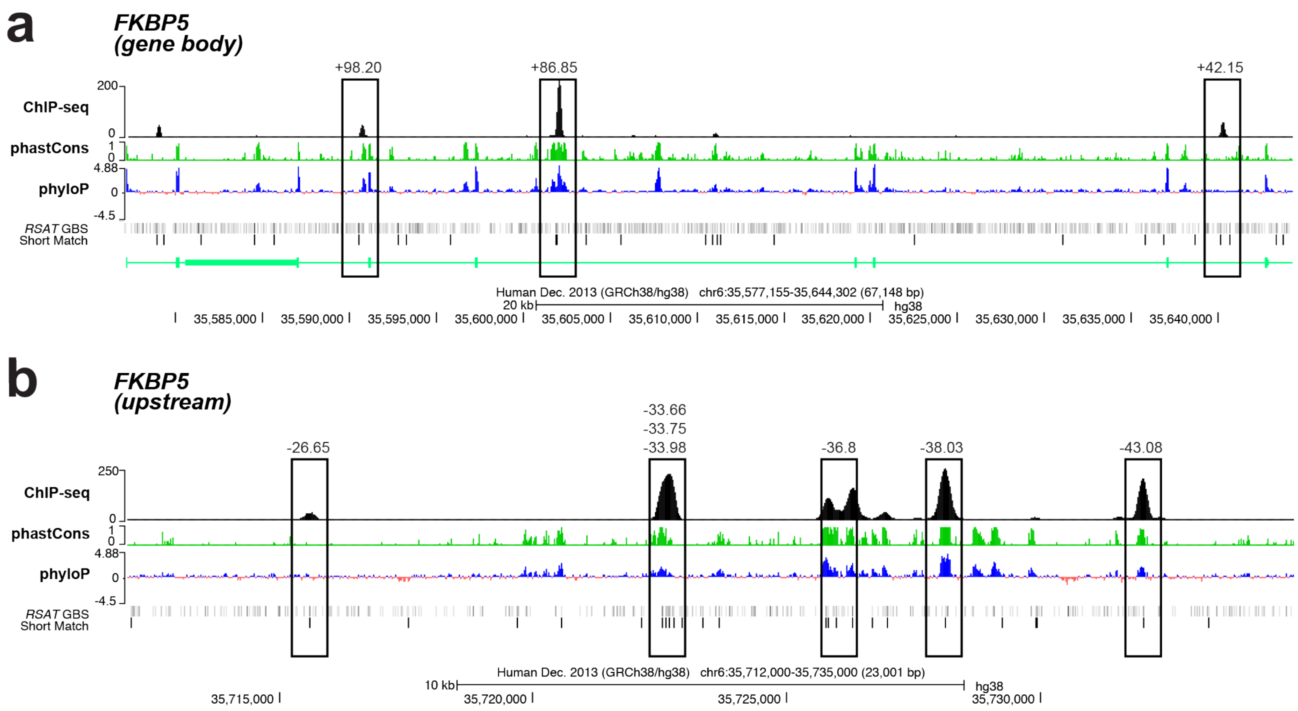

**S10. Procedure for automated delivery of i7 and i5 primers to 384-well plates for PCR2, using the Echo® liquid handler (Labcyte, San Jose, CA). The format illustrated preserves barcode relationships assigned to samples in 96-well format, and as elaborated in an Illumina® Sample Sheet using SampleSheet.py.**

(a) Index oligo stocks can be transferred from 96-well source plates (typical source format from oligo supplier) to 384-well plates and adjusted to a practical stock concentration, using a 12-channel pipet. In this example, i5 oligos are arrayed in rows 1-4 of a Labcyte Echo®-qualified 384-well polypropylene source plate (well IDs and numbers correspond to ID in 96-well plate), and i7 oligos are arrayed in rows 9-12 (in up to 65 μL volumes, plates sealed with adhesive foil and stored at -20 °C). Preparing the index oligos in 384-well format facilitates convenient, automated transfer of oligos to destination wells as defined in Labcyte Plate Reformat software;

(b) Example workflow for PCR1 & PCR2 setup in 384-well plates. Here, four source plates (Plates 1-4) are indicated. (1) *Master Mix* for PCR1 is delivered using a multi-channel pipet to wells of a 384-well PCR plate (*labeled #1*); (2) Appropriate volume of lysate is transferred from 96-well source plates to wells of a 384-well plate, enabling content from four 96-well plates to be arrayed into a single 384-well plate to expedite processing; (3) in a separate 384-well PCR plate (*labeled #2*);, PCR2 Master Mix can be delivered by hand (12-channel pipet) or by Echo (*e.g.,* from 6-reservoir source plate); (4) *i7 and i5 indices can be delivered in small volumes to target wells using Echo acoustic technology (i5 across individual groups of four rows that correspond to a single 96-well plate; i7 to individual wells within a block of four rows);* (5) finally, PCR2 template (small volume of PCR1 template) can be delivered by hand (12-channel pipet) from PCR1 plate to PCR2 plate.

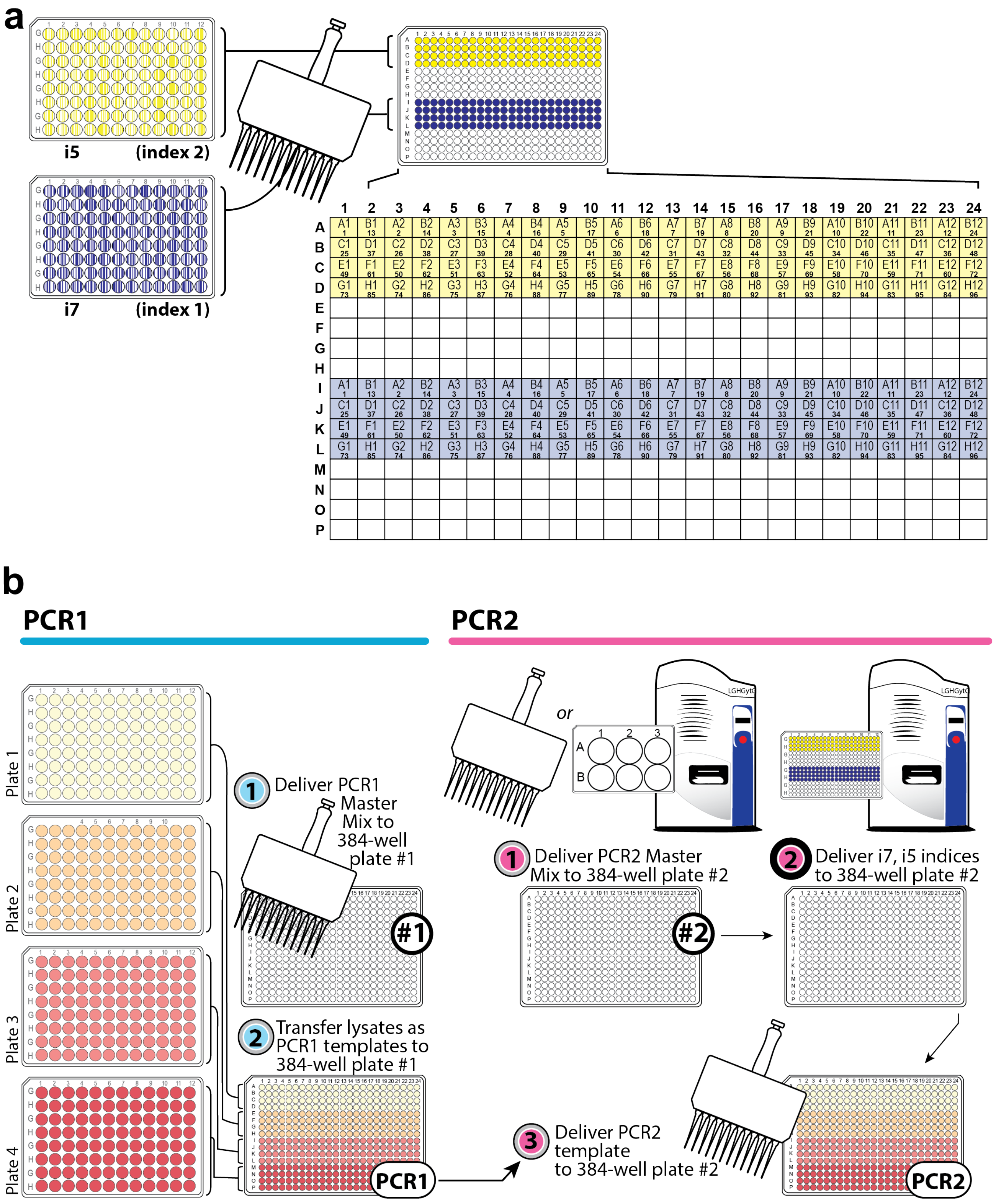
a
i5
(index 2)
i7
(index 1)
1 2 3 4 5 6 7 8 9 10 11 12 13 14 15 16 17 18 19 20 21 22 23 24
A1 1 B1 13 A2 2 B2 14 A3 3 B3 15 A4 4 B4 16 A5 5 B5 17 A6 6 B6 18 A7 7 B7 19 A8 8 B8 20 A9 9 B9 21 A10 10 B10 22 A11 11 B11 23 A12 12 B12 24
C1 25 D1 37 C2 26 D2 38 C3 27 D3 39 C4 28 D4 40 C5 29 D5 41 C6 30 D6 42 C7 31 D7 43 C8 32 D8 44 C9 33 D9 45 C10 34 D10 46 C11 35 D11 47 C12 36 D12 48
E1 49 F1 61 E2 50 F2 62 E3 51 F3 63 E4 52 F4 64 E5 53 F5 65 E6 54 F6 66 E7 55 F7 67 E8 56 F8 68 E9 57 F9 69 E10 58 F10 70 E11 59 F11 71 E12 60 F12 72
G1 73 H1 85 G2 74 H2 86 G3 75 H3 87 G4 76 H4 88 G5 77 H5 89 G6 78 H6 90 G7 79 H7 91 G8 80 H8 92 G9 81 H9 93 G10 82 H10 94 G11 83 H11 95 G12 84 H12 96
b
PCR1
PCR2
Plate 1
Plate 2
Plate 3
Plate 4
1 Deliver PCR1 Master Mix to 384-well plate #1
#1
2 Transfer lysates as PCR1 templates to 384-well plate #1
PCR1
or
1 Deliver PCR2 Master Mix to 384-well plate #2
#2
2 Deliver i7, i5 indices to 384-well plate #2
3 Deliver PCR2 template to 384-well plate #2
PCR2

**S11. BaseSpace Sequencing Hub metadata for files processed in use case.** Screenshots of metadata preserved in BaseSpace Sequencing Hub, https://basespace.illumina.com/run/36555528/171004-M00582-0217-000000000-AV3NW. 9,216 fastq files were demultiplexed from sequencing RunID: 171004_M00582_0217_000000000-AV3NW (10-04-2017).

#### General Info

| | |
|---|---|
| Run Status | Complete |
| Lane QC Status | QC Passed |
| Flowcell ID | 000000000-AV3NW |
| Run ID | 171004_M00582_0217_000000000-AV3NW |
| Instrument Name | M00582 |
| Instrument Type | MiSeq |
| %PF | |
| Avg %Q30 | 92.14% |
| Yield | 88.45% |
| Cycles | 2.53242 bp |
| Created | 151 \| 8 \| 8 \| 151 |
| Owner | 2017-10-04 11:32 |
| User | Kirk Ehmsen |
| File Count/Size | Kirk Ehmsen |
| | 9.796 files (4 GB) |

Per Lane Metrics

Columns

| LANE | READ | CLUSTER PF (%) | %≥Q30 | YIELD | ERROR RATE% | READS PF | DENSITY | TILES | LEGACY PHAS/PREPHAS (%) | INTENSITY | COMMENTS | STATUS |
|---|---|---|---|---|---|---|---|---|---|---|---|---|
| 1 | 1 | 92.15 ±1.95 | 93.93 | 1.21 Gbp | 0.62 ±0.10 | 8,065,046 | 449 ±11 | 28 | 0.155 / 0.396 | 121 ±12 | | QC Passed |
| | 2(I) | | 75.65 | 56.46 Mbp | 0.00 ±0.00 | | | | 0.000 / 0.000 | 343 ±43 | | |
| | 3(I) | | 95.51 | 56.46 Mbp | 0.00 ±0.00 | | | | 0.000 / 0.000 | 317 ±40 | | |
| | 4 | | 83.24 | 1.21 Gbp | 0.73 ±0.08 | | | | 0.250 / 0.373 | 91 ±11 | | |


Per Lane Metrics

Columns

| LANE | READ | CLUSTER PF (%) | %≥Q30 | YIELD | ERROR RATE% | READS PF | DENSITY | TILES | LEGACY PHAS/PREPHAS (%) | INTENSITY | COMMENTS | STATUS |
|---|---|---|---|---|---|---|---|---|---|---|---|---|
| 1 | 1 | 92.15 ±1.95 | 93.93 | 1.21 Gbp | 0.62 ±0.10 | 8,065,046 | 449 ±11 | 28 | 0.155 / 0.396 | 121 ±12 | | QC Passed |
| | 2(I) | | 75.65 | 56.46 Mbp | 0.00 ±0.00 | | | | 0.000 / 0.000 | 343 ±43 | | |
| | 3(I) | | 95.51 | 56.46 Mbp | 0.00 ±0.00 | | | | 0.000 / 0.000 | 317 ±40 | | |
| | 4 | | 83.24 | 1.21 Gbp | 0.73 ±0.08 | | | | 0.250 / 0.373 | 91 ±11 | | |


| TOTAL READS | PF READS | % READS IDENTIFIED (PF) | CV | MIN | MAX |
|---|---|---|---|---|---|
| 17,505,188 | 16,130,092 | 75.3308 | 0.6960 | 0.0000 | 0.0332 |

**S12. Barcode distributions among sequenced reads.**

Among 9,216 barcode pairs used to uniquely label amplicons, 99.4% (9,164) were identified as linked to reads by sequencing on an Illumina® MiSeq instrument. Barcode pairs not identified (52, <0.6%) are suspected to have been assigned to wells with insufficient or no template source.

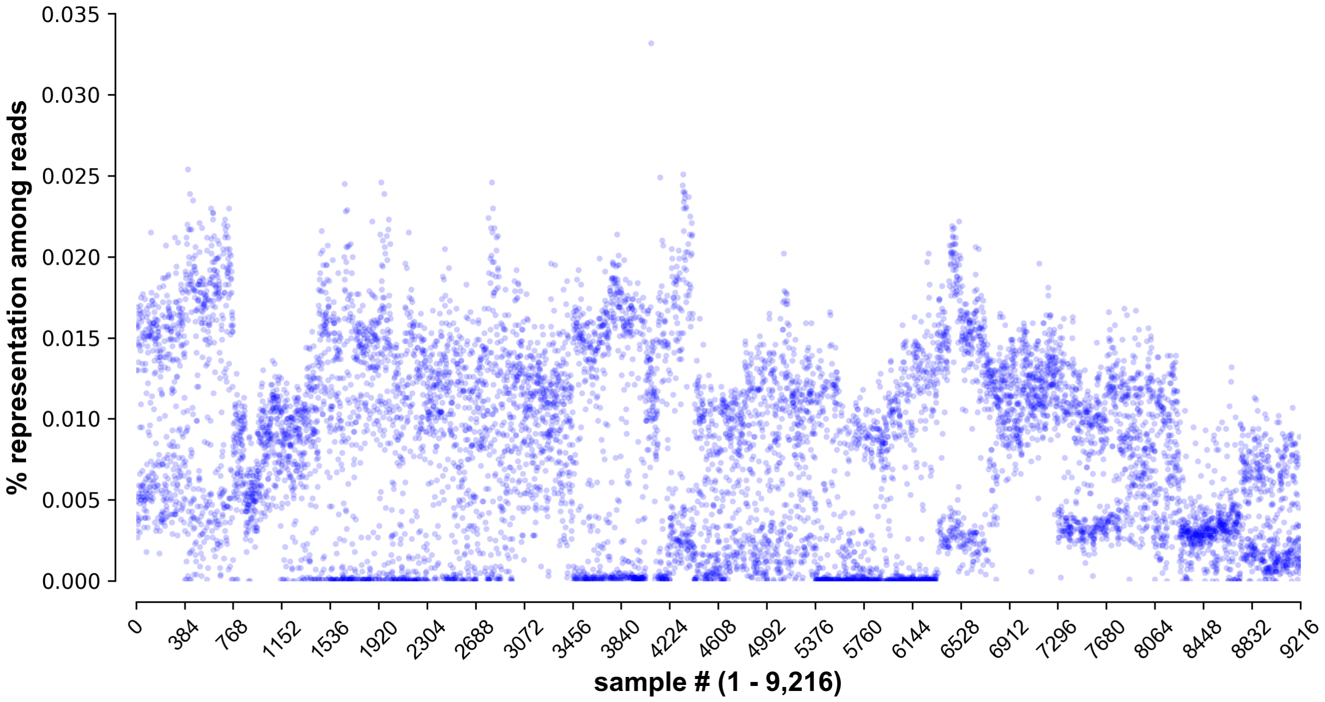

**S13. Selected mutant GORs: disruption of individual peak summit GBS motif(s) at two GR ChIP-seq peaks active in a luciferase assay (Supp. Fig. 14) is not sufficient to impact *FKBP5* dex induction.** Examples of regulatory analysis and interpretation informed by gain/loss of TFBS (CollatedMotifs.py).

(a) **GOR 2: +86.85 kb** (intronic) peak with high vertebrate conservation (**Supp. Fig. 9**)
A biallelic deletion (two distinct deletion alleles) in two independently isolated clones (4-4 G02, *clone #1* & 4-4 G10, *clone #2*: identical genotypes) causes loss of native *GBS +86.85 kb* (5'-CAGAACACCCTGTTCT-3', maroon arrows), but a *novel GBS motif match* (red arrows) is restored in one allele. *Clone #1* (4-4 G02) exhibits no evident loss of *FKBP5* transcript induction in dex; *clone #2* (4-4 G10) exhibits slightly elevated *FKBP5* transcript induction at 1 nM and 100 nM dex relative to wt (attributed to a lower basal transcript level relative to wild-type (EtOH), which reaches wild-type induced levels as quantified as deltaCT). CollatedMotifs.py was run with default p-value setting, <0.0001.

(b) **GOR 5: -33.61 kb, -33.75 kb, -33.99 kb** (upstream) peak with high vert. conservation.
This ChIP-seq peak is characterized by multiple high-scoring (low p-value) GBS matches near its summit:

- *GBS -33.61 kb*—GBS disruption appears to slightly increase *FKBP5* transcript induction (<2-fold) in two examined clones (12-3 B09, *clone #1* & 12-5 G05, *clone #3*), primarily *via* increased transcript levels at 100 nM dex; a third examined clone (12-5 C04, *clone #2*), however, exhibits slightly blunted *FKBP5* transcript induction (<2-fold) *via* moderately increased basal levels and reduced induction at 100 nM dex. Examination of unlinked dex-responsive loci (*ANKRD1* on chr10*, PER1* on chr17*, SCNN1A* on chr12*, IL8* on chr4 *vs. FKBP5* on chr. 6) across these clones, however, demonstrates that weak fold-change differences are not unique to *FKBP5* and may relate instead to moderate clonal or experimental variation (supported by examination of 13-4 B01, *clone #4*, which also exhibits slight increase in *FKBP5* transcript induction, with small homozygous deletion *outside* GBS). CollatedMotifs.py was run with default p-value setting, <0.0001.
- *GBS -33.75 kb*—GBS disruption appears to slightly increase *FKBP5* transcript induction (~2-fold) in one examined clone (14-2 B09, *clone #1* & 14-5 B05, *clone #3*) *via* increased transcript levels at 100 nM dex, but not in a third examined clone (14-4 E08, *clone #2*); native summit GBS is ablated in all three clones, but mutation in *clones #2 & #3* re-populate distinct alternative GBS matches. CollatedMotifs.py was run with default p-value setting, <0.001; screenshot from *collated_TFBS.txt* file shows subset of 'new' and 'lost' motifs for *clone #3* (edited for space here).

- *GBS -33.99 kb*—GBS disruption does not appear to affect *FKBP5* transcript basal level or induction. CollatedMotifs.py was run with default p-value setting, <0.001.

*Note:* For *GBS -33.75 kb* & *GBS -33.99 kb,* CollatedMotifs.ipynb was manually edited as indicated below (' --thresh 1e-3') prior to operation, to account for *GBS -33.75 kb* & *GBS -33.99 kb* p-values below the FIMO default threshold (default p-value = 0.0001),

```
#Reference sequence(s): FIMO command (usage: fimo --bfile <background file> <motif file> <sequence file>)

cmd_TFBS = str(fimo_path)+' --bfile '+str(markovbackground_output)+' --o '+str(ref_TFBS_output)+
' --thresh 1e-3'+' '+str(fimo_motifs_path)+' '+str(ref_input)

os.system(cmd_TFBS)


#Alleles: FIMO command (usage: fimo --bfile <background file> <motif file> <sequence file>)

cmd_TFBS = str(fimo_path)+' --bfile '+str(markovbackground_output)+' --o '+str(allele_TFBS_output)+
' --thresh 1e-3'+' '+str(fimo_motifs_path)+' '+str(query_input)

os.system(cmd_TFBS)
```

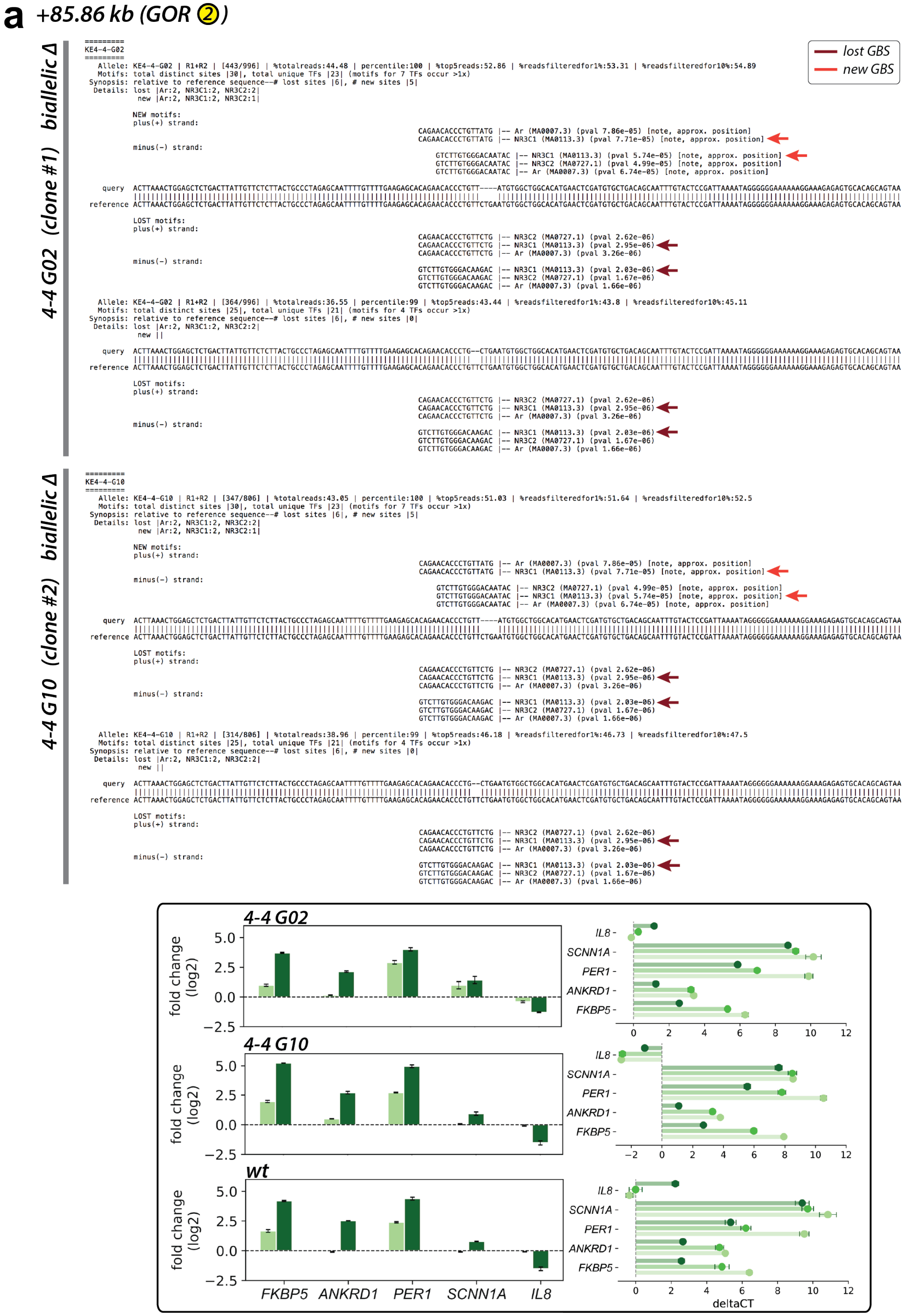
a
+85.86 kb (GOR 2)
lost GBS
new GBS
4-4 G02 (clone #1) biallelic Δ
KE4-4-G02
Allele: KE4-4-G02 | R1+R2 | [443/996] | %totalreads:44.48 | percentile:100 | %top5reads:52.86 | %readsfilteredfor1%:53.31 | %readsfilteredfor10%:54.89
Motifs: total distinct sites |30|, total unique TFs |23| (motifs for 7 TFs occur >1x)
Synopsis: relative to reference sequence--# lost sites |6|, # new sites |5|
Details: lost |Ar:2, NR3C1:2, NR3C2:2|
new |Ar:2, NR3C1:2, NR3C2:1|
NEW motifs:
plus(+) strand:
CAGAACACCCTGTTATG |-- Ar (MA0007.3) (pval 7.86e-05) [note, approx. position]
CAGAACACCCTGTTATG |-- NR3C1 (MA0113.3) (pval 7.71e-05) [note, approx. position]
minus(-) strand:
GTCTTGTGGGACAATAC |-- NR3C1 (MA0113.3) (pval 5.74e-05) [note, approx. position]
GTCTTGTGGGACAATAC |-- NR3C2 (MA0727.1) (pval 4.99e-05) [note, approx. position]
GTCTTGTGGGACAATAC |-- Ar (MA0007.3) (pval 6.74e-05) [note, approx. position]
query
reference
LOST motifs:
plus(+) strand:
CAGAACACCCTGTTCTG |-- NR3C2 (MA0727.1) (pval 2.62e-06)
CAGAACACCCTGTTCTG |-- NR3C1 (MA0113.3) (pval 2.95e-06)
CAGAACACCCTGTTCTG |-- Ar (MA0007.3) (pval 3.26e-06)
minus(-) strand:
GTCTTGTGGGACAAGAC |-- NR3C1 (MA0113.3) (pval 2.03e-06)
GTCTTGTGGGACAAGAC |-- NR3C2 (MA0727.1) (pval 1.67e-06)
GTCTTGTGGGACAAGAC |-- Ar (MA0007.3) (pval 1.66e-06)
Allele: KE4-4-G02 | R1+R2 | [364/996] | %totalreads:36.55 | percentile:99 | %top5reads:43.44 | %readsfilteredfor1%:43.8 | %readsfilteredfor10%:45.11
Motifs: total distinct sites |25|, total unique TFs |21| (motifs for 4 TFs occur >1x)
Synopsis: relative to reference sequence--# lost sites |6|, # new sites |0|
Details: lost |Ar:2, NR3C1:2, NR3C2:2|
new ||
4-4 G10 (clone #2) biallelic Δ
KE4-4-G10
Allele: KE4-4-G10 | R1+R2 | [347/806] | %totalreads:43.05 | percentile:100 | %top5reads:51.03 | %readsfilteredfor1%:51.64 | %readsfilteredfor10%:52.5
Motifs: total distinct sites |30|, total unique TFs |23| (motifs for 7 TFs occur >1x)
Synopsis: relative to reference sequence--# lost sites |6|, # new sites |5|
Details: lost |Ar:2, NR3C1:2, NR3C2:2|
new |Ar:2, NR3C1:2, NR3C2:1|
Allele: KE4-4-G10 | R1+R2 | [314/806] | %totalreads:38.96 | percentile:99 | %top5reads:46.18 | %readsfilteredfor1%:46.73 | %readsfilteredfor10%:47.5
Motifs: total distinct sites |25|, total unique TFs |21| (motifs for 4 TFs occur >1x)
Synopsis: relative to reference sequence--# lost sites |6|, # new sites |0|
Details: lost |Ar:2, NR3C1:2, NR3C2:2|
new ||
4-4 G02
4-4 G10
wt
fold change (log2)
FKBP5
ANKRD1
PER1
SCNN1A
IL8
deltaCT

**b** *-33.61, -33.75, -33.99 kb (GOR ⑤)*

GBS -33.61 kb

lost GBS

***12-3 B09 (clone #1) biallelic Δ***

```
==========
KE12-3-B09
==========
    Allele: KE12-3-B09 | R1+R2 | [204/490] | %totalreads:41.63 | percentile:100 | %top5reads:49.76 | %readsfilteredfor1%:50.87 | %readsfilteredfor10%:50.87
    Motifs: total distinct sites |45|, total unique TFs |44| (motifs for 1 TFs occur >1x)
  Synopsis: relative to reference sequence--# lost sites |6|, # new sites |3|
   Details: lost |Ar:2, NR3C1:2, NR3C2:2|
             new |CEBPA:1, Rarb:1, Rarg:1|

            NEW motifs:
            plus(+) strand:
                                                                              CTTGCACCATC |-- CEBPA (MA0102.3) (pval 4.42e-05) [note, approx. position]
            minus(-) strand:
                                                                                          CAGTTGAGAACGTGGT |-- Rarg (MA0859.1) (pval 3.95e-05) [note, approx. position]
                                                                                           AGTTGAGAACGTGGTA |-- Rarb (MA0857.1) (pval 7.16e-05) [note, approx. position]

     query GCCCTGAAGCTCCCGGGCTGCAGGGAACCATGAGGCTGCCTGGGCTTCCGGCTGCTGCAGGATAAAGGTCAACTCT------------------TGCACCATCTGGGCTTGCCTGCCTCTGGCCTCATCTCCCCGGGCTCTCCAACCTGCACTCCATGTCCAGCCAGACCAAACA
           ||||||||||||||||||||||||||||||||||||||||||||||||||||||||||||||||||||||||||||                  |||||||||||||||||||||||||||||||||||||||||||||||||||||||||||||||||||||||||||||||||
 reference GCCCTGAAGCTCCCGGGCTGCAGGGAACCATGAGGCTGCCTGGGCTTCCGGCTGCTGCAGGATAAAGGTCAACTCTTGCTGGAACACGAGGTTCTGCACCATCTGGGCTTGCCTGCCTCTGGCCTCATCTCCCCGGGCTCTCCAACCTGCACTCCATGTCCAGCCAGACCAAACA

            LOST motifs:
            plus(+) strand:
                                                                                   TGGAACACGAGGTTCTG |-- NR3C2 (MA0727.1) (pval 1.33e-05)
                                                                                   TGGAACACGAGGTTCTG |-- Ar (MA0007.3) (pval 1.52e-05)
                                                                                   TGGAACACGAGGTTCTG |-- NR3C1 (MA0113.3) (pval 8.43e-06)
            minus(-) strand:
                                                                                   ACCTTGTGCTCCAAGAC |-- Ar (MA0007.3) (pval 1.23e-05)
                                                                                   ACCTTGTGCTCCAAGAC |-- NR3C2 (MA0727.1) (pval 2.36e-05)
                                                                                   ACCTTGTGCTCCAAGAC |-- NR3C1 (MA0113.3) (pval 1.79e-05)

    Allele: KE12-3-B09 | R1+R2 | [197/490] | %totalreads:40.2 | percentile:98 | %top5reads:48.05 | %readsfilteredfor1%:49.13 | %readsfilteredfor10%:49.13
    Motifs: total distinct sites |44|, total unique TFs |44| (motifs for 0 TFs occur >1x)
  Synopsis: relative to reference sequence--# lost sites |6|, # new sites |2|
   Details: lost |Ar:2, NR3C1:2, NR3C2:2|
             new |HIC2:1, Mitf:1|

            NEW motifs:
            plus(+) strand:
                                                                                     GGGCACGAGG |-- Mitf (MA0620.1) (pval 6.76e-05) [note, approx. position]
            minus(-) strand:
                                                                                   CGACCCGTG |-- HIC2 (MA0738.1) (pval 1e-05) [note, approx. position]

     query GCCCTGAAGCTCCCGGGCTGCAGGGAACCATGAGGCTGCCTGGGCTTCCGGCTGCTGCAGGATAAAGGTCAACTCTTGCTGGG-CACGAGGTTCTGCACCATCTGGGCTTGCCTGCCTCTGGCCTCATCTCCCCGGGCTCTCCAACCTGCACTCCATGTCCAGCCAGACCAAACA
           ||||||||||||||||||||||||||||||||||||||||||||||||||||||||||||||||||||||||||||||||||  |||||||||||||||||||||||||||||||||||||||||||||||||||||||||||||||||||||||||||||||||||||||||||||||
 reference GCCCTGAAGCTCCCGGGCTGCAGGGAACCATGAGGCTGCCTGGGCTTCCGGCTGCTGCAGGATAAAGGTCAACTCTTGCTGGAACACGAGGTTCTGCACCATCTGGGCTTGCCTGCCTCTGGCCTCATCTCCCCGGGCTCTCCAACCTGCACTCCATGTCCAGCCAGACCAAACA

            LOST motifs:
            plus(+) strand:
                                                                                   TGGAACACGAGGTTCTG |-- NR3C2 (MA0727.1) (pval 1.33e-05)
                                                                                   TGGAACACGAGGTTCTG |-- Ar (MA0007.3) (pval 1.52e-05)
                                                                                   TGGAACACGAGGTTCTG |-- NR3C1 (MA0113.3) (pval 8.43e-06)
            minus(-) strand:
                                                                                   ACCTTGTGCTCCAAGAC |-- Ar (MA0007.3) (pval 1.23e-05)
                                                                                   ACCTTGTGCTCCAAGAC |-- NR3C2 (MA0727.1) (pval 2.36e-05)
                                                                                   ACCTTGTGCTCCAAGAC |-- NR3C1 (MA0113.3) (pval 1.79e-05)
```

***12-5 C04 (clone #2) homozygous Δ***

```
==========
KE12-5-C04
==========
    Allele: KE12-5-C04 | R1+R2 | [379/484] | %totalreads:78.31 | percentile:100 | %top5reads:94.04 | %readsfilteredfor1%:96.19 | %readsfilteredfor10%:100.0
    Motifs: total distinct sites |42|, total unique TFs |42| (motifs for 0 TFs occur >1x)
  Synopsis: relative to reference sequence--# lost sites |6|, # new sites |0|
   Details: lost |Ar:2, NR3C1:2, NR3C2:2|
             new ||

     query GCCCTGAAGCTCCCGGGCTGCAGGGAACCATGAGGCTGCCTGGGCTTCCGGCTGCTGCAGGATAAAGGTCAACTCTTGCTGGAACCACGAGGTTCTGCACCATCTGGGCTTGCCTGCCTCTGGCCTCATCTCCCCGGGCTCTCCAACCTGCACTCCATGTCCAGCCAGACCAAACA
           |||||||||||||||||||||||||||||||||||||||||||||||||||||||||||||||||||||||||||||||||||| |||||||||||||||||||||||||||||||||||||||||||||||||||||||||||||||||||||||||||||||||||||||||||
 reference GCCCTGAAGCTCCCGGGCTGCAGGGAACCATGAGGCTGCCTGGGCTTCCGGCTGCTGCAGGATAAAGGTCAACTCTTGCTGGAA-CACGAGGTTCTGCACCATCTGGGCTTGCCTGCCTCTGGCCTCATCTCCCCGGGCTCTCCAACCTGCACTCCATGTCCAGCCAGACCAAACA

            LOST motifs:
            plus(+) strand:
                                                                                    TGGAACACGAGGTTCTG |-- NR3C2 (MA0727.1) (pval 1.33e-05) [note, approx. position]
                                                                                    TGGAACACGAGGTTCTG |-- Ar (MA0007.3) (pval 1.52e-05) [note, approx. position]
                                                                                    TGGAACACGAGGTTCTG |-- NR3C1 (MA0113.3) (pval 8.43e-06) [note, approx. position]
            minus(-) strand:
                                                                                    ACCTTGTGCTCCAAGAC |-- Ar (MA0007.3) (pval 1.23e-05) [note, approx. position]
                                                                                    ACCTTGTGCTCCAAGAC |-- NR3C2 (MA0727.1) (pval 2.36e-05) [note, approx. position]
                                                                                    ACCTTGTGCTCCAAGAC |-- NR3C1 (MA0113.3) (pval 1.79e-05) [note, approx. position]
```

***12-5 G05 (clone #3) homozygous Δ***

```
==========
KE12-5-G05
==========
    Allele: KE12-5-G05 | R1+R2 | [311/418] | %totalreads:74.4 | percentile:100 | %top5reads:93.96 | %readsfilteredfor1%:96.58 | %readsfilteredfor10%:100.0
    Motifs: total distinct sites |45|, total unique TFs |44| (motifs for 1 TFs occur >1x)
  Synopsis: relative to reference sequence--# lost sites |6|, # new sites |3|
   Details: lost |Ar:2, NR3C1:2, NR3C2:2|
             new |CEBPA:1, Rarb:1, Rarg:1|

            NEW motifs:
            plus(+) strand:
                                                                              CTTGCACCATC |-- CEBPA (MA0102.3) (pval 4.42e-05) [note, approx. position]
            minus(-) strand:
                                                                                          CAGTTGAGAACGTGGT |-- Rarg (MA0859.1) (pval 3.95e-05) [note, approx. position]
                                                                                           AGTTGAGAACGTGGTA |-- Rarb (MA0857.1) (pval 7.16e-05) [note, approx. position]

     query GCCCTGAAGCTCCCGGGCTGCAGGGAACCATGAGGCTGCCTGGGCTTCCGGCTGCTGCAGGATAAAGGTCAACTCT------------------TGCACCATCTGGGCTTGCCTGCCTCTGGCCTCATCTCCCCGGGCTCTCCAACCTGCACTCCATGTCCAGCCAGACCAAACA
           ||||||||||||||||||||||||||||||||||||||||||||||||||||||||||||||||||||||||||||                  |||||||||||||||||||||||||||||||||||||||||||||||||||||||||||||||||||||||||||||||||
 reference GCCCTGAAGCTCCCGGGCTGCAGGGAACCATGAGGCTGCCTGGGCTTCCGGCTGCTGCAGGATAAAGGTCAACTCTTGCTGGAACACGAGGTTCTGCACCATCTGGGCTTGCCTGCCTCTGGCCTCATCTCCCCGGGCTCTCCAACCTGCACTCCATGTCCAGCCAGACCAAACA

            LOST motifs:
            plus(+) strand:
                                                                                   TGGAACACGAGGTTCTG |-- NR3C2 (MA0727.1) (pval 1.33e-05)
                                                                                   TGGAACACGAGGTTCTG |-- Ar (MA0007.3) (pval 1.52e-05)
                                                                                   TGGAACACGAGGTTCTG |-- NR3C1 (MA0113.3) (pval 8.43e-06)
            minus(-) strand:
                                                                                   ACCTTGTGCTCCAAGAC |-- Ar (MA0007.3) (pval 1.23e-05)
                                                                                   ACCTTGTGCTCCAAGAC |-- NR3C2 (MA0727.1) (pval 2.36e-05)
                                                                                   ACCTTGTGCTCCAAGAC |-- NR3C1 (MA0113.3) (pval 1.79e-05)
```

***13-4 B01 (clone #4) homozygous Δ***

```
==========
KE13-4-B01
==========
Collated Motifs

    Allele: KE13-4-B01 | R1+R2 | [410/510] | %totalreads:80.39 | percentile:100 | %top5reads:93.18 | %readsfilteredfor1%:94.25 | %readsfilteredfor10%:100.0
    Motifs: total distinct sites |96|, total unique TFs |45| (motifs for 3 TFs occur >1x)
  Synopsis: relative to reference sequence--# lost sites |0|, # new sites |0|
   Details: lost ||
             new ||

     query GCCCTGAAGCTCCCGGGCTGCAGGGAACCATGAGGCTGCCTGGGCTTCCGGCTGCTGCAGGATAAAGGTCAACTC-TGCTGGAACACGAGGTTCTGCACCATCTGGGCTTGCCTGCCTCTGGCCTCATCTCCCCGGGCTCTCCAACCTGCACTCCATGTCCAGCCAGACCAAACA
           ||||||||||||||||||||||||||||||||||||||||||||||||||||||||||||||||||||||||||| |||||||||||||||||||||||||||||||||||||||||||||||||||||||||||||||||||||||||||||||||||||||||||||||||||
 reference GCCCTGAAGCTCCCGGGCTGCAGGGAACCATGAGGCTGCCTGGGCTTCCGGCTGCTGCAGGATAAAGGTCAACTCTTGCTGGAACACGAGGTTCTGCACCATCTGGGCTTGCCTGCCTCTGGCCTCATCTCCCCGGGCTCTCCAACCTGCACTCCATGTCCAGCCAGACCAAACA
```

### GBS -33.75 kb

lost GBS
new GBS

#### 14-2 B09 (clone #1) homozygous Δ

```
==========
KE14-2-B09
==========
   Allele: KE14-2-B09 | R1+R2 | [416/520] | %totalreads:80.0 | percentile:100 | %top5reads:96.74 | %readsfilteredfor1%:98.58 | %readsfilteredfor10%:100.0
   Motifs: total distinct sites |245|, total unique TFs |151| (motifs for 68 TFs occur >1x)
 Synopsis: relative to reference sequence--# lost sites |18|, # new sites |5|
  Details: lost |CENPB:1, EWSR1-FLI1:2, GCM1:1, GCM2:1, MTF1:1, NFYA:1, NKX2-8:1, NR3C1:2, NR3C2:2, Nkx2-5(var.2):2, RXRA::VDR:1, SNAI2:1, Vdr:1, ZNF263:1|
            new |Ar:1, NR3C2:1, Nkx2-5(var.2):1, PRDM1:1, YY2:1|

          NEW motifs:
          plus(+) strand:
                                                         GGGCTCTCCATGTCCAG |-- NR3C2 (MA0727.1) (pval 0.000884) [note, approx. position]
                                                         GGGCTCTCCAT |-- Nkx2-5(var.2) (MA0503.1) (pval 0.000226) [note, approx. position]
                                                           GCTCTCCATGT |-- YY2 (MA0748.1) (pval 0.000879) [note, approx. position]
          minus(-) strand:
                                                                     CCCGAGAGGTACAGG |-- PRDM1 (MA0508.1) (pval 0.000835) [note, approx. position]
                                                                     CCCGAGAGGTACAGGTC |-- Ar (MA0007.3) (pval 0.000862) [note, approx. position]

    query GGTCAACTCTTGCTGGAACACGAGGTTCTGCACCATCTGGGCTTGCCTGCCTCTGGCCTCATCTCCCCGGGCT------------CTCCATGTCCAGCCAGACCAAACAACTCCCAGTGCCCACCAGAGAACACCGGGCTGCTTCACGCCTGTGTGCTTTTATACACACTGTTCC
          |||||||||||||||||||||||||||||||||||||||||||||||||||||||||||||||||||||||||            ||||||||||||||||||||||||||||||||||||||||||||||||||||||||||||||||||||||||||||||||||||||||||
reference GGTCAACTCTTGCTGGAACACGAGGTTCTGCACCATCTGGGCTTGCCTGCCTCTGGCCTCATCTCCCCGGGCTCTCCAACCTGCACTCCATGTCCAGCCAGACCAAACAACTCCCAGTGCCCACCAGAGAACACCGGGCTGCTTCACGCCTGTGTGCTTTTATACACACTGTTCC

          LOST motifs:
          plus(+) strand:
                                                                       GCACTCCAT |-- NKX2-8 (MA0673.1) (pval 0.000514)
                                                             TCTCCAACCTGCAC |-- MTF1 (MA0863.1) (pval 0.000231)
                                                                     CTGCACTCCAT |-- Nkx2-5(var.2) (MA0503.1) (pval 0.000418)
                                                                     CTGCACTCCATGTCCAG |-- NR3C1 (MA0113.3) (pval 0.000902)
                                                         GGGCTCTCCAA |-- Nkx2-5(var.2) (MA0503.1) (pval 0.000128)
                                                                     CTGCACTCCATGTCCAG |-- NR3C2 (MA0727.1) (pval 0.000399)
                                                       CCCGGGCTCTCCAAC |-- CENPB (MA0637.1) (pval 0.000469)
          minus(-) strand:
                                                                TTGGACGTGA |-- GCM2 (MA0767.1) (pval 0.000892)
                                                          GAGAGGTTGGACGTGAGGTAC |-- ZNF263 (MA0528.1) (pval 0.00075)
                                                              GGTTGGACGTGAGGTACA |-- EWSR1-FLI1 (MA0149.1) (pval 0.000773)
                                                               GTTGGACGT |-- SNAI2 (MA0745.1) (pval 0.000268)
                                                          GAGAGGTTGGACGTGAGG |-- EWSR1-FLI1 (MA0149.1) (pval 0.000653)
                                                                      ACGTGAGGTACAGGT |-- RXRA::VDR (MA0074.1) (pval 0.000563)
                                                                     GACGTGAGGTACAGGTC |-- NR3C1 (MA0113.3) (pval 0.000894)
                                                          GAGAGGTTGGACGTGAGG |-- NFYA (MA0060.2) (pval 0.000631)
                                                                     GACGTGAGGTACAGGTC |-- NR3C2 (MA0727.1) (pval 0.000377)
                                                                      ACGTGAGGTACAGGTC |-- Vdr (MA0693.1) (pval 0.000979)
                                                               GTTGGACGTGA |-- GCM1 (MA0646.1) (pval 0.000969)
```

#### 14-4 E08 (clone #2) homozygous +1bp

```
==========
KE14-4-E08
==========
   Allele: KE14-4-E08 | R1+R2 | [220/316] | %totalreads:69.62 | percentile:100 | %top5reads:94.02 | %readsfilteredfor1%:97.35 | %readsfilteredfor10%:100.0
   Motifs: total distinct sites |263|, total unique TFs |160| (motifs for 74 TFs occur >1x)
 Synopsis: relative to reference sequence--# lost sites |13|, # new sites |18|
  Details: lost |EWSR1-FLI1:2, GCM1:1, GCM2:1, MTF1:1, NFYA:1, NR3C1:2, NR3C2:2, Nkx2-5(var.2):1, SNAI2:1, ZNF263:1|
            new |Esrra:1, Esrrg:1, HNF4G:1, Hnf4a:1, NR2C2:1, NR3C1:2, NR3C2:2, Nkx2-5(var.2):1, Nr2e3:1, Nr5a2:1, RXRB:1, RXRG:1, Rxra:2, ZNF143:1, ZNF263:1|

          NEW motifs:
          plus(+) strand:
                                                                    TTGCACTCCAT |-- Nkx2-5(var.2) (MA0503.1) (pval 0.000411)
                                                                    TTGCACTCCATGTCCAG |-- NR3C1 (MA0113.3) (pval 0.0008)
                                                            CTCCAACCTTGCACTC |-- ZNF143 (MA0088.2) (pval 0.000743)
                                                               CAACCTT |-- Nr2e3 (MA0164.1) (pval 0.000384)
                                                                    TTGCACTCCATGTCCAG |-- NR3C2 (MA0727.1) (pval 0.000365)
          minus(-) strand:
                                                              GGTTGGAACG |-- Esrrg (MA0643.1) (pval 0.000578)
                                                              GGTTGGAACGTGAGG |-- HNF4G (MA0484.1) (pval 0.000612)
                                                           AGAGGTTGGAACGTGAGGTAC |-- ZNF263 (MA0528.1) (pval 0.000413)
                                                             AGGTTGGAACGTGAG |-- Nr5a2 (MA0505.1) (pval 0.000197)
                                                              GGTTGGAACGT |-- Esrra (MA0592.2) (pval 0.000855)
                                                              GGTTGGAACGTGAGG |-- NR2C2 (MA0504.1) (pval 0.000121)
                                                              GGTTGGAACGTGAG |-- Rxra (MA0512.2) (pval 0.000378)
                                                                    AACGTGAGGTACAGGTC |-- NR3C1 (MA0113.3) (pval 0.000807)
                                                              GGTTGGAACGTGAG |-- RXRB (MA0855.1) (pval 0.000222)
                                                               GTTGGAACGTGAGG |-- Rxra (MA0512.2) (pval 0.000852)
                                                                    AACGTGAGGTACAGGTC |-- NR3C2 (MA0727.1) (pval 0.000402)
                                                              GGTTGGAACGTGAG |-- RXRG (MA0856.1) (pval 0.000155)
                                                            GAGGTTGGAACGTGAG |-- Hnf4a (MA0114.3) (pval 0.000263)

    query GGTCAACTCTTGCTGGAACACGAGGTTCTGCACCATCTGGGCTTGCCTGCCTCTGGCCTCATCTCCCCGGGCTCTCCAACCTTGCACTCCATGTCCAGCCAGACCAAACAACTCCCAGTGCCCACCAGAGAACACCGGGCTGCTTCACGCCTGTGTGCTTTTATACACACTGTTCC
          ||||||||||||||||||||||||||||||||||||||||||||||||||||||||||||||||||||||||||||||||| ||||||||||||||||||||||||||||||||||||||||||||||||||||||||||||||||||||||||||||||||||||||||||||||
reference GGTCAACTCTTGCTGGAACACGAGGTTCTGCACCATCTGGGCTTGCCTGCCTCTGGCCTCATCTCCCCGGGCTCTCCAACC-TGCACTCCATGTCCAGCCAGACCAAACAACTCCCAGTGCCCACCAGAGAACACCGGGCTGCTTCACGCCTGTGTGCTTTTATACACACTGTTCC

          LOST motifs:
          plus(+) strand:
                                                                         TCTCCAACCTGCAC |-- MTF1 (MA0863.1) (pval 0.000231) [note, approx. position]
                                                                      CTGCACTCCAT |-- Nkx2-5(var.2) (MA0503.1) (pval 0.000418) [note, approx. position]
                                                                  CTGCACTCCATGTCCAG |-- NR3C1 (MA0113.3) (pval 0.000902) [note, approx. position]
                                                                  CTGCACTCCATGTCCAG |-- NR3C2 (MA0727.1) (pval 0.000399) [note, approx. position]
          minus(-) strand:
                                                                 TTGGACGTGA |-- GCM2 (MA0767.1) (pval 0.000892) [note, approx. position]
                                                           GAGAGGTTGGACGTGAGGTAC |-- ZNF263 (MA0528.1) (pval 0.00075) [note, approx. position]
                                                               GGTTGGACGTGAGGTACA |-- EWSR1-FLI1 (MA0149.1) (pval 0.000773) [note, approx. position]
                                                                GTTGGACGT |-- SNAI2 (MA0745.1) (pval 0.000268) [note, approx. position]
                                                           GAGAGGTTGGACGTGAGG |-- EWSR1-FLI1 (MA0149.1) (pval 0.000653) [note, approx. position]
                                                                    GACGTGAGGTACAGGTC |-- NR3C1 (MA0113.3) (pval 0.000894) [note, approx. position]
                                                           GAGAGGTTGGACGTGAGG |-- NFYA (MA0060.2) (pval 0.000631) [note, approx. position]
                                                                    GACGTGAGGTACAGGTC |-- NR3C2 (MA0727.1) (pval 0.000377) [note, approx. position]
                                                                GTTGGACGTGA |-- GCM1 (MA0646.1) (pval 0.000969) [note, approx. position]
```

#### 14-5 B05 (clone #3) bialleleic sub/+1bp

```
==========
KE14-5-B05
==========
   Allele: KE14-5-B05 | R1+R2 | [121/323] | %totalreads:37.46 | percentile:100 | %top5reads:48.99 | %readsfilteredfor1%:50.42 | %readsfilteredfor10%:50.42
   Motifs: total distinct sites |261|, total unique TFs |158| (motifs for 74 TFs occur >1x)
 Synopsis: relative to reference sequence--# lost sites |15|, # new sites |18|
  Details: lost |EWSR1-FLI1:2, GCM1:1, GCM2:1, MTF1:1, NFYA:1, NR3C1:2, NR3C2:2, Nkx2-5(var.2):1, RXRA::VDR:1, SNAI2:1, Vdr:1, ZNF263:1|
            new |CENPB:1, EGR4:1, GCM1:1, GCM2:1, MTF1:1, NFYA:1, NR3C1:1, NR3C2:2, Nkx2-5(var.2):1, RREB1:1, RUNX2:1, RUNX3:1, RXRA::VDR:1, SP4:1, SP8:1, ZNF263:1, ZNF740:1|

          NEW motifs:
          plus(+) strand:
                                                              CCGCACTCCATGTCCAG |-- NR3C2 (MA0727.1) (pval 0.000464)
          minus(-) strand:
                                                              GGCGTGAGGTACAGGTC |-- NR3C2 (MA0727.1) (pval 0.000332)
                                                              GGCGTGAGGTACAGGTC |-- NR3C1 (MA0113.3) (pval 0.000853)

    query GGTCAACTCTTGCTGGAACACGAGGTTCTGCACCATCTGGGCTTGCCTGCCTCTGGCCTCATCTCCCCGGGCTCTCCAACCCGCACTCCATGTCCAGCCAGACCAAACAACTCCCAGTGCCCACCAGAGAACACCGGGCTGCTTCACGCCTGTGTGCTTTTATACACACTGTTCC
          ||||||||||||||||||||||||||||||||||||||||||||||||||||||||||||||||||||||||||||||||| |||||||||||||||||||||||||||||||||||||||||||||||||||||||||||||||||||||||||||||||||||||||||||||||
reference GGTCAACTCTTGCTGGAACACGAGGTTCTGCACCATCTGGGCTTGCCTGCCTCTGGCCTCATCTCCCCGGGCTCTCCAACCTGCACTCCATGTCCAGCCAGACCAAACAACTCCCAGTGCCCACCAGAGAACACCGGGCTGCTTCACGCCTGTGTGCTTTTATACACACTGTTCC

          LOST motifs:
          plus(+) strand:
                                                              CTGCACTCCATGTCCAG |-- NR3C1 (MA0113.3) (pval 0.000902)
                                                              CTGCACTCCATGTCCAG |-- NR3C2 (MA0727.1) (pval 0.000399)
          minus(-) strand:
                                                              GACGTGAGGTACAGGTC |-- NR3C1 (MA0113.3) (pval 0.000894)

   Allele: KE14-5-B05 | R1+R2 | [119/323] | %totalreads:36.84 | percentile:98 | %top5reads:48.18 | %readsfilteredfor1%:49.58 | %readsfilteredfor10%:49.58
   Motifs: total distinct sites |263|, total unique TFs |160| (motifs for 74 TFs occur >1x)
 Synopsis: relative to reference sequence--# lost sites |13|, # new sites |18|
  Details: lost |EWSR1-FLI1:2, GCM1:1, GCM2:1, MTF1:1, NFYA:1, NR3C1:2, NR3C2:2, Nkx2-5(var.2):1, SNAI2:1, ZNF263:1|
            new |Esrra:1, Esrrg:1, HNF4G:1, Hnf4a:1, NR2C2:1, NR3C1:2, NR3C2:2, Nkx2-5(var.2):1, Nr2e3:1, Nr5a2:1, RXRB:1, RXRG:1, Rxra:2, ZNF143:1, ZNF263:1|

          NEW motifs:
          plus(+) strand:
                                                              TTGCACTCCATGTCCAG |-- NR3C1 (MA0113.3) (pval 0.0008)
                                                              TTGCACTCCATGTCCAG |-- NR3C2 (MA0727.1) (pval 0.000365)
          minus(-) strand:
                                                              AACGTGAGGTACAGGTC |-- NR3C1 (MA0113.3) (pval 0.000807)
                                                              AACGTGAGGTACAGGTC |-- NR3C2 (MA0727.1) (pval 0.000402)

    query GGTCAACTCTTGCTGGAACACGAGGTTCTGCACCATCTGGGCTTGCCTGCCTCTGGCCTCATCTCCCCGGGCTCTCCAACCTTGCACTCCATGTCCAGCCAGACCAAACAACTCCCAGTGCCCACCAGAGAACACCGGGCTGCTTCACGCCTGTGTGCTTTTATACACACTGTTCC
          ||||||||||||||||||||||||||||||||||||||||||||||||||||||||||||||||||||||||||||||||| ||||||||||||||||||||||||||||||||||||||||||||||||||||||||||||||||||||||||||||||||||||||||||||||
reference GGTCAACTCTTGCTGGAACACGAGGTTCTGCACCATCTGGGCTTGCCTGCCTCTGGCCTCATCTCCCCGGGCTCTCCAACC-TGCACTCCATGTCCAGCCAGACCAAACAACTCCCAGTGCCCACCAGAGAACACCGGGCTGCTTCACGCCTGTGTGCTTTTATACACACTGTTCC

          LOST motifs:
          plus(+) strand:
                                                             CTGCACTCCATGTCCAG |-- NR3C1 (MA0113.3) (pval 0.000902) [note, approx. position]
                                                             CTGCACTCCATGTCCAG |-- NR3C2 (MA0727.1) (pval 0.000399) [note, approx. position]
          minus(-) strand:
                                                              GACGTGAGGTACAGGTC |-- NR3C1 (MA0113.3) (pval 0.000894) [note, approx. position]
                                                              GACGTGAGGTACAGGTC |-- NR3C2 (MA0727.1) (pval 0.000377) [note, approx. position]
```

*17-G05 biallelic Δ*

#### GBS -33.99 kb

*lost GBS*

```
==========
KE17-5-G05
==========
   Allele: KE17-5-G05 | R1+R2 | [138/466] | %totalreads:29.61 | percentile:100 | %top5reads:52.47 | %readsfilteredfor1%:53.49 | %readsfilteredfor10%:54.55
   Motifs: total distinct sites |142|, total unique TFs |106| (motifs for 28 TFs occur >1x)
 Synopsis: relative to reference sequence--# lost sites |17|, # new sites |13|
  Details: lost |EGR3:1, ETV6:1, HIC2:1, KLF13:1, KLF16:1, KLF5:1, Klf12:1, MZF1:1, NR3C1:2, NR3C2:2, RREB1:1, SP3:1, SP4:1, STAT3:1, ZIC4:1|
            new |BATF3:1, Dux:1, EGR2:1, EGR3:1, EGR4:1, GLIS2:1, KLF13:1, Klf12:1, NFYB:1, Nr1h3::Rxra:1, SP4:1, ZBED1:1, ZIC4:1|

           NEW motifs:
           plus(+) strand:
                                                                                           ACACCCAGTCA |-- EGR2 (MA0472.2) (pval 0.000891) [note, approx. position]
                                                                                         CCACACCCAGTCAAGA |-- EGR4 (MA0733.1) (pval 0.000587) [note, approx. position]
                                                                                       GGCCACACCCAGTCA |-- Klf12 (MA0742.1) (pval 1.41e-06) [note, approx. position]
                                                                                     CTGGCCACACCCAGTCA |-- SP4 (MA0685.1) (pval 1.8e-05) [note, approx. position]
                                                                                          CACACCCAGTCAAG |-- GLIS2 (MA0736.1) (pval 0.000699) [note, approx. position]
                                                                                         CCACACCCAGTCAAG |-- EGR3 (MA0732.1) (pval 0.000191) [note, approx. position]
                                                                                                CAGTCAAGACATT |-- ZBED1 (MA0749.1) (pval 0.000373) [note, approx. position]
                                                                                        GCCACACCCAGTCAA |-- NFYB (MA0502.1) (pval 0.00074) [note, approx. position]
                                                                                      TGGCCACACCCAGTCAAG |-- KLF13 (MA0657.1) (pval 0.000107) [note, approx. position]
                                                                                               CCAGTCAA |-- Dux (MA0611.1) (pval 0.00096) [note, approx. position]
                                                                                                   TCAAGACATTATAA |-- BATF3 (MA0835.1) (pval 0.000676) [note, approx. position]
                                                                                       GGCCACACCCAGTCA |-- ZIC4 (MA0751.1) (pval 0.000944) [note, approx. position]
           minus(-) strand:
                                                                                              GAGACCGGTGTGGGTCAGT |-- Nr1h3::Rxra (MA0494.1) (pval 0.000617) [note, approx. position]

    query  CACACTGTTCCTTCTACCTGGAACGTCCTGCACCCTTGCTCTGCCTGGCCAATTCCTCTCTCCCTGAACCTTCTGCTCTGGCCACACCCAGT-----------CAAGACATTATAATGATCCGTTTTTGTGCATTTTCCCAAGTAAACCAGGGACTCAGACTTTCCAAGTAGCCC
           ||||||||||||||||||||||||||||||||||||||||||||||||||||||||||||||||||||||||||||||||||||||||||||           ||||||||||||||||||||||||||||||||||||||||||||||||||||||||||||||||||||||||
reference  CACACTGTTCCTTCTACCTGGAACGTCCTGCACCCTTGCTCTGCCTGGCCAATTCCTCTCTCCCTGAACCTTCTGCTCTGGCCACACCCAGTCCCCAACTTCCCAAGACATTATAATGATCCGTTTTTGTGCATTTTCCCAAGTAAACCAGGGACTCAGACTTTCCAAGTAGCCC

           LOST motifs:
           plus(+) strand:
                                                                                              CCCAGTCCCC |-- KLF5 (MA0599.1) (pval 0.000935)
                                                                                       GGCCACACCCAGTCC |-- Klf12 (MA0742.1) (pval 1.32e-06)
                                                                                            CACCCAGTCCCCAACTTCCC |-- RREB1 (MA0073.1) (pval 0.000126)
                                                                                      TGGCCACACCCAGTCCCC |-- KLF13 (MA0657.1) (pval 0.000233)
                                                                                       GGCCACACCCAGTCC |-- ZIC4 (MA0751.1) (pval 0.00073)
                                                                                              CCCAGTCCCCA |-- SP3 (MA0746.1) (pval 0.000633)
                                                                                       GGCCACACCCAGTCCCC |-- NR3C1 (MA0113.3) (pval 0.000686)
                                                                                                  GTCCCCAAC |-- HIC2 (MA0738.1) (pval 0.000737)
                                                                                              CCCAGTCCCCA |-- KLF16 (MA0741.1) (pval 0.000895)
                                                                                         CCACACCCAGTCCCC |-- EGR3 (MA0732.1) (pval 0.000354)
                                                                                       GGCCACACCCAGTCCCC |-- NR3C2 (MA0727.1) (pval 0.000427)
                                                                                     CTGGCCACACCCAGTCC |-- SP4 (MA0685.1) (pval 1.4e-05)
           minus(-) strand:
                                                                                                        GAAGGGTTCTG |-- STAT3 (MA0144.2) (pval 0.00064)
                                                                                                      TTGAAGGGTT |-- ETV6 (MA0645.1) (pval 0.000944)
                                                                                       CCGGTGTGGGTCAGGGG |-- NR3C1 (MA0113.3) (pval 0.000565)
                                                                                                   AGGGGT |-- MZF1 (MA0056.1) (pval 0.000154)
                                                                                       CCGGTGTGGGTCAGGGG |-- NR3C2 (MA0727.1) (pval 0.000343)

   Allele: KE17-5-G05 | R1+R2 | [115/466] | %totalreads:24.68 | percentile:99 | %top5reads:43.73 | %readsfilteredfor1%:44.57 | %readsfilteredfor10%:45.45
   Motifs: total distinct sites |137|, total unique TFs |101| (motifs for 26 TFs occur >1x)
 Synopsis: relative to reference sequence--# lost sites |17|, # new sites |8|
  Details: lost |EGR3:1, HIC2:1, KLF13:1, KLF14:1, KLF16:1, KLF5:1, Klf12:1, MZF1:1, NR3C1:2, NR3C2:2, RREB1:1, SP3:1, SP4:1, SP8:1, ZIC4:1|
            new |INSM1:1, KLF13:1, KLF14:1, Klf12:1, RREB1:1, Rarb(var.2):1, SP4:1, SP8:1|

           NEW motifs:
           plus(+) strand:
                                                                                        GCCACACCCAGA |-- SP8 (MA0747.1) (pval 2.59e-05) [note, approx. position]
                                                                                             ACCCAGAACTTCCCAAGACA |-- RREB1 (MA0073.1) (pval 0.00039) [note, approx. position]
                                                                                       GGCCACACCCAGAA |-- KLF14 (MA0740.1) (pval 9.69e-05) [note, approx. position]
                                                                                     CTGGCCACACCCAGAAC |-- SP4 (MA0685.1) (pval 3.4e-05) [note, approx. position]
                                                                                      TGGCCACACCCAGAACTT |-- KLF13 (MA0657.1) (pval 0.00027) [note, approx. position]
                                                                                       GGCCACACCCAGAAC |-- Klf12 (MA0742.1) (pval 1.37e-06) [note, approx. position]
           minus(-) strand:
                                                                                           ACCGGTGTGGGTCTTGA |-- Rarb(var.2) (MA0858.1) (pval 0.000963) [note, approx. position]
                                                                                               GTGTGGGTCTTG |-- INSM1 (MA0155.1) (pval 0.000531) [note, approx. position]

    query  CACACTGTTCCTTCTACCTGGAACGTCCTGCACCCTTGCTCTGCCTGGCCAATTCCTCTCTCCCTGAACCTTCTGCTCTGGCCACACCCAG-----AACTTCCCAAGACATTATAATGATCCGTTTTTGTGCATTTTCCCAAGTAAACCAGGGACTCAGACTTTCCAAGTAGCCC
           |||||||||||||||||||||||||||||||||||||||||||||||||||||||||||||||||||||||||||||||||||||||||||     |||||||||||||||||||||||||||||||||||||||||||||||||||||||||||||||||||||||||||||||
reference  CACACTGTTCCTTCTACCTGGAACGTCCTGCACCCTTGCTCTGCCTGGCCAATTCCTCTCTCCCTGAACCTTCTGCTCTGGCCACACCCAGTCCCCAACTTCCCAAGACATTATAATGATCCGTTTTTGTGCATTTTCCCAAGTAAACCAGGGACTCAGACTTTCCAAGTAGCCC

           LOST motifs:
           plus(+) strand:
                                                                                              CCCAGTCCCC |-- KLF5 (MA0599.1) (pval 0.000935)
                                                                                       GGCCACACCCAGTCC |-- Klf12 (MA0742.1) (pval 1.32e-06)
                                                                                            CACCCAGTCCCCAACTTCCC |-- RREB1 (MA0073.1) (pval 0.000126)
                                                                                        GCCACACCCAGT |-- SP8 (MA0747.1) (pval 1.05e-05)
                                                                                      TGGCCACACCCAGTCCCC |-- KLF13 (MA0657.1) (pval 0.000233)
                                                                                       GGCCACACCCAGTCC |-- ZIC4 (MA0751.1) (pval 0.00073)
                                                                                              CCCAGTCCCCA |-- SP3 (MA0746.1) (pval 0.000633)
                                                                                       GGCCACACCCAGTCCCC |-- NR3C1 (MA0113.3) (pval 0.000686)
                                                                                       GGCCACACCCAGTC |-- KLF14 (MA0740.1) (pval 3.74e-05)
                                                                                                  GTCCCCAAC |-- HIC2 (MA0738.1) (pval 0.000737)
                                                                                              CCCAGTCCCCA |-- KLF16 (MA0741.1) (pval 0.000895)
                                                                                         CCACACCCAGTCCCC |-- EGR3 (MA0732.1) (pval 0.000354)
                                                                                       GGCCACACCCAGTCCCC |-- NR3C2 (MA0727.1) (pval 0.000427)
                                                                                     CTGGCCACACCCAGTCC |-- SP4 (MA0685.1) (pval 1.4e-05)
           minus(-) strand:
                                                                                       CCGGTGTGGGTCAGGGG |-- NR3C1 (MA0113.3) (pval 0.000565)
                                                                                                   AGGGGT |-- MZF1 (MA0056.1) (pval 0.000154)
                                                                                       CCGGTGTGGGTCAGGGG |-- NR3C2 (MA0727.1) (pval 0.000343)
```

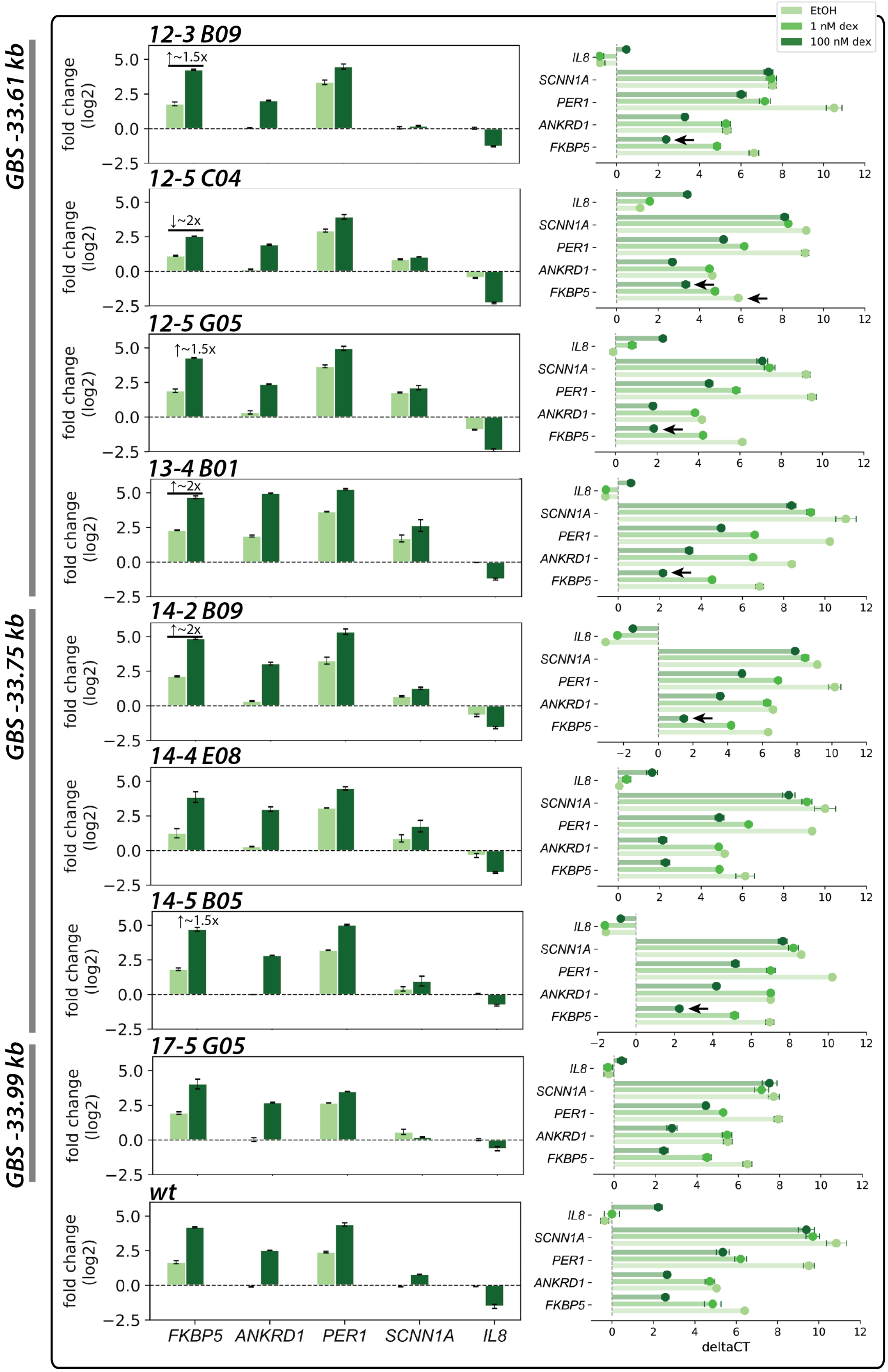

GBS -33.61 kb
GBS -33.75 kb
GBS -33.99 kb
EtOH
1 nM dex
100 nM dex
12-3 B09
12-5 C04
12-5 G05
13-4 B01
14-2 B09
14-4 E08
14-5 B05
17-5 G05
wt
↑~1.5x
↓~2x
↑~2x
fold change (log2)
FKBP5
ANKRD1
PER1
SCNN1A
IL8
deltaCT

**S14. Luciferase reporter activity for a subset of sampled *FKBP5* GORs, cloned as 500-bp fragments flanking a ChIP-seq peak summit in pGL4.10 with minimal promoter (E4TATA).** GBS coordinates are indicated as positions relative to *FKBP5* transcription start site (+86.85 kb (GOR2) intronic, others upstream).

- **GOR2** (peak summit GBS = +86.85 kb) is dex-responsive in a luciferase reporter assay, however, biallelic deletions of the native GBS do not impact dex induction of *FKBP5* from the chromosomal locus (**Supp. Fig. 13a**, with caveat that one allele in each of the example clones reconstituted a new GBS match, and effect on GR occupancy was not experimentally tested).
- **GOR5** (multiple GBS matches at peak summit, including -33.61, -33.75, -33.99 kb) is also dex-responsive, and similarly, biallelic deletions of motif matches in this region do not ablate dex induction of *FKBP5* from the chromosomal locus (**Supp. Fig. 13b**).
- **GOR6**, centered at two GR ChIP-seq peaks, is not dex-responsive in the luciferase reporter assay, and as for other GOR summits tested individually by indel editing, its deletion does not ablate dex induction of *FKBP5* from the chromosomal locus in A549 (data not shown).
- **GOR8** (peak summit GBS = -43.08 kb) is weakly dex-responsive, and as for other GOR summits tested individually by indel editing, its deletion does not ablate dex induction of *FKBP5* from the chromosomal locus in A549 (data not shown). *Note*, not all GORs indicated in Fig. 5 were tested in the luciferase reporter assay.

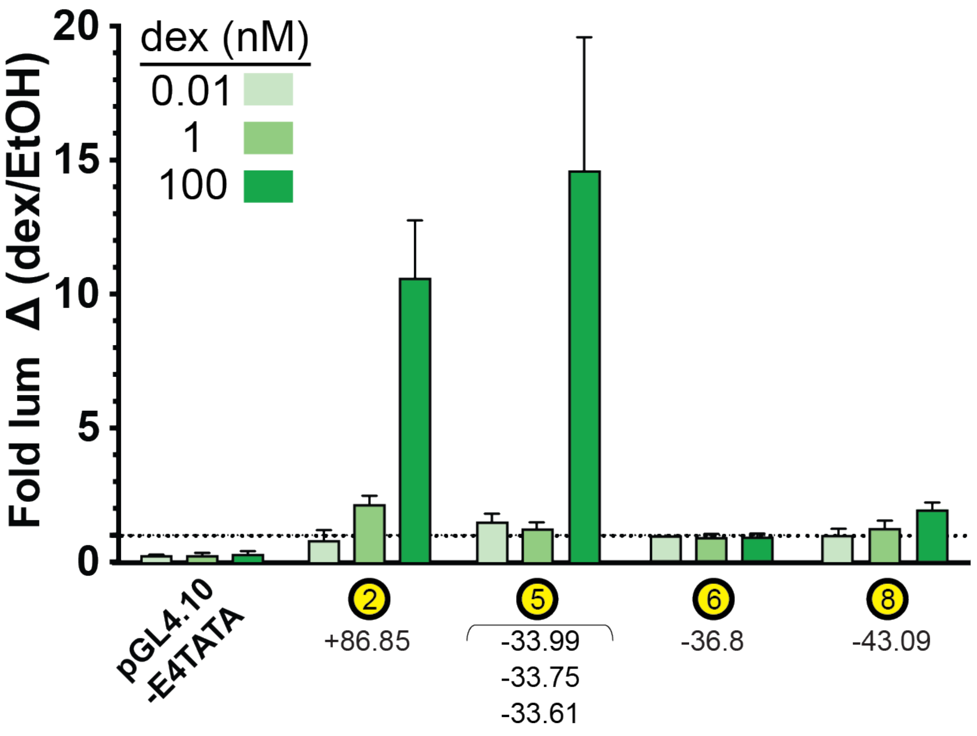

**S15. Amplicon length considerations in library preparation: PCR product purification and sequencing recovery.**

Some editing cases may present situations where amplicons of varied length are pooled as a common library for sequencing; we empirically tested amplicon length-dependent recovery of reads to assess length-dependent recovery of short read sequences on an Illumina® platform.

***(a) Purification of PCR products.*** In library preparation, SPRI clean-up retains longer amplicons more effectively than column clean-up. Four independent PCR products (labeled *red* 1-4, ranging from ~358-801 bp) were pooled, purified either by SPRISelect (Beckman Coulter, Brea, CA) or DNA Clean & Concentrator-5 (Zymo Research, Irvine, CA), and analyzed by Bioanalyzer 2100 (High Sensitivity DNA Assay, Agilent). Shown are traces (*left*), gel-like image and expected sizes of wild type PCR amplicons without adaptors (*right*).

***(b) Amplicon length-dependent sequence recovery on MiSeq.*** To test the amplicon lengths most readily compatible with Illumina® MiSeq technology, we prepared a library of seven amplicons ranging in lengths from 100 – 1500 bp, identical in sequence at their ends outside of internal extensions for longer amplicons. Amplicons were uniquely barcoded such that they could be pooled in separate library sets, defined by molar ratios specified as 1:1:1:1:1:1:1 or 1:10:50:100:200:500:1000 (relative to 100:175:300:500:800:1000:1500 bp amplicons) (*see* **Supp. Methods**).

*(i)* among total mapped reads, amplicon size >500 bp was associated with representation below expected (expected ~14.3% for each amplicon in 1:1 ratio [*horizontal dashed line*, upper left plot], *vertical black lines*, lower left plot); *insets*: absolute mapped read counts relative to amplicon size (bp);

*(ii)* among top 15 most abundant reads identified for each amplicon in each library ratio, the % of R1 & R2 reads that mapped to each amplicon reference sequence appeared relatively consistent for amplicons up to 800 bp, and was markedly blunted for an amplicon 1500 bp in length. This suggested to us that amplicons up to 800 bp generally perform well for clustering and sequencing, but that performance drops for amplicons beyond these lengths (pronounced for 1500 bp).

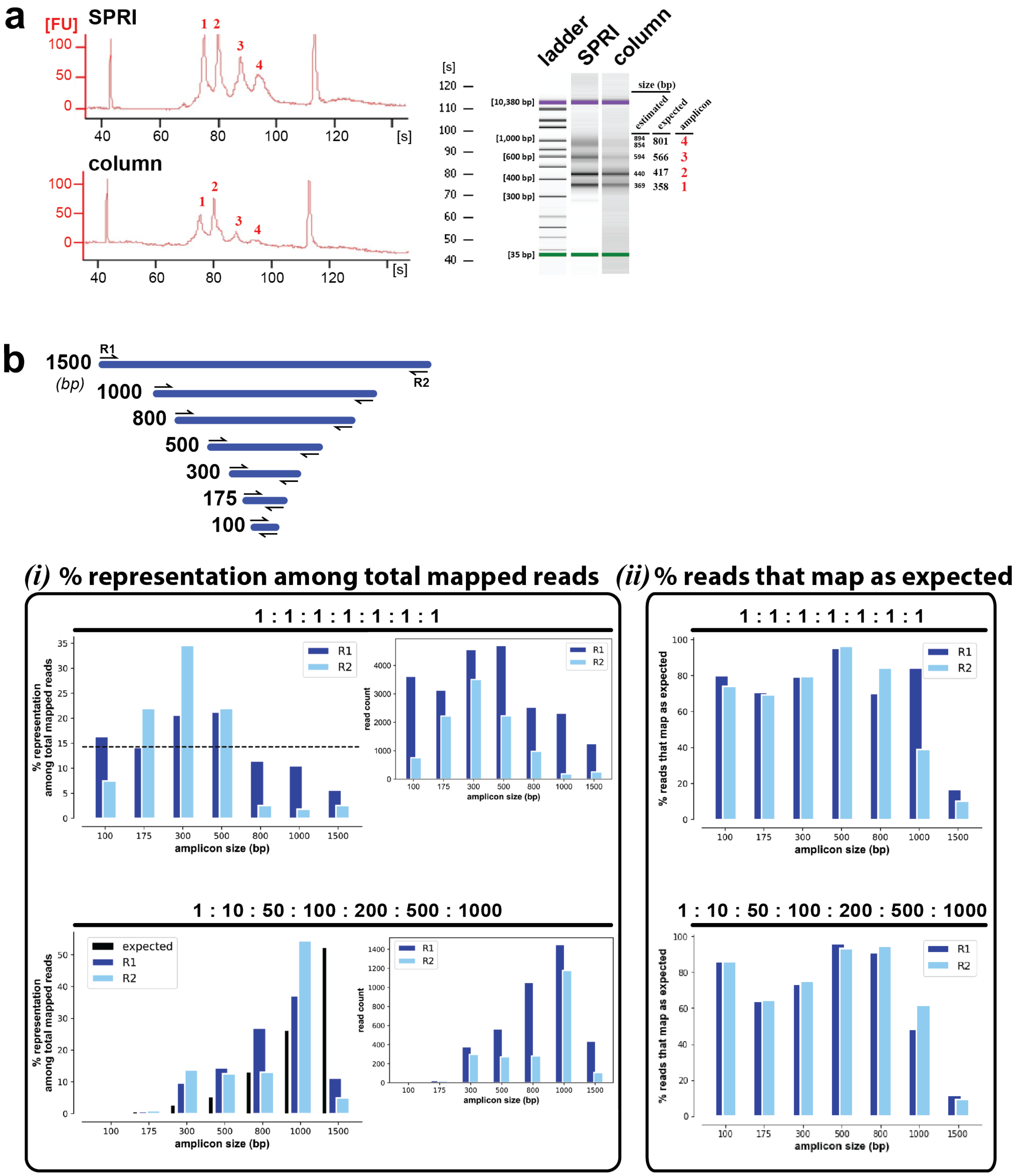
a
[FU]
SPRI
column
[s]
ladder
SPRI
column
size (bp)
estimated
expected
amplicon
[10,380 bp]
[1,000 bp]
[600 bp]
[400 bp]
[300 bp]
[35 bp]
894
854
801
4
594
566
3
440
417
2
369
358
1
b
1500
(bp)
1000
800
500
300
175
100
R1
R2
(i) % representation among total mapped reads
(ii) % reads that map as expected
1 : 1 : 1 : 1 : 1 : 1 : 1
1 : 10 : 50 : 100 : 200 : 500 : 1000
% representation among total mapped reads
% reads that map as expected
read count
amplicon size (bp)
expected
R1
R2

#### Supplemental Methods

***sgRNA cloning into pSpCas9(BB)-2A-Puro vectors***—20-nt single guide (sg) RNA sequences[1] were designed using online tools: CRISPR-MIT (crispr.mit.edu) and sgRNA Designer[2] (https://portals.broadinstitute.org/gpp/public/analysis-tools/sgrna-design). Guides were selected with predicted incision position within or near to targeted GBS motifs underlying GOR peak summits, and prioritized based on predicted on-target/off-target (CRISPR-MIT) and efficiency scores (sgRNA Designer). Oligo sequences for sgRNA duplexes with short single-stranded DNA overhangs (compatible with *Bbs*I cloning) were designed using the shell script *'sgRNA_BbsI_oligo_conversions.sh'* (https://github.com/YamamotoLabUCSF/sgRNA_BbsI_oligo_conversions.sh), which appends 'CACCG' to the 5' end of the sgRNA DNA sequence ('fwd'), and generates a reverse complement with 'AAAC' appended at its 5' and 'C' appended at its 3' end ('rev'). Oligos (unphosphorylated) were ordered from Integrated DNA Technologies (IDT, Coralville, IA). Oligos ('gRNA top' and 'gRNA bottom') were phosphorylated and annealed by mixing 1 µL 'top' (100 µM) and 1 µL 'bottom' (100 µM) oligos with 1 µL T4 PNK (New England Biolabs) and 1 µL T4 DNA ligase buffer in a 10 µL reaction volume, with incubation in a Peltier PTC-200 DNA Engine Thermal Cycler using program GRNA*ANL (37 °C for 30 min, 95 °C for 5 min, ramp to 25 °C at 0.1 °C/sec, 10 °C indefinitely). Phosphorylated, annealed sgRNA duplexes were diluted 1:20 with nuclease-free $H_2O$ (Ambion).

1 µL 1:20 sgRNA duplex was mixed with 1 µL pSpCas9(BB)-2A-Puro (PX459) V2.0 (100 ng/µL, gift from Feng Zhang; Addgene plasmid #62988, http://n2t.net/addgene:62988, RRIS:Addgene_62988), 1 µL CutSmart buffer, 1 µL DTT (10 mM), 0.4 µL ATP (25 mM), 0.5 µL *Bbs*I, 0.5 µL T4 DNA ligase + nuclease-free $H_2O$ to 20 µL. Vectors were assembled in a thermocycler using program CAS9ASMB ([37 °C for 5 min, 21 °C for 5 min], cycle 5x, 10 °C indefinitely). 1 µL ligation products were transformed into 8 µL competent *E. coli*, incubated on ice 30 for min., heat shocked for 30 sec, 42 °C, followed by incubation on ice for 5 min. 950 µL SOC was added, followed by incubation at 37 °C with shaking for 60 min.; finally, 100 µL was spread onto LB+Amp plates for overnight growth and selection. Candidates were Sanger sequenced (QuintaraBio, South San Francisco, CA) with oKE198 (U6_fwd, 5'-GAGGGCCTATTTCCCATGATTCC-3') to identify and confirm sgRNA cloned into *Bbs*I sites.

***A549 transfection and puromycin selection***—A549 cells (authenticated by short tandem repeat (STR) profiling (ATCC, Manassas, VA)) were plated in wells of a 24-well tissue culture plate (Corning Inc., Corning, NY) at $3x10^4$ cells/well in 500 µL DMEM/low glucose (HyClone Laboratories, Logan UT) containing 5% FBS (GemCell, Gemini Bio-Products, West Sacramento, CA), and maintained in an IncuSafe $CO_2$ incubator (Sanyo Scientific) at 37 °C. 24 h later, plasmids diluted to 100 ng/µL were transfected into plated cells under the following conditions: 500 ng plasmid was mixed in Opti-MEM Reduced Serum Media (ThermoFisher Scientific, Waltham, MA) in a final volume of 25 µL, then mixed with a 25 µL Opti-MEM volume containing 1 µL Lipofectamine 3000 (ThermoFisher); after 5 min incubation, the 50 µL transfection complex volume was delivered to plated cells (in total, twenty-five distinct Cas9 treatment combinations). After 24 hours, puromycin selection was imposed at 0.61 µg/mL (2-day SF90 determined for A549) by replacing media with puromycin (InvivoGen, San Diego, CA) @ SF90. Four days later, single cells were sorted into wells of 96-well plates containing 100 µL HAM'S F-12 (Lonza, Basel, Switzerland)/10% FBS using a BD FACSAria2 (Center for Advanced Technology, UCSF), and cultured for >2 weeks with media changes until ready for genotyping. For each of twenty-five

Cas9 treatment combinations, cells were sorted into 3-4 96-well plates (therefore sampling up to 288-384 clones (~1-1.3%) of each treatment population for genotypic evaluation), amounting to a total of ninety-six 96-well plates across Cas9 treatments. Following sequencing and genotypic analysis (*below*), selected clones were scaled up to 10-cm plates and aliquoted to 3 cryogenic vials for long-term storage in liquid $N_2$.

***Assessment of amplicon length-dependent sequence recovery on MiSeq (Supp. Fig. 15b)***—Seven amplicons ranging in size from 100-1500 bp (100, 175, 300, 500, 800, 1000, 1500) were PCR-amplified from pGL4.10 (Promega, GenBank AY738222); 100-1000 bp constructs were produced as stitched amplicons in two consecutive PCRs (A & B), such that all amplicons shared the same end sequences up to the maximum possible for each length: two fragments with engineered overlap were separately amplified in a PCR-1A, then stitched together as a single amplicon in a second PCR-1B. Full-length amplicons were isolated by agarose gel extraction (Qiagen QIAQuick Gel Extraction Kit), and re-amplified in a PCR-2 with primers to add extensions compatible with i5 and i7 barcode primers. Amplicon sets were uniquely barcoded in a PCR-3 as follows: amplicon set 'A1': all 7 amplicons barcoded with i5 A01; unique sizes independently barcoded with i7 A01-A07 (1500 bp-100 bp), amplicon set 'A3': all 7 amplicons barcoded with i5 A03; unique sizes independently barcoded with i7 A01-A07 (1500 bp-100 bp).

Amplicons were purified by DNA Clean & Concentrator-5 (Zymo Research, Irvine, CA), diluted to 5 pg/µL (approximated from NanoDrop spectrophotometry (ThermoFisher)), and each indexed amplicon was independently quantified by KAPA Library Quantification Kit for Illumina® Platforms (KK4844, KAPA Biosystems, Wilmington, MA) in duplicate, with six standards in triplicate. Amplicons were pooled in defined molar (amplicon) ratios as libraries (100 bp : 175 bp : 300 bp : 500 bp : 800 bp : 1000 bp : 1500 bp: 'A1'—1:1:1:1:1:1:1, 'A3'—1:10:50:100:200:500:1000). Libraries were diluted to 4 nM and prepared for sequencing using the MiSeq Nano Reagent Kit v2: PE, 2x150 bp (Illumina®, San Diego, CA), according to manufacturer instructions (final library concentration 7 pM, 10% each 'A1' and 'A3' library, 40% PhiX DNA spike-in (PhiX Control v3, Illumina®)). MiSeq cluster density was 1237 ± 1 k/mm$^2$, with 76.64% of reads passing filter @ %Q30=85.15 to yield 1,379,866 reads.

For 'A1' and 'A3' libraries, the top 15 read types for each amplicon were identified and quantified by bash, converted to fasta format (Python) and manually evaluated for alignment against amplicon sequences using the 'Align Multiple Sequences' function in SnapGene (GSL BioTech LLC, Chicago, IL). *% representation among total mapped reads* and *% reads that map as expected* were plotted in Python matplotlib as a function of amplicon size (bp) (**Supp. Fig. 15b**).

***MiSeq primer design and $T_m$ determination***—To genotype cells in 96-well plates, media was removed from wells by aspiration, cells were washed with 100 µL DPBS/-calcium,-magnesium (HyClone), and cells were trypsinized in 30 µL trypsin-EDTA (Gibco); 15 µL trypsinized volume was lysed with an equal volume of 2X Lysis Buffer (2X: 100 mM KCl, 20 mM Tris-HCl, pH 8.3, 5 mM $MgCl_2$, 0.9% NP-40, 0.9% Tween-20 + Proteinase K (Roche, recombinant PCR-grade, 19 mg/mL) @1:100) in a thermocycler programmed at 65 °C for 30 min, 95 °C for 15 min. Primers to amplify Cas9-targeted loci were designed in SnapGene to span 175 bp flanking a targeted GBS with predicted $T_m$ > 55 °C; 5' extensions to make PCR1 amplicons compatible with PCR2 amplification were added as described in **Supp. Fig. 5** (sequences available in **Supp. Excel file**). Primers were evaluated for empirical $T_a$ using the thermocycler gradient program KEPRIMERCHK (95 °C for 10 min., [95 °C for 10 sec, 55→65 °C for 20 sec over 8 temperature increments (55, 55.8, 57.1, 59, 61.2, 63.1, 64.4, 65 °C), 72 °C for 15 sec], cycle 35x, 72 °C for 5 min., 12 °C indefinitely).

***PCR1 & PCR2 for dual-indexed (i5 & i7) amplicon library preparation***—4 μL lysed sample was used as template in 20 μL PCR1 volumes (1X HF Phusion buffer, 200 μM dNTP (each), 225 nM each primer, ~1 U Phusion pol). PCR1 was performed using thermocycler program KEGENTYPE (98 °C for 2.5 min, [98 °C for 30 sec, Ta for 20 sec, 72 °C for 30 sec], cycle 30x total, 72 °C for 5 min, 12 °C indefinitely). 0.5 μL PCR1 was used as template in 20 μL PCR2 volumes (1X HF Phusion buffer, 200 μM dNTP (each), 200 nM each i7/i5 primer, ~1 U Phusion pol). PCR2 primers (i7/i5) are described in **Supp. Fig. 1**. In some cases (larger anticipated deletions with two Cas9 constructs), amplicon products were visualized by agarose gel electrophoresis with SybrSafe (Invitrogen) on a Bio-Rad SubCell Model 192 (Bio-Rad Laboratories, Hercules, CA).
***MiSeq deep sequencing***—10 μL amplicons from each PCR2 well were pooled and cleaned by SPRISelect (Beckman Coulter, Brea, CA) @1:1 ratio (100 μL pooled amplicons + 100 μL SPRISelect beads). The pooled library was diluted to within range of standards provided with the KAPA Library Quantification Kit for Illumina® Platforms (KK4844, KAPA Biosystems, Wilmington, MA). qPCR quantification was performed as specified (6 μL 2X premix + 4 μL DNA standard or library, standards in 3 dilutions in triplicate (1:1,000,000, 1:10,000,000, 1:100,000,000), yielding estimated working concentration of 143.62 nM). Sample Sheet was prepared by a pilot version of *SampleSheet.py*, using 96 lines of plate:barcode assignments to populate 9,216 sample:barcode [Data] relationships as detailed in **ExampleTestFiles (10.5281/zenodo.3406862).** The library was diluted to 4 nM and prepared for sequencing using the MiSeq Reagent Kit v2: PE, 2x150 bp (Illumina®, San Diego, CA), according to manufacturer instructions (10 pM with 10% PhiX DNA spike-in (PhiX Control v3, Illumina®)). MiSeq cluster density was 449 ± 11 k/mm$^2$, with 90% of reads passing filter to yield 8,065,046 reads (on average 875 reads/well).
***Genotypic analysis***—18,432 fastq files (representing R1 & R2 files for 9,216 demultiplexed barcode (sample)) were generated by MiSeq Controller Software onboard the MiSeq instrument. For genotype processing by *ImputedGenotypes,* fastq files were sorted to subdirectories by overarching sample ID (Cas9 treatment combination, *e.g.,* KE-1, KE-2, …KE-25) and submitted to the Jupyter notebook script in batches.
***TFBS analysis***—As for genotype processing, TFBS collation by *CollatedMotifs.py* was performed for fastq files in sample batches.
***Regulatory analysis (dex treatment, RNA isolation and RT-qPCR of ablated clones)***—Cells revived from liquid $N_2$ storage were grown to near-confluency in 10-cm dishes, trypsinized and counted by hemocytometer, and diluted to 1.5 x $10^5$ cells/mL. 3 mL diluted cells were transferred to each well of a 6-well tissue culture dish (Corning), and the entire plate was gently vortexed at speed setting 4-5 while loosely held *flat* on a 3-inch platform attachment of the Vortex Genie 2 (ThermoFisher), to evenly distribute cells in wells. Plates were transferred to a 37 °C incubator; 24 h later, media was aspirated and replaced with 2400 μL DMEM/5% FBS (charcoal/dextran-stripped, Omega Scientific, Tarzana, CA), and plates were returned to incubator for 3 h. During this time, dexamethasone (Sigma, St. Louis, MO) stock at 5 mM in ethanol (Decon Koptec, King of Prussia, PA) was diluted to 1:100 and (serially) to 1:10,000 in ethanol. At 3 h time point, 1 μL each dex stock (or ethanol for control) was added to 10 mL media/charcoal-stripped FBS. 600 μL of appropriate treatment stock was added to appropriate wells, mixed gently by rocking, and plates were returned to incubator. 4 h later, media was aspirated; cells were washed with 3 mL PBS, lysed in 350 μL RLT buffer (Qiagen, Hilden, Germany) with β-mercaptoethanol (Bio-Rad Laboratories), transferred to 1.5 mL Eppendorfs, flash-frozen in liquid $N_2$, and stored at -80 °C until processing.

For RNA isolation, cell lysates were transferred to QIAShredder (Qiagen) and centrifuged at full speed, 2 min. 300 µL 70% ethanol was added to flow-through and the volume was transferred to an RNeasy Mini Kit tube and centrifuged at full speed, 30 sec. Flow-through was discarded and 350 µL Buffer RW1 (wash buffer) was added to the column, then centrifuged full speed, 30 sec. Flow-through was discarded. For each sample, 10 µL RNase-free DNase I (Qiagen) stock solution was added to 70 µL Buffer RDD, mixed, added directly to the RNeasy column, and incubated on benchtop for 15 min. 350 µL Buffer RW1 was added, followed by centrifugation, then 500 µL RPE followed by centrifugation, then 500 µL RPE with centrifugation for 2 min. RNA was eluted into a 1.5 mL collection tube in 30 µL RNase-free $H_2O$, and concentration was determined by NanoDrop.

In preparation for qPCR, a small volume of RNA sample was set aside to use as a 'no RT' control. For reverse transcription, 1 µg RNA was used as template in 500 µL RNase-free tubes (Ambion) using iScript (1 µL) plus 4 µL 5X iScript reaction mix (Bio-Rad) and nuclease-free $H_2O$ to 20 µL reaction volume, with incubation using thermocycler program ISCRIPT (25 °C for 5 min, 42 °C for 30 min, 85 °C for 5 min, 4 °C indefinitely).

For qPCR, reactions were assembled on ice in MicroAmp Optical 384-Well Reaction Plates (Applied Biosystems, Life Technologies, Foster City, CA). cDNA from iScript reaction was diluted 4-fold in nuclease free $H_2O$, targeting 4 µL (50 ng) cDNA/reaction. Oligos were diluted by mixing fwd and rev primers (100 µM stocks) 1:1, with further 60.24-fold dilution in nuclease-free $H_2O$ to a working stock concentration of 0.83 M each oligo. 6 µL diluted primer mix volumes were added to appropriate wells of a 96-well qPCR plate using low-retention tips and multichannel pipet (targeting 250 nM final working concentration in 20 µL qPCR volume). 4 µL diluted cDNA was then added, followed by 10 µL SsoAdvanced Supermix (Bio-Rad). Plates were sealed with clear Microseal B adhesive seals (Bio-Rad), vortexed briefly, then centrifuged 5 min, 1500*g* in a table-top centrifuge. Cycling was performed using 95 °C, 30 sec, [95 °C, 5 sec, 57 °C, 30 sec], cycle 39x, with melt curve analysis in QuantStudio™ Real-Time PCR Software v1.3 on a QuantStudio 6 Flex 384-well instrument (Applied Biosystems). Three loci were used as internal reference controls (*RPL19, HMBS, GAPDH*) for geometric mean normalization[3]. Fold-change was expressed as ΔΔCT (dex/EtOH), with analysis performed *via* Python pandas data frames and plots generated by matplotlib. ΔCT was calculated as $CT_{experimental\ gene} - CT_{geometric\ mean\ [reference\ controls]}$. A549 clone identities for ΔΔ-26.65 kb mutants in Fig. 5 are 11-1 D06 (*clone #1*) and 11-5 G06 (*clone #2*).
